# Cell autonomous inflammation in VEXAS is mediated by cGAS-STING

**DOI:** 10.64898/2026.05.26.727520

**Authors:** Samuel J. Magaziner, Jason C. Collins, Brecca Miller, Patrick Zheng, Amy K. Wang, Jerome Hadjadj, Juan Carlos Baladrán, Maria Sirenko, Maya English, James Bertlin, Rebecca Murray, Peter H. Whitney, Tania J. González-Robles, Deborah Rivera, Yan Wang, Duy T. Tran, Zulfeqhar A Syed, Valentina Baena, Timothee Lionnet, Kelly V. Ruggles, Iannis Aifantis, Dan A. Landau, Achim Werner, David B. Beck

## Abstract

VEXAS (vacuoles, E1 enzyme, X-linked, autoinflammatory, somatic) is a severe adult-onset inflammatory disease caused by somatic mutations that reduce cytoplasmic activity of UBA1, the primary initiating enzyme for ubiquitylation. How this hypomorphic state drives cell-intrinsic immune activation in mature myeloid cells is unknown. Using unbiased multi-omic, biochemical, and cell biological analyses of model systems and patient-derived cells, we show that loss of cytoplasmic UBA1 activity convergently disrupts endoplasmic reticulum– associated degradation (ERAD) and mitochondrial homeostasis. ERAD failure arises from preferential under-charging of ERAD E2 enzymes, explaining hallmark VEXAS features, including ER-derived vacuoles and unfolded protein response activation, and promotes accumulation of the ERAD substrate STING. Simultaneously, mitochondrial dysfunction drives cytosolic leakage of mitochondrial DNA, inducing cGAS-dependent STING signaling and inflammatory cytokine production. STING inhibition or reversal of mitochondrial DNA leakage resolves multi-cytokine inflammation in VEXAS models and patient myeloid cells, establishing the cGAS–STING pathway as a therapeutically actionable vulnerability.

---

Systemic autoinflammatory diseases are monogenic disorders that cause constitutive activation of the innate immune system (*1*). VEXAS (vacuoles, E1 enzyme, X-linked, autoinflammatory somatic) is a severe, late-onset autoinflammatory condition that affects 1:4000 males over 50 years of age (*2, 3*). VEXAS is caused by somatic hypomorphic mutations in UBA1, which encodes the E1 enzyme responsible for initiating the majority of ubiquitylation. The disease-causing mutations in VEXAS are acquired and restricted to the hematopoietic system, most prominently in the myeloid lineage. Patients with VEXAS suffer from severe inflammation, which can affect various organs, including the skin, lungs, and joints, along with hematologic conditions such as pancytopenia and myelodysplastic syndrome (*4, 5*). VEXAS is associated with significantly increased mortality, and highly effective treatments are currently lacking (*6*).

UBA1 is ubiquitously expressed and functions as the initial enzyme in the ubiquitin-proteasome system, charging most E2 enzymes in the cell (*7*). As a result, UBA1 is pivotal for processes such as the cell cycle, proliferation, growth, and dynamic responses to stimuli. Ubiquitylation is also essential for maintaining a balanced proteome by facilitating the degradation of unnecessary or misfolded proteins through mechanisms such as autophagy-mediated degradation, the endoplasmic reticulum-associated degradation (ERAD) pathway, and the unfolded protein response (UPR) (*8-10*).

Most VEXAS-causing mutations occur at p.Met41 and result in the loss of cytoplasmic UBA1 activity through decreased translation of the cytoplasmic isoform or are characterized by impaired E2 enzyme charging (*11, 12*). Bulk and single cell RNA sequencing from VEXAS patient cells have revealed pronounced inflammation, particularly within the myeloid lineage (*13-15*). This inflammation is characterized by the activation of the type I interferon pathway and other proinflammatory cytokines (e.g., IL-6 and TNFα). Furthermore, these correlative studies have identified concomitant activation of the UPR, which has been implicated as a downstream mechanism of inflammation in disorders related to ubiquitylation, including proteasomopathies (*13, 16-18*). Recent work has also elucidated the role of aberrant TN-Fα-dependent cell death in non-cell autonomous inflammation and propagating systemic disease pathologies (*19, 20*). However, because TNF blockade is not effective in treating VEXAS patients (*6*) and existing mutant models do not show spontaneous inflammation or cell death (*14, 21, 22*) additional upstream mechanisms must initiate and maintain inflammation. In this study, we utilized multiple complementary cellular models and patient-derived samples to define how loss of cytoplasmic UBA1 activity drives cell autonomous inflammation in mutant myeloid lineages. Our data identify the central role of ERAD failure and mitochondrial dysfunction as key upstream events that converge on cGAS-STING activation, thereby initiating and sustaining VEXAS-associated inflammation.

## RESULTS

### UBA1 Pathogenic Variants Lead to Inflammation, ERAD Dysregulation, UPR activation, and Vesicle Disruption

To dissect the molecular events driving inflammation in VEXAS myeloid cells, prior to the pleiotropic effects and adaptation after loss of ubiquitylation and without the complexities of patient-derived tissues, we developed an inducible in vitro model (***Fig. S1A***). This system uses human THP1 monocytes/ macrophages engineered to undergo CRISPR-Cas9-mediated deletion of endogenous UBA1 upon induction with doxycycline (dox), complemented with either UBA1^WT^ or UBA1^M41V^ (***Fig. 1A***). Our VEXAS model, THP1 UBA1^M41V,^ successfully recapitulated key disease features, including the cytoplasmic isoform switch from active UBA1b to catalytically impaired UBA1c (***Fig. 1B, Fig. S1B***), reduced global ubiquitylation (***Fig. 1B, Fig. S1B***), UPR activation (***Fig. 1B, Fig. S1C***), vacuole formation (***Fig. S1D,E***), and inflammation of VEXAS hallmark pathways (e.g., Type-I interferon, IL-6, and TNFα) that was present in the monocyte and more pronounced in the macrophage state (***Fig. 1B-D, S1C***) at day 9 post-induction. Notably, no spontaneous apoptosis, pyroptosis, or necroptosis was observed in THP1 UBA1^M41V^ monocytes or macrophages despite robust inflammation (***Fig. 1B***), suggesting novel cell-intrinsic pathways are required to initiate inflammation in VEXAS myeloid cells.

**Fig. 1.**
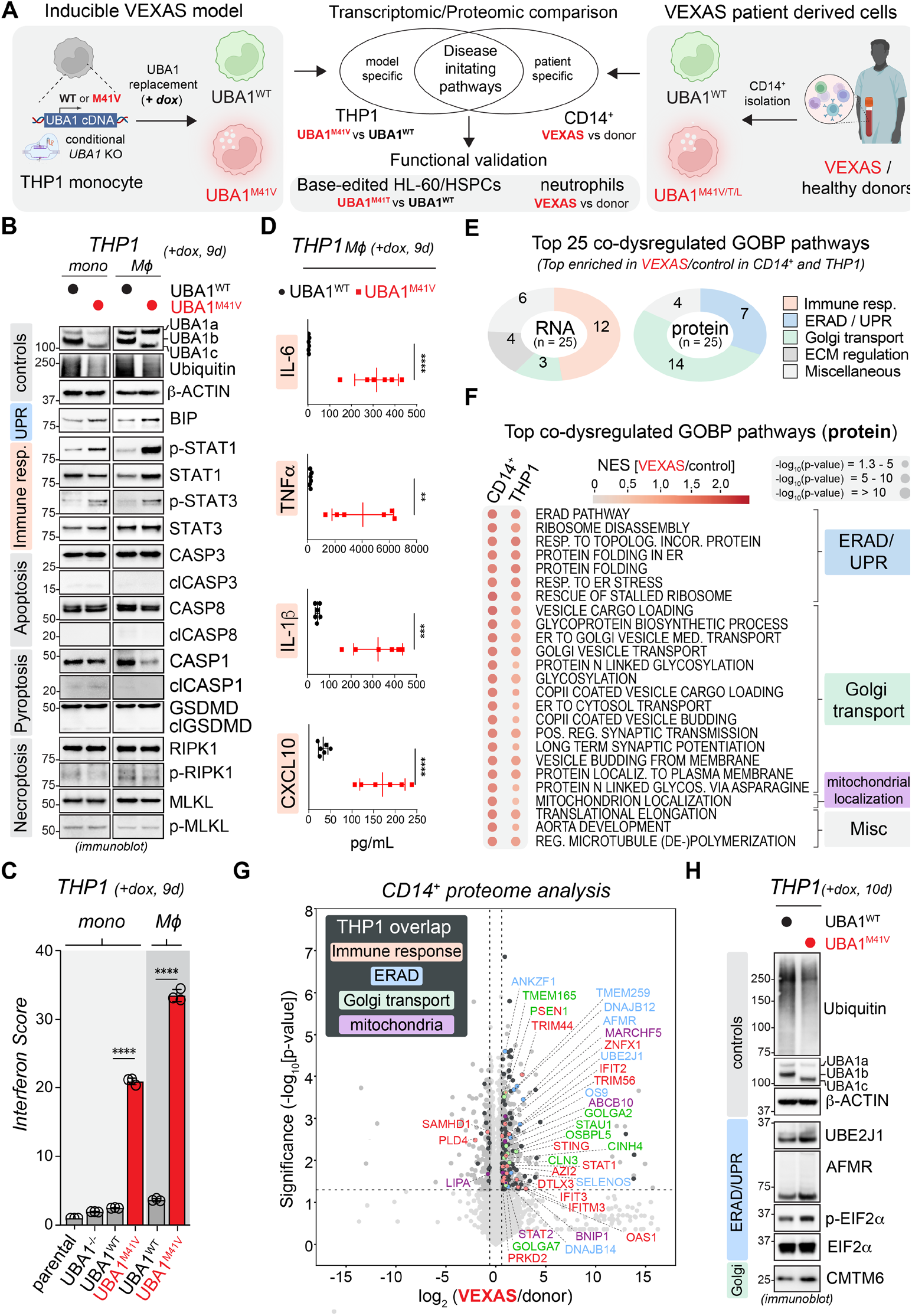
ERAD dysfunction, Golgi disruption, and UPR activation are early consequences following loss of cytoplasmic UBA1 function in VEXAS. (**A**) Schematic workflow of proteomic and transcriptomic analyses across both model THP1 monocytes (*left*), and patient derived control and VEXAS monocytes (*right*) to identify disease initiating pathways. (**B**) Loss of cytoplasmic UBA1 function in THP1 model cells leads to ERAD dysregulation, UPR activation, and inflammatory signaling without evidence for cell death. UBA1^WT^ and UBA1^M41V^ THP1 cells were treated with dox for 9 d and subjected to immunoblotting using the indicated antibodies. Results are representative of n ≥ 3 biological replicates. (**C**) THP1 VEXAS model cells exhibit type I interferon signatures, confirming the results of the transcriptomic analyses. Parental, UBA1-/-, UBA1^WT^ and UBA1^M41V/L^ THP1 monocytes or macrophages (M ϕ) were treated with dox for 9 d and analyzed by qPCR for an Interferon Score by averaging the fold change of 6-hallmark ISGs (IFI44L, IFI27, IFIT1, ISG15, RSAD2, and SIGLEC1). n ≥ 3 biological replicates, error bars = s.e.m., **** = p < 0.0001, student’s t-test. (**D**) Loss of cytoplasmic UBA1 function in THP1 model cells leads to multi-cytokine inflammation. UBA1^WT^ and UBA1^M41V^ THP1 macrophages (Mϕ) were treated with dox for 9 d and ELISA for IL-6, TNFα, IL-1b, and CXCL-10. n = 3 biological replicates, ** = p < 0.01, error bar = s.e.m, student’s t-test. (**E**) Proteomic and transcriptomic analyses reveal overlapping and unique altered pathways in VEXAS and nominate ERAD and Golgi transport dysregulation as initial signals in disease pathogenesis. Pie charts depict the top 25 gene ontology biological pathways (GOBPs) significantly enriched in both patient-derived VEXAS over control monocytes (CD14^+^) and the VEXAS model cells (THP1). While in the transcriptomic comparison (*upper chart*) the majority of the top 25 enriched GOBPs are related to the immune response, the proteomic comparison (*lower chart*) reveals predominantly GOBPs associated with ERAD and Golgi transport. (**F**) Dot plots show gene set enrichment scores for the top 25 dysregulated GOBPs as identified by proteomic comparison of patient-derived control and VEXAS monocytes (CD14^+^). For each CD14^+^ top dysregulated GOBP, the mean normalized enrichment score (NES) and the Benjamini-Hochberg-adjusted p-values are shown for the two indicated comparisons (CD14^+^ protein, THP1 protein). n=3 biological replicates for each condition. (**G**) The proteomes of patient derived CD14^+^ and THP1 model VEXAS cells are highly similarly dysregulated and exhibit altered immune responses and ERAD dysregulation. The volcano plot depicts differentially expressed proteins (DEPs) of healthy donor and VEXAS CD14+ monocytes. DEPs colored in black are significantly regulated in both VEXAS CD14+ and THP1 model cells. Within this overlap, DEPs related to immune responses, ERAD and Golgi transport are highlighted. (**H**) Immunoblot analysis of THP1 model cells with indicated antibodies reveals that loss of cytoplasmic ubiquitylation causes ERAD and Golgi dysregulation.

To identify such pathways in an unbiased manner, we conducted transcriptomic, proteomic, and ubiquitylated protein enrichment analyses in both THP1 UBA1^M41V^ and VEXAS patient-derived monocytes (***Fig. S2-3, Tables S1-5***). Compared to their control counterparts, both patient-derived and model cells exhibited highly correlated dysregulated pathways, further confirming that THP1 UBA1^M41V^ cells faithfully replicate key features of VEXAS (***Fig. S3A***). Focusing on pathways enriched in VEXAS CD14^+^ monocytes, we identified shared activation of inflammatory signaling and the unfolded protein response (UPR) in both the THP1 model and patient cells (***Fig. 1E,F, Fig. S3B, Table S1***). These findings are consistent with prior studies using patient samples (*13, 17*). Notably, when we analyzed proteomic datasets, we identified novel dysregulated pathways, including ERAD (endoplasmic reticulum-associated degradation) and vesicle-mediated transport, which had not been previously implicated in VEXAS pathogenesis or prioritized in transcriptomic studies (***Fig. 1E,F, Tables S2,3***).

Given the role of UBA1 in post-translational regulation and the stronger correlation between model and patient datasets at the proteomic level (***Fig. S3A***), we next focused on proteomic pathways enriched in both patient and THP1 UBA1^M41V^ monocytes (***Fig. 1E-G, Tables S2,3***). While inflammatory and UPR pathways were identified as expected in transcriptomic analyses, ERAD, Golgi transport and mitochondrial regulation emerged amongst the most enriched pathways, with many critical proteins showing significant enrichment compared to controls (***Fig. 1G***). Immunoblotting and immunofluorescence of the THP1 UBA1^M41V^ model validated these findings, confirming ER stress and dysregulated ERAD and Golgi vesicle transport (***Fig. 1H, Fig. S1G***). Additionally, bulk and scRNA seq data from patient monocytes corroborated deregulation of ERAD pathways in VEXAS (***Fig. S4A,B, Tables S4,5***). We made similar observations for hematopoietic stem and progenitor cells (HSPCs) (***Fig. S4B***). Using a base editor approach we constructed an HSPC model of VEXAS, in which the UBA1^M41T^ mutation is introduced into CD34^+^ donor HSPCs (*14,17,23*) and also found ERAD as the top dysregulated GOBP pathway in proteomic analyses along with dysregulated pathways suggesting possible mitochondrial dysfunction, such as reactive oxygen species metabolism and the electron transport chain (***Fig. S4C-E***)

Taken together, through unbiased approaches, our inducible model of VEXAS successfully reproduces dysregulated pathways observed in patient cells and identifies ERAD dysfunction and mitochondrial dysregulation as early disease signals in VEXAS pathogenesis.

### ERAD Failure Precedes Inflammation and Results from a Preferential Loss of Ubiquitin Charging to ER-tethered E2s

To further delineate the stepwise pathways causing inflammation in VEXAS, we monitored the proteomes of our THP1 model at different time periods after replacement of endogenous UBA1 with either UBA1^WT^ or UBA1^M41V^ (***Fig. S2A,B***). Notably, ERAD- and UPR-related pathways and proteins were significantly enriched in VEXAS model cells before the onset of inflammation (***Fig. 2A***), suggesting that loss of ERAD function with subsequent activation of the UPR precedes VEXAS inflammation. To test whether ERAD was indeed impaired in VEXAS cells, we utilized a fluorescent reporter (CD3δ -GFP) that is degraded through the ERAD pathway (***Fig. S5A,B***). In VEXAS THP1 cells, reporter levels were elevated under basal conditions and were not as efficiently degraded upon inhibition of protein translation using cycloheximide, demonstrating impaired ERAD function (***Fig. 2B***). We made similar observations for 32D UBA1^M41L^ cells, a recently described constitutive murine cell model of VEXAS (***Fig. S5C***) (*22*). Additionally, ERAD inhibition using EerI caused selective lethality in VEXAS cell lines, similar to treatment with the UBA1 inhibitor TAK-243, further indicating impairment of the pathway in VEXAS (***Fig. S6A,B***).

**Fig. 2.**
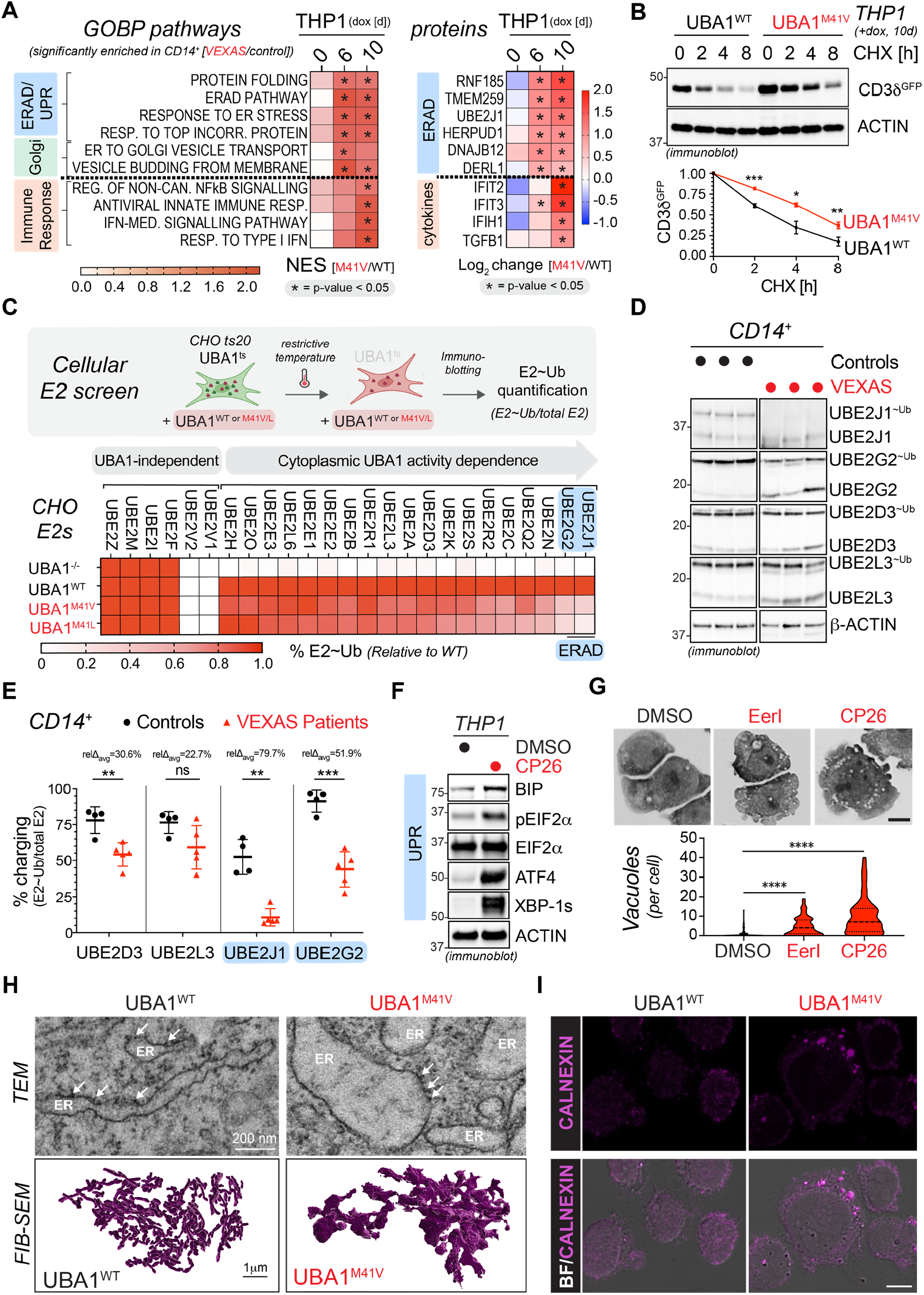
ERAD failure precedes VEXAS inflammation, results from an undercharging of ER-tethered E2s, and causes UPR activation and ER-derived vacuoles. (**A**) ERAD alterations precede inflammation in THP1 VEXAS models, shown via heatmap representation of gene ontology biologic processes (GOBPs) across indicated time periods after doxycycline induction. Representative GOBPs and proteins were selected from those significantly dysregulated in patient CD14^+^ cell proteomics. Color scale indicates the mean normalized enrichment score (NES) of UBA1^M41V^ versus UBA1^WT^ THP1 monocytes with asterisks representing Benjamini-Hochberg-adjusted p-value of -log_10_(p-value) ≥ 1.3. (**B**) ERAD is impaired in VEXAS model cells. UBA1^WT^ and UBA1^M41V^ THP1 macrophages expressing the ERAD reporter, CD3d-GFP, were induced with doxycycline for 10 days, treated with cycloheximide (CHX) for indicated time periods, and subjected to immunoblotting with indicated antibodies. CD3d-GFP quantifications were normalized to Actin. N ≥ 3 biologic replicates, error bars = s.e.m., * = p < 0.05, ** = p < 0.01, *** = p < 0.001, students t-test. (**C**) Loss of cytoplasmic UBA1 activity leads to a preferential loss of ubiquitin charging to the ERAD-associated E2 enzymes UBE2J1 and UBE2G2. Schematic depicts the workflow of the biochemical screen to identify E2 charging defects associated with VEXAS mutations. Chinese hamster ovary (CHO) cells with a temperature sensitive Uba1 allele (ts20) and complemented with human UBA1^WT^, UBA1^M41V^, or UBA1^M41L^ were incubated at the restrictive temperature and subjected to immunoblotting using antibodies against the 28 detectable E2s in CHO cells. Heatmap depicts the quantifications of the ubiquitin charging status of each E2 enzyme based on immunoblot signal of ubiquitin-charged over total E2, normalized to UBA1^WT^. Some E2 enzymes are known to be charged independent of UBA1 (UBA1-independent), while the majority of other E2s show varying degrees of sensitivity to loss of cytoplasmic UBA1 activity (UBE2Ds = UBE2D1/2/3/4). (**D**) Confirmation of the specific E2 charging defect of ERAD-associated UBE2J1 and UBE2G2 in patient cells. CD14^+^ monocytes from control or VEXAS patients were isolated and subjected to immunoblotting with indicated antibodies. (**E**) Quantification of the ubiquitin charging status for each E2 (E2∼Ub/ total E2) depicted in panel e. n ≥ 4 biological replicates, error bar = s.e.m, ** = p < 0.01, *** = p < 0.001, student’s t-test. (**F**) ERAD inhibition leads to activation of the unfolded protein response (UPR), as shown by immunoblotting for UPR and integrated stress response (ISR) components upon treatment of parental THP1 cells with 5 μM ERAD inhibitor CP26. (**G**) ERAD inhibition leads to cytoplasmic vacuole formation, as assessed via Wright-Giemsa staining after treatment of THP1 cells with the ERAD inhibitors EerI (2.5 μM) and CP26 (5 μM). Vacuoles were quantified via a machine learning algorithm (see methods). n = 3 biologic replicates, **** = p < 0.001, student’s t-test. (**H**) Loss of cytoplasmic UBA1 activity in THP1 cells causes abnormal ER morphology with dilated, vacuole-sized, and proteinaceous cisterna. UBA1^WT^ and UBA1^M41V^ THP1 cells were treated with doxycycline for 10 days and analyzed by transmission electron microscopy (TEM, upper panel) or focused ion beam scanning electron microscopy (FIB-SEM, lower panel). For TEM, Representative images of 2 biologic replicates are shown (for quantifications, see Fig. S10). For FIB-SEM, 3D reconstructions depict a representative UBA1^WT^ and UBA1^M41V^ THP1 cell, demonstrating dilated ER coursing through the UBA1^M41V^ THP1 cell. (**I**) VEXAS-associated vacuoles originate from ER, as demonstrated by overlay of brightfield (BF) and confocal anti-calnexin immunofluorescence images of UBA1^WT^ and UBA1^M41V^ THP1 cells treated with doxycycline for 10 days in both single plane and maximal intensity projection (max IP). Scale bar = 10 μm

Having established that ERAD is defective in VEXAS, we investigated why the loss of UBA1 impairs ERAD. Our previous findings revealed that VEXAS is caused by cytoplasmic defects, with all disease-causing mutations most prominently impairing E2 activation (*11*). Based on these insights, we performed an unbiased biochemical screen using the ts20 CHO cell line, which is hemizygous for a temperature-sensitive Uba1 allele (*24, 25*). At restrictive temperatures, Uba1 loss results in decreased ubiquitylation, E2 charging, vacuole formation and cell death (***Fig. 2C***)(*24*). Using this system, we complemented ts20 cells with wild-type human UBA1 or VEXAS-associated mutations and assessed their effects on E2 charging.

As expected, loss of Uba1 caused broad E2 charging defects for all UBA1-dependent E2 enzymes, which were rescued by wild-type UBA1 expression (***Fig. 2C***). In contrast, VEXAS-associated UBA1^M41V^ or UBA1^M41L^ mutations only re-established ubiquitin charging of a subset of E2s and with varying efficiency, revealing UBE2J1 and UBE2G2 as most sensitive to loss of cytoplasmic UBA1 function (***Fig. 2C, Fig. S7A***). Intriguingly, these E2 enzymes are key components of the ERAD pathway, forming part of the multi-protein complex required for the retrotranslocation and degradation of misfolded ER proteins (*26*). Consistent with their involvement in disease pathogenesis, we confirmed preferential loss of ubiquitin charging of these enzymes in four distinct systems, including THP1 and 32D VEXAS models, patient-derived cells, and UBA1^M41T^ base-edited HSPCs (***Fig. 2D,E, Fig. S7B-E***). Interestingly, when combined with the UBA1 inhibitor TAK-243, knockdown of these E2s induced a synthetic lethality akin to TAK-243 treatment of UBA1^WT^ versus UBA1^M41V^ THP1 cells (***Fig. S6C***). This preferential E2 charging defect detected in vivo was absent in in vitro assays, in which recombinant UBA1c was similarly defective in transferring ubiquitin to ERAD E2s and 20 other tested E2 enzymes (***Fig. S8A-F***). This indicates that cellular signals, E2 concentrations, or organelle-specific contexts contribute to the preferential under-charging of UBE2J1 and UBE2G2 *in vivo*.

Collectively, these findings demonstrate that ERAD impairment in VEXAS arises from a preferential E2 charging defect to ER-tethered enzymes in cells, providing a mechanistic link to disease pathogenesis.

### ERAD Impairment is Required for VEXAS-associated Phenotypes

To investigate whether ERAD failure is required for inducing cellular VEXAS phenotypes, we performed targeted inhibition of the ERAD pathway in THP1 cells. First, we used the chemical inhibitors CP26 or EerI, which block retrotranslocation and result in accumulation of ERAD substrates in the ER(*27, 28*). ERAD-inhibited cells exhibited active UPR and integrated stress response (ISR), characterized by increased phosphorylation of eIF2α, XBP1 splicing, elevated ATF6 and BIP levels, as well as cytoplasmic vacuoles—all hallmarks of VEXAS cells (***Fig. 2F,G***). Similar findings were made with siRNA or shRNA-mediated depletion of UBE2J1/G2 (***Fig. S9A-D***).

Loss of ERAD function has previously been shown to cause accumulation of unfolded proteins in and extensive swelling of the ER, suggesting that VEXAS-associated vacuoles in monocytes could be ER-derived (*29*). Indeed, transmission electron microscopy (TEM) and focused ion beam scanning electron microscopy (FIB-SEM) analysis of our VEXAS THP1 model revealed that, while ER in UBA1^WT^ cells appeared as long and thin interconnected cisternae, ER in many UBA1^M41V^ cells appeared swollen and fragmented, with vacuoles emerging as membrane-bound, ribosome-associated structures filled with proteinaceous electron-dense material (***Fig. 2H, Fig. S10,11, Movie S1***). We could corroborate these findings by immunofluorescence microscopy experiments, in which the ER-resident protein CALNEXIN labeled structures that appeared as vacuoles in brightfield images (***Fig. 2I***). Therefore, many VEXAS-associated vacuoles originate from the ER after ERAD failure.

Despite the above overlapping phenotypes induced by ERAD inhibition alone, qPCR, immunoblotting, or ELISA analyses failed to detect VEXAS-associated inflammation (***Fig. S12A-D***). We hypothesized that this absence of inflammatory signaling might be due to another cytoplasmic ubiquitylation event, which is disrupted in VEXAS but not captured by ERAD inhibition alone. To test this, we treated cells with low doses of the UBA1 inhibitor TAK-243 following ERAD inhibition (***Fig. S12E***). Indeed, some aspects of VEXAS-associated inflammation could only be elicited when both UBA1 and ERAD inhibition treatments were combined (***Fig. S12F-H***). These results suggest that ERAD failure is necessary but not sufficient for initiating inflammation in VEXAS (***Fig. S12I***).

The UPR and ISR have previously been linked to a variety of inflammatory diseases including proteasome-related disorders, with PKR inhibition specifically reversing inflammation in PRAAS (18, *30-32*). Thus, we hypothesized that UPR/ISR activation may be the mechanism by which ERAD failure contributes to inflammatory signaling in VEXAS. We investigated whether inflammation could be reversed by inhibition of different branches of the UPR and ISR (PERK, IRE1α, PKR, GCN2) in THP1 model cells (***Fig. S13A***). Surprisingly, IRE1α or PKR inhibition was able to resolve Type I interferon mediated inflammation as measured by qPCR (***Fig. S13B***); however, both caused paradoxical increases in TNFα and other cytokines when measured by ELISA (***Fig. S13C,D***). This aligns with recent work which has shown a reliance of UPR and PERK signaling for VEXAS mutant HSPC viability (*17*). Other targeted UPR/ISR inhibitors conferred modest resolution with equally paradoxical activation of different inflammatory pathways (***Fig. S13***). Altogether, these findings indicate that the UPR and ISR only partially contribute or modulate inflammatory signaling in VEXAS and suggest that additional mechanisms following ERAD failure must be responsible for innate immune activation.

### ERAD failure causes STING accumulation and signaling

To identify additional factors contributing to inflammation in VEXAS, we hypothesized that ERAD failure, beyond causing increased unfolded proteins and UPR activation, could lead to the accumulation of specific proteins that amplify inflammatory signaling. Proteomic analysis of the THP1 model revealed a progressive accumulation of various ERAD substrates (***Fig. S14A, Table S6***). Amongst these, STING (Stimulator of interferon genes), a pathogen sensor, was the only identified protein associated with innate immunity and exhibited increased levels in the THP1 and 32D models of VEXAS and VEXAS patient-derived cells (***Fig. 3A, Fig. S14B-D***). Importantly, STING was not identified as an upregulated gene in our transcriptomic data sets (***Tables S1-5***), suggesting its accumulation is not secondary to interferon stimulation. Consistent with STING accumulating post-translationally, mainly through disrupted ERAD in VEXAS cells, only inhibition of ERAD, rather than inhibition of other reported degradation pathways (autophagy (*33*) and ESCRT-dependent (*34*)) led to significant increases of STING levels in THP1 cells similar to those observed in VEXAS model and patient cells (***Fig. 3B***). STING increases also inversely associated with a decrease in ubiquitylated STING and closely paralleled inflammation in our THP1 time course (***Fig. 2A, Fig. S14E,F***). In addition, previous studies have shown that STING accumulation occurs during ERAD failure, priming cells for STING-mediated inflammation when a secondary trigger is introduced (*35*). Similarly, UBA1 inhibition has been reported to enhance STING-mediated inflammation (*34*).

**Fig. 3.**
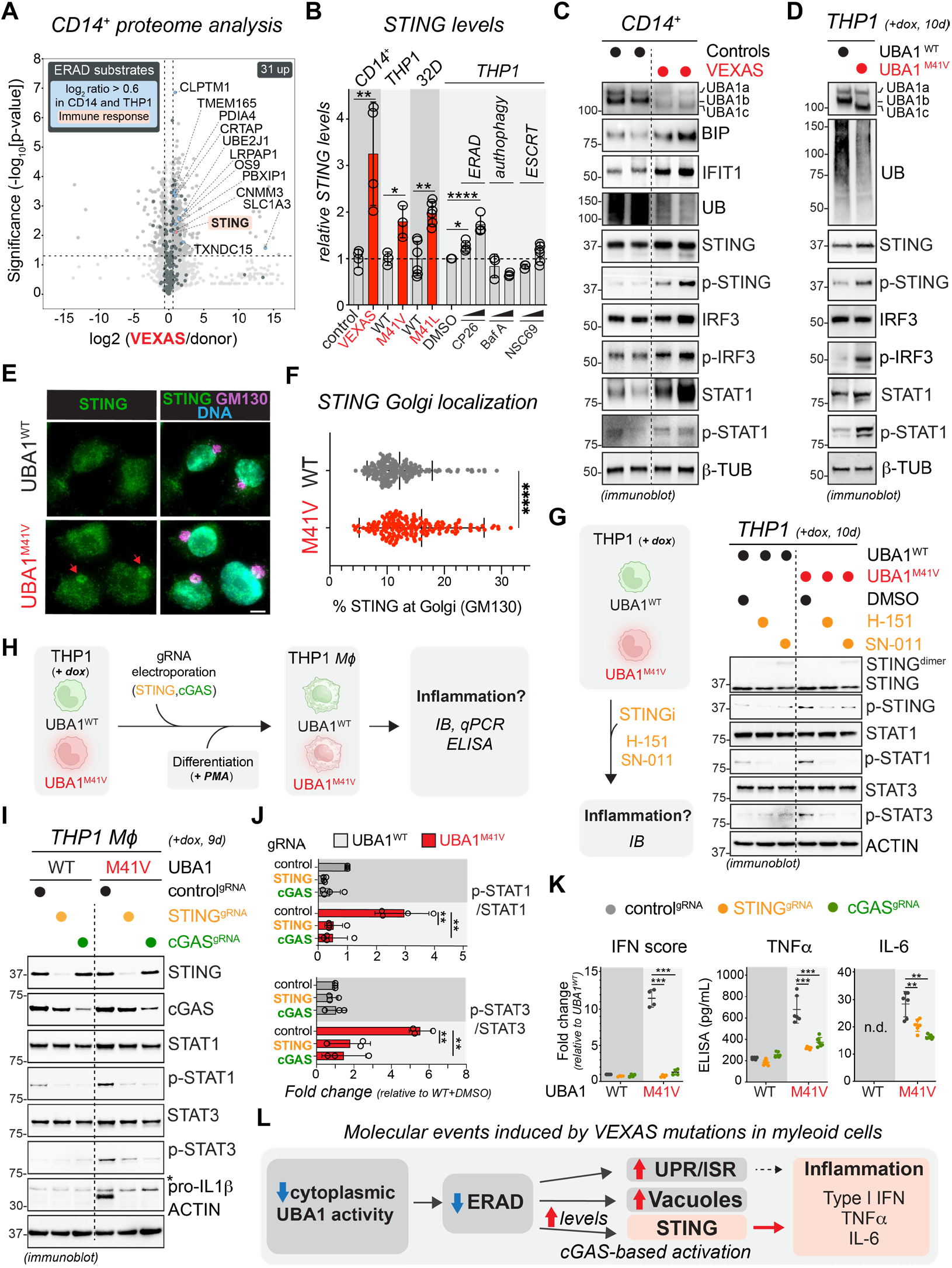
ERAD failure induces increases in STING, which drives VEXAS-associated inflammation in a cGAS-dependent manner. (**A**) STING is the sole immune response-related ERAD/ HRD1 substrate that accumulates in VEXAS CD14^+^ cells. The volcano plot depicts differentially expressed proteins (DEPs) of healthy donor and VEXAS CD14^+^ monocytes. DEPs colored in black highlight previously identified HRD1 substrates and DEPs in blue highlight HRD1 substrates significantly upregulated in VEXAS CD14^+^ cells. (**B**) Innate immune adaptor STING accumulates in VEXAS-patient derived CD14^+^ cells, UBA1^M41V^ THP1 monocytes, UBA1^M41L^ 32D cells, and upon ERAD inhibition (CP26, 1 μM or 2 μM for 24h), but not upon autophagy inhibition (BafA, 1 μM or 10 μM for 24h) or ESCRT-mediated degradation pathway inhibition (NSC69, 3 μM and 4 μM for 24h) in THP1 parental cells, as quantified from anti-STING immunoblots. n ≥ 3 biologic replicates as indicated, error bar = s.e.m, * = p < 0.05, ** = p < 0.01, student’s t-test. (**C**) The STING pathway is active in CD14^+^ cells from VEXAS patients, as shown via immunoblotting of cell lysates with indicated antibodies. **(D**) The STING pathway is active in UBA1^M41V^ THP1 monocytes as shown via immunoblotting of cell lysates with indicated antibodies. (E) STING is activated in VEXAS model cells, as evidenced by increased STING trafficking to the Golgi in UBA1^M41V^ THP1 monocytes observed by anti-STING and anti-GM130 immunofluorescence. Scale bar = 10 μm. (**F**) Quantification of the percentage of Golgi-localized STING of the experiment shown in panel e. n ≥ 150 cells across 10 fields, error bar = s.e.m, **** = p < 0.0001, student’s t-test. (**G**) STING inhibition reverses VEXAS-associated inflammatory signaling in THP1 model cells. UBA1^WT^ and UBA1^M41V^ THP1 macrophages (Mϕ) were treated with STING inhibitors H-151 (1 μM) and SN-011 (10 μM) for 24h and subjected to immunoblot analysis using indicated antibodies. (**H**) Schematic representation of experimental design for STING or cGAS knockout (KO) and assessment of inflammation. (**I**) STING or cGAS KO inhibit VEXAS-associated inflammatory signaling in THP1 model cells. UBA1^WT^ and UBA1^M41V^ THP1 macrophages (Mϕ) were electroporated with indicated gRNAs and subjected to immunoblot analysis using indicated antibodies. (**J**) Graphs depict quantifications of p-STAT1/STAT1 ratio normalized to control gRNA-electroporated UBA1^WT^ THP1 macrophages. n = 3 biological replicates, error bar = s.e.m, ** = p < 0.01, student’s t-test. (**K**) STING and cGAS KO inhibit VEXAS-associated inflammatory signaling in THP1 model cells. Same experiments as in panel I, but cell lysates were either subjected to qPCR analysis to determine mRNA levels of different type I interferon stimulated genes (*left graph*, IFN score), or cell supernatants were subjected to ELISA to measure secreted protein concentrations of TNFα (*middle graph*) or IL-6 (*right graph*). (**L**) Scheme depicting molecular events triggered by canonical VEXAS mutations in myeloid cells. Loss of cytoplasmic UBA1 activity causes ERAD impairment, which induces UPR/ISR signaling, vacuoles, and accumulation of the innate immune adaptor STING, which is activated in a cGAS-dependent manner.

In support of the role of STING in VEXAS-related inflammation, we observed aberrant activation of STING in patient and THP1 cells, as indicated by phosphorylation at Ser366 and downstream signaling activation of IRF3 (***Fig. 3C,D***) and increased STING localization to the Golgi in VEXAS model cells (***Fig. 3E,F***). To investigate the role of STING in mediating inflammation in VEXAS, we treated the THP1 model with two small molecule inhibitors of STING, H-151 or SN-011, and monitored inflammatory signaling (***Fig. 3G***). In contrast to blocking UPR and ISR components (***Fig. S13***), pharmacological inhibition of STING reversed most axes of inflammation in both monocytes and macrophages (Type I interferon, IL-6, TNFα, IL-1β) (***Fig. 3G, Fig. S15A***).

To understand the mechanisms that cause abnormal STING activation in VEXAS myeloid cells, we performed genetic knockout (KO) of STING and its upstream activator and dsDNA sensor cGAS (***Fig. 3H***). As expected from our inhibitor studies (***Fig. 3G, Fig. S15A***) STING KO suppressed inflammatory signaling and cytokine release in THP1 macrophages (***Fig. 3I-K***). Importantly, cGAS KO similarly reversed the multi-cytokine inflammatory phenotype, indicating that activation of STING in VEXAS requires a dsDNA ligand.

Together, these data demonstrate that STING accumulates as a consequence of ERAD failure in VEXAS myeloid cells and undergoes cGAS-dependent activation, thereby driving the inflammatory phenotype (***Fig. 3L***).

### cGAS-mediated sensing of mtDNA drives STING-dependent inflammation

Multiple sources of self-DNA, including DNA from engulfed dead cells, genomic DNA from ruptured micronuclei, or mitochondrial DNA (mtDNA) released into the cytoplasm, have been shown to cause inappropriate cGAS-STING activation across disease contexts (*36*). Chronic ER stress, such as those observed in our VEXAS models, has been extensively shown to drive calcium dysregulation and lead to non-lethal mitochondrial dysfunction (*37*). Indeed, we observed increased levels of the ER-resident calcium pump SERCA2 (***Fig. S15B***) and a marked elevation in cytosolic calcium levels in UBA1^M41V^ THP1 cells using the Indo-1am intracellular calcium stain (***Fig. S15C***).

We did not observe an increase in micronuclei (***Fig. S16A-D***), activation of LINE-1 retrotransposon elements (***Fig. S16E***) or appreciable spontaneous cell death (***Fig. 1B***). Instead, and corroborating our previous unbiased observations (***Fig 1F, Fig S4E***), immunofluorescence, TEM, and FIB-SEM analyses of the THP1 model revealed that UBA1^M41V^ cells contained fewer mitochondria with morphological abnormalities including dilated and vacuolized cristae, suggesting mitochondrial dysfunction and damage (***Fig. 4A-C, Fig. S16A,B, S17A,B, Movie S2***). Consistent with this, cellular fractionation and qPCR analysis of dsDNA content identified an increase in the proportion of cytosolic mtDNA as compared to control cells (***Fig. 4D***). A TMRE-mediated mitochondrial potential assay showed no loss in membrane potential in mutant cells, suggesting that mitochondrial depolarization is not the primary driver of mtDNA release (***Fig. S17C***).

**Fig. 4.**
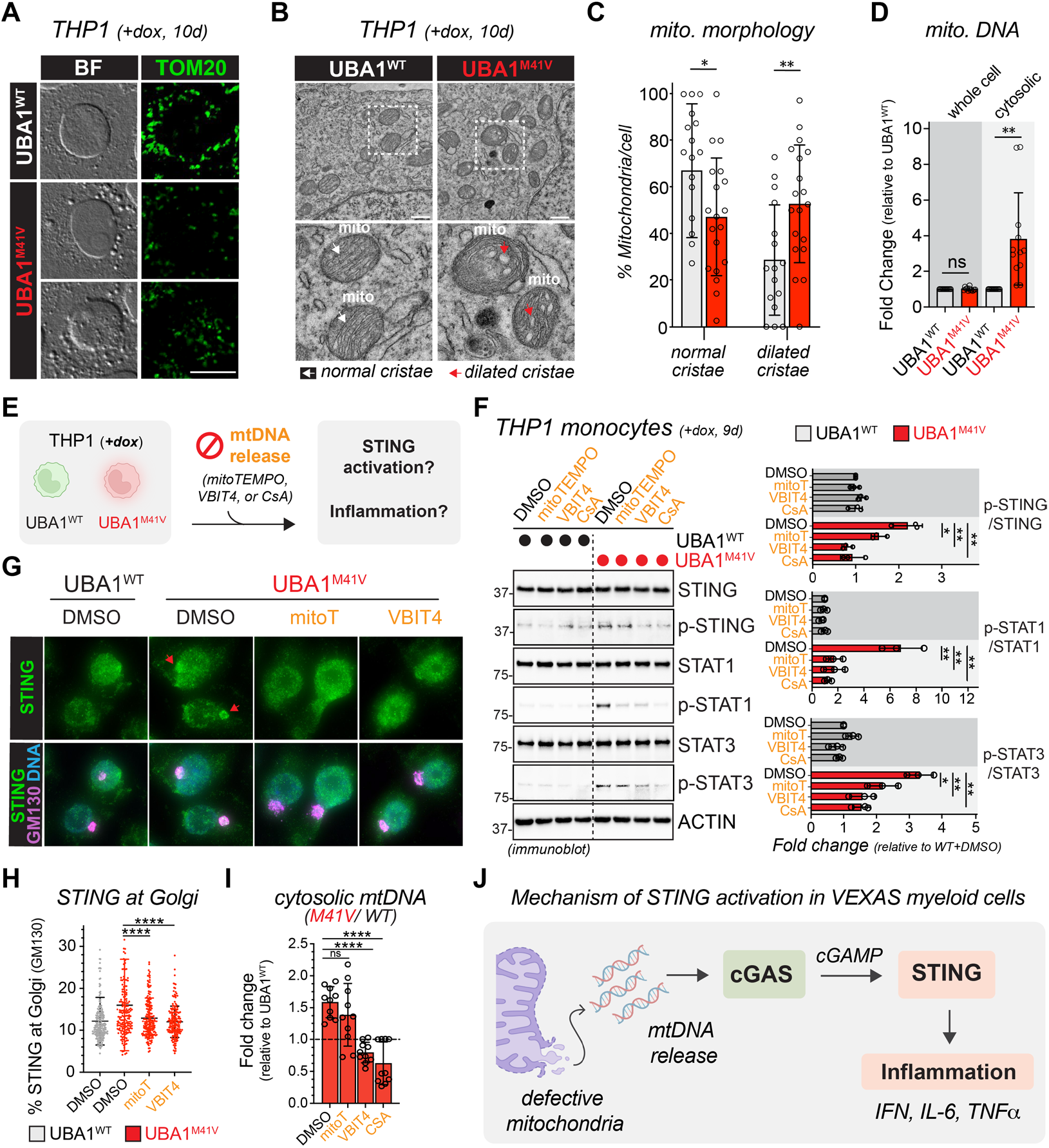
cGAS-STING-mediated inflammation in VEXAS cells requires mtDNA leaking into the cytosol. (**A**) Vacuole-containing VEXAS THP1 cells contain less mitochondria, as demonstrated by brightfield (BF) and single-plane confocal anti-TOM20 immunofluorescence images of UBA1^WT^ and UBA1^M41V^ THP1 monocytes treated with doxycycline for 10 d. Representative images of 2 biological replicates are shown. For more examples, see Fig. S16. Scale bar = 10 μm. (**B**) UBA1^M41V^ THP1 monocytes exhibit mitochondrial morphology abnormalities, indicative of impaired function. UBA1^WT^ and UBA1^M41V^ THP1 cells were treated with dox for 10 d and analyzed by transmission electron microscopy. Scale bar = 1 μm. Red arrowheads point to dilated cristae often seen in UBA1^M41V^ mitochondria (see also Movie S2) (**C**) UBA1^M41V^ THP1 monocytes exhibit a higher percentage of mitochondria with dilated cristae per cell as compared to UBA1^WT^ control cells. Graphs depict quantifications of mitochondrial morphology on TEM images of the experiment shown in panel b. n > 20 cells, error bar = s.e.m, * = p < 0.05, ** = p < 0.01, student’s t-test. (**D**) VEXAS THP1 cells exhibit elevated levels of cytoplasmic mitochondrial DNA, as revealed by qPCR analysis of whole cell or cytoplasmic fractions of UBA1^WT^ and UBA1^M41V^ THP1 monocytes. n = 3 biological replicates with 2 technical replicates for two different mitochondrial DNA probes (see methods), error bar = s.e.m, ** = p < 0.01, student’s t-test. (**E**) Schematic representation of experimental design to test whether mtDNA release into the cytosol activates cGAS-STING-based VEXAS inflammation. (**F**) Blocking mtDNA release into the cytosol using either a scavenger of reactive oxygen species (MitoTEMPO), an inhibitor of VDAC oligomerization (VBIT4), or an inhibitor of the mitochondrial permeability transition pore (CsA) dampens STING signaling and attenuates VEXAS inflammation in THP1 UBA1^M41V^ model cells. UBA1^WT^ and UBA1^M41V^ THP1 monocytes were treated with indicated inhibitors (10 μM MitoTEMPO, 10 μM VDAC, and 1 μM CsA) for 24h and subjected to immunoblot analysis using indicated antibodies. Graphs depict quantifications of p-STING/STING, p-STAT1/STAT1, or p-STAT3/STAT3 ratio normalized to DMSO-treated UBA1^WT^ THP1 monocytes. n = 3 biological replicates, error bar = s.e.m, * = p < 0.05, ** = p < 0.01, student’s t-test. (**G**) Blocking mtDNA release into the cytosol restores STING localization by preventing its trafficking to the Golgi, as evidenced by anti-STING and anti-GM130 immunofluorescence analysis of THP1 model monocytes treated as described in panel F. (**H**) Quantification of the percentage of Golgi-localized STING of the experiment depicted in panel g. n ≥ 150 cells across 10 fields, error bar = s.e.m, **** = p < 0.0001, one-way ANOVA. (**I**) Verification that treatment of THP1 VEXAS model cells with indicated inhibitors blocks mtDNA release into the cytosol, as revealed by qPCR analysis of cytoplasmic fractions of UBA1^WT^ and UBA1^M41V^ THP1 monocytes. n = 3 biological replicates, 2 technical replicates for two different mitochondrial DNA probes (see methods), error bar = s.e.m, **** = p < 0.0001, one-way ANOVA. **(J)**Scheme depicting the molecular mechanisms triggering STING activation in VEXAS myeloid cells. Mitochondria in VEXAS myeloid cells are defective and release mtDNA into the cytosol, causing activation of cGAS-STING signaling, which drives multi-cytokine inflammation.

To test whether mtDNA leaked from dysfunctional mitochondria drives STING signaling, we blocked mtDNA release in THP1 VEXAS model cells using either a scavenger of reactive oxygen species (MitoTEMPO), an inhibitor of VDAC oligomerization (VBIT4), or an inhibitor of the mitochondrial permeability transition pore (CsA) (*38-40*) (***Fig. 4E***). Intriguingly, all three drugs dampened STING signaling and attenuated all axes of VEXAS inflammation in THP1 UBA1M41V monocytes (***Fig. 4F***) and also restored STING localization by preventing its trafficking to the Golgi (***Fig. 4G,H***), with the magnitude of phenotype correction closely matching the reduction in cytosolic mtDNA levels (***Fig. 4I***).

ERAD inhibition in THP1 monocytes via J1/G2 knockdown also induced mitochondrial pathway dysregulation and morphological abnormalities (***Fig. S18A-E***) but did not increase cytosolic mtDNA levels (***Fig. S18F***). Thus, while impaired ERAD contributes to mitochondrial dysregulation in VEXAS cells, it is not sufficient to drive mtDNA leakage. Instead, an additional disrupted cytoplasmic ubiquitylation event is required, consistent with our observations that aspects of VEXAS-associated inflammation in THP1 macrophages is only elicited upon combined ERAD and UBA1 inhibition (***Fig. S12F-H***).

Altogether, these results demonstrate that mitochondrial dysregulation leads to leakage of mtDNA, which acts as a ligand of cGAS in the cytoplasm of VEXAS cells to initiate STING signaling (***Fig. 4J***).

### Pharmacological inhibition of STING reverses inflammation in patient tissue

Finally, we examined whether pharmacological inhibition of STING would reverse inflammation in primary VEXAS patient tissue (***Fig 5A, Tables S7-9***). To this end, macrophages derived from patient or control donor peripheral blood were treated with either DMSO or H-151 and analyzed by bulk RNA-seq. Strikingly, treatment with STINGi H-151 led to a marked resolution of major inflammatory pathways in VEXAS patient-derived macrophages (***Fig. 5B, Tables S8,9***).

**Fig. 5.**
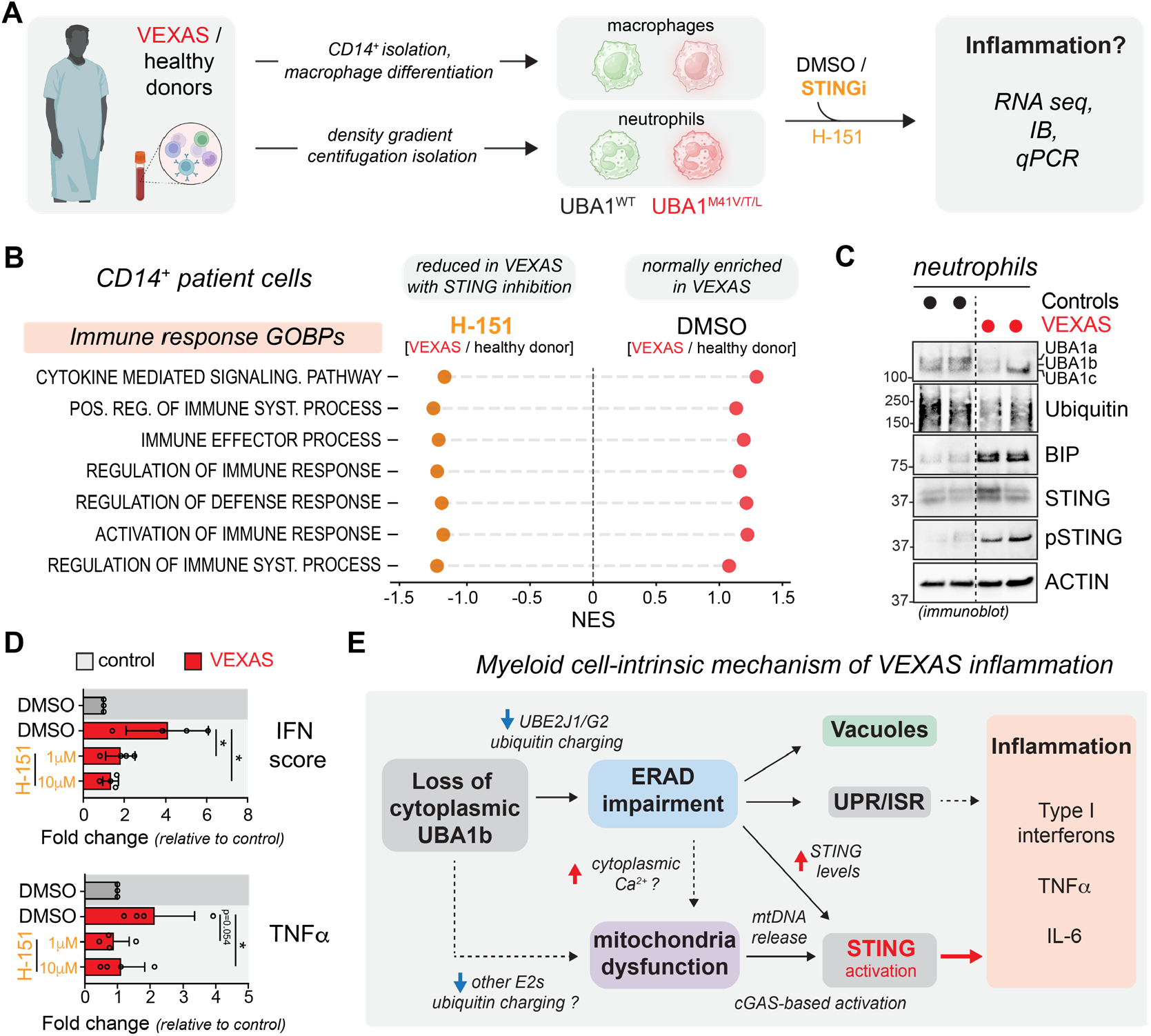
STING inhibition reverses inflammation in VEXAS patient myeloid cells. (**A**) Experimental scheme to test whether STING inhibition can reverse inflammation in VEXAS patient-derived CD14+ macrophages or neutrophils. (**B**) STING inhibition with H-151 (1 μM for 24 h) reduces inflammatory signaling in VEXAS myeloid cells, identifying the cGAS-STING pathway as a critical therapeutic target. Patient-derived healthy donor or VEXAS CD14^+^ monocytes were isolated, differentiated into macrophages, and treated as depicted in panel a and analyzed by RNA sequencing. The plot depicts normalized enrichment scores (NES) for immune response related GOBPs that are enriched in VEXAS over healthy donor cells (right side) and reduced in VEXAS cells upon STING treatment (left side). (**C**) The STING pathway is active in neutrophils from VEXAS patients, as shown via immunoblotting of cell lysates with indicated antibodies. (**D**) STING inhibition reduces inflammatory signaling in VEXAS neutrophils cells. Patient-derived healthy donor or VEXAS neutrophils were isolated, treated with 1 or 10 μM H-151 for 4 h and analyzed by qPCR analysis to determine mRNA levels of different type I interferon stimulated genes and TNFa. n = 3 biological replicates, error bar = s.e.m, * = p < 0.05, student’s t-test. (**E**) Model of cell intrinsic mechanisms causing VEXAS-associated inflammation in myeloid cells. Loss of cytoplasmic UBA1b activity due to canonical VEXAS mutations causes a preferential loss of ubiquitin charging of UBE2J1 and UBE2G2, inhibiting ER-associated degradation (ERAD). ERAD impairment results in accumulation of misfolded proteins, ER-derived vacuoles, activation of the unfolded protein and integrated stress responses (UPR/ISR), and accumulation of the innate immune adaptor STING. While UPR/ISR signaling contributes to some aspects of inflammation, cGAS-STING signaling predominantly drives the VEXAS-associated inflammatory response. cGAS-based activation is driven by mtDNA because of mitochondrial dysfunction. Together, our work identifies that the cGAS-STING axis as a critical regulator of multi-cytokine inflammation in VEXAS, nominating this pathway as a therapeutic target.

Recent studies have implicated neutrophils as a key mediator of disease in VEXAS, supported by evidence of neutrophilic infiltrates from patients (*41, 42*). Consistent with our findings in monocytes and macrophages, proteomic analysis of VEXAS patient-derived neutrophils revealed that ERAD and inflammatory pathways are amongst the most dysregulated (***Fig. S19A,B***) and STING levels are significantly increased (***Fig. S19C***). In addition, we detected elevated BiP expression and phosphorylation of STING at Ser366 (***Fig. 5C***), indicative of ER stress and active STING signaling. Correspondingly, VEXAS neutrophils exhibited elevated type I interferon and TNFα levels, both of which were reduced following 4-hour treatment with the STING inhibitor H-151 (***Fig. 5D***). Similar results were observed in a HL-60 promyelocyte– and neutrophil-differentiated base-edited VEXAS model (***Fig. S19D-G***). Together, these data identify cGAS-STING as a therapeutic vulnerability in VEXAS patients across multiple myeloid cell types.

We next attempted to extend these findings to an in vivo hematopoiesis model using healthy donor human HSPCs base-edited to express UBA1^M41T^, followed by xenotransplantation, a system previously shown to recapitulate key features of VEXAS (*14*). We found that VEXAS mutant cells diminished over time, most prominently in the context of STING knockout, suggesting a functional co-dependence between mutant UBA1 and STING signaling while precluding downstream inflammatory analyses (***Fig. S20A-D***).

Collectively, our findings suggest that the early, cell intrinsic inflammatory mechanisms in VEXAS arise from loss of cytoplasmic ubiquitylation, causing ERAD dysregulation and subsequent ER-vacuolization, UPR activation, and STING accumulation. This expanded STING reservoir is activated in a cGAS-dependent manner by excess cytosolic mtDNA, driving pathological, multi-cytokine inflammation (***Fig. 5E***).

## DISCUSSION

This study advances our understanding of VEXAS pathogenesis by dissecting the interplay between UBA1, ERAD, mitochondrial homeostasis, and inflammatory responses in mature immune cells. Despite the pleiotropic effects of global cytoplasmic ubiquitylation loss and the likely many cellular consequences, we show that ERAD dysregulation leads to accumulation of its endogenous substrate, STING. We further show that parallel mitochondrial dysfunction and mtDNA leakage leads to chronic STING signaling in a cGAS-dependent manner. We thus define a major mechanism through which cell-autonomous inflammation occurs in VEXAS.

Importantly, in our study, we show that inflammation occurs without spontaneous apoptotic, necroptotic, or pyroptotic cell death. This is consistent with a model in which inappropriate cGAS-STING activation initiates pro-inflammatory cytokine production (e.g. TNF α, IL-6, and Type-I interferon) through a cell-intrinsic mechanism. This primary inflammatory state may then promote abnormal TNFα-mediated inflammatory cell death of VEXAS mutant cells, which recently was shown to propagate tissue inflammation (*19,20,43*). Furthermore, STING hyperactivation may help explain the constellation of some of the unique symptoms observed in VEXAS, including lung and skin inflammation, which share similarities with other STING associated vasculopathies such as COPA Syndrome and STING-associated vasculopathy with onset in infancy (SAVI) (*44-49*).

Beyond its critical role in initiating and maintaining multi-cytokine inflammation in mature myeloid cells, our data suggests that inappropriate STING signaling may contribute to additional aspects of VEXAS, including hematopoietic abnormalities. In HSPCs, loss of cytoplasmic UBA1 function has been shown to lead to UPR activation, allowing for PERK-dependent survival of early progenitors (*17*). We further these observations by showing that across multiple myeloid cell types, major effectors of VEXAS, and HSPCs, loss of cytoplasmic ubiquitylation causes ERAD failure which drives UPR activation, and additionally, vacuole formation of expanded ER, and ultimately STING activation. These observations imply a cell-autonomous contribution of aberrant STING signaling to hematopoietic defects in VEXAS, consistent with previous reports implicating STING in promoting myeloid differentiation and commitment of HSPCs (*50,51*).

However, key questions remain regarding the critical factors that determine the non-cell-autonomous manifestations of the disease, such as competitive advantages of mutant clones, and other cell-specific effects including plasma cell expansion, multiple myeloma, and anemia.

Interestingly, ERAD is known to play a critical regulatory role in the maturation of both Band T-cells (*52,53*). Classically, VEXAS mutations are predominantly myeloid restricted with patients possessing no UBA1 mutant B- or T-cells; however, patients do possess UBA1 mutant NK-cells. Thus, aberrant ERAD in UBA1 mutant progenitors may help explain this lineage restriction. ERAD activation has also been implicated in acute myeloid leukemia (AML), suggesting a possible role for ERAD/UBA1 inhibition in preventing disease progression (*54*). Similarly, common genetic variants in UBE2G1, an ERAD related E2, have been linked to increased risk of clonal hematopoiesis (*55*).

More broadly, our findings position VEXAS within an expanding class of disorders in which failure of ubiquitin-proteasome-mediated protein quality control is directly coupled to chronic innate immune activation. Perturbations in ubiquitylation or proteasomal degradation have previously linked extreme proteostatic burden to monogenic autoinflammatory syndromes such as OTULIN-related autoinflammatory syndrome (*56*) and PSMB8-associated CANDLE/PRAAS (*18*), as well as to acquired inflammatory states including multiple myeloma and other plasma cell dyscrasias (*57*). Across these contexts, impaired clearance of misfolded or regulatory proteins can act as a primary inflammatory trigger, engaging stress-responsive and innate immune pathways independently of classical pathogen sensing or overt cell death. Our data extend this framework by revealing a generalizable mechanism whereby disruption of ubiquitin-dependent proteostasis, through ERAD failure, licenses innate immune activation via accumulation and sensitization of STING, coupled to secondary mitochondrial dysfunction. In this model, defects in protein quality control integrate disturbances in 1) ER homeostasis, 2) mitochondrial integrity, and 3) cytosolic DNA surveillance which converge to sustain cGAS–STING signaling. This conceptual framework might help explain the inflammatory phenotypes seen across a broad range of human disease characterized by proteostatic stress, including other autoinflammatory syndromes, neurodegeneration, cardiovascular disease, and cancer (*16,58-60*).

## MATERIAL & METHODS

### Cell culture and reagents

Human THP1 cells (ATCC, TIB-202) were cultured at 37 °C in a humidified atmosphere with 5% CO2 in RPMI 1640 Medium (Gibco) supplemented with 10% heat-inactivated FBS, 100 U/mL Penicillin-Streptomycin (Gibco), 10 mM HEPES, 1 mM sodium pyruvate, 4500 mg/L glucose, and 1500 mg/L sodium bicarbonate. Differentiation of THP1 monocytes into macrophages was induced using 20 nM phorbol 12-myristate 13-acetate (PMA) (MedChemExpress, HY18739) supplemented in standard media for 48-72 h. Human HL-60 cells (ATCC, CCL-240) were cultured at 37 °C in a humidified atmosphere with 5% CO2 in RPMI 1640 Medium (Gibco) supplemented with 20% heat-inactivated FBS and 100 U/mL Penicillin-Streptomycin (Gibco). Differentiation of HL-60 cells into neutrophils was induced using 1 μM all-trans-Retinoic acid (ATRA; MilliporeSigma, R2625) treatment for 5 days, validating differentiation by flow cytometry for positive human CD11b expression (Biolegend, 101207). Chinese Hamster Ovary ts20 cells were cultured in complete CHO ts20 medium MEM α (Gibco, 12571063) supplemented with 1.8 g/L glucose, 10% FBS, and Penicillin-Streptomycin 100 U/mL as previously described at 30.5 °C in a humidified atmosphere with 5% CO2 (*11*). CD34+ cells were isolated from peripheral blood or bone marrow from male donors and cultured at a concentration of 5×10^5^ cells/mL in a “HSPC medium” containing StemSpan (StemCell Technologies) supplemented with 100 U/mL Penicillin-Streptomycin (Gibco) and the following recombinant human cytokines: human stem cell factor, Fmslike tyrosine kinase receptor 3 ligand, thrombopoietin, interleukin-3 and interleukin-6 (StemSpan expansion supplement, StemCell Technologies). 32D cells (DSMZ, ACC 411) were cultured in RPMI 1640 medium (Gibco) with 10% heat-inactivated FBS, 100 U/mL Penicillin-Streptomycin (Gibco), and 2 ng/mL recombinant mouse interleukin-3 (IL-3) (PeproTech, 213-13-10UG).

Cells were treated for specified time periods with the following: H-151 (1 μM or 10 μM; MedChemExpress, HY-112693), SN-011 (10 μM; MedChemExpress HY-145010), MitoTEMPO (10 μM; MedChemExpress, HY-112879), Cyclosporin A (1 μM; MedChemExpress HY-B0579), VBIT4 (10 μM; MedChemExpress, HY-129122), 4μ8c (100 μM; MedChemExpress, HY-19707), GSK2606414 (5 μM; MedChemExpress, HY-18072), Ceapin-A7 (6 μM; MedChemExpress, HY-108434), PKR-IN-C16 (0.6 μM; MedChemExpress, HY-13977A), GCN2-IN-1 (1 μM; MedChemExpress, HY-100877), CP26 (1 μM, 1.25 μM, 2 μM, and 5 μM; MedChemExpress, HY-49116), Eeyarestatin I (10 μM; MedChemExpress, HY-110078), TAK-243 (5 nM; MedChemExpress, HY-100487), cyclohexamide (50 µg/mL; Sigma Aldrich, C4859), BafA1 (1 μM or 10 μM; MedChemExpress, HY-100558), and NSC697923 (3 μM or 4 μM; MedChem Express, HY-13811).

### Human subjects and patient sample processing

The Ethical Review Boards of NYU (NCT06004349) approved the study. All investigations were performed in accordance with the ethical standards of the 1964 Declaration of Helsinki and its later amendments. Consent was obtained from all subjects for the experiments performed. Patient and control blood was collected in sodium heparin BD Vacutainer tubes and PBMCs (peripheral blood mononuclear cells) and BMMNCs (bone marrow mononuclear cells) isolated within 6 h from collection via density centrifugation of blood using Lymphoprep (Stem Cell Technologies, 18061) in 50 mL SepMate Tubes (Stem Cell Technologies, 85450). Isolated PBMCs were then further purified by magnetic-activated cell sorting (MACS) utilizing human CD14^+^ microbeads (Miltenyi Biotec, 130-097-052) in a QuadroMACS Separator (Miltenyi Biotec, 130-097-052) with LS Columns (Miltenyi Biotec, 130-042-401) per manufacturer’s directions. Patient information included in Table S9. CD14^+^ human PBMCs media-free pellets were then either snap-frozen in liquid nitrogen for downstream proteomics and transcriptomics or differentiated into macrophages by resuspending cells in pre-warmed RPMI 1640 media supplemented with 10% heat-inactivated FBS, 100 U/mL Penicillin-Streptomycin, and 50 ng/mL human GM-CSF (PeproTech, 300-03-50UG). Differentiation media was changed every 2 days with full differentiation observed by day 6. Isolated BMMNCs were further purified into CD34^+^ populations using human CD34 microbeads (Miltenyi Biotec, 130-046-702) in a manner identical to CD14+ MACS purification. Neutrophil isolation was conducted as previously published (*61*). Patient information for samples used in manuscript is included in ***Table S10***.

### Proteomic sample preparation

For mass spectroscopy experiments, THP1-UBA1 cKO cells induced with doxycycline were collected after 10 days of induction. In brief, cells were harvested by resuspension of settled cells, washed in PBS, and centrifuged at 300 × g for 5 min. The cell pellets were stored at −80 °C or directly used for isolation of protein peptides for MS analysis. Cell pellets were resuspended in ∼0.5-1ml of ice-cold TB buffer (20 mM HEPES [pH 7.3], 110 mM potassium acetate, 2 mM magnesium acetate, 50 mM NaCl, 1 mM EGTA, 0.1% NP-40, protease inhibitors (Roche) and incubated on ice for 30min. Cells were further lysed by 3 cycles of high frequency sonication with 10s on 50s off cycles to prevent heating of the sample. Lysates were subsequently cleared by centrifugation at 4°C at 15,000 × g for 30 min and filtered through a 0.22-μm filter (Millex-GV) to remove residual lipids. Isolated proteins were precipitated by adding TCA to 20% (v/v) followed by overnight incubation on ice. Precipitated proteins were collected by centrifugation, 10 min at max rpm. Pelleted proteins were washed three times with ice-cold 90% acetone in 0.01 M HCl and briefly allowed to air-dry. Dried proteins were prepared for mass spectrometry analysis following standard protocol for S-TrapTM mini columns (Protifi). Tryptic peptides were dried under vacuum and stored at -20 °C until LC-MS/MS analysis.

### DiGly-peptide isolation and enrichment

DiGlycine (K-ε-GG)-peptides were enriched from isolated total proteomic peptides using the PTMScan® Ubiquitin Remnant Motif (K-ε-GG) Kit (Cell Signaling Technologies). In brief, isolated dehydrated tryptic peptides previously purified by reversed-phase, solid-phase extraction (S-trap), were applied to an immunoaffinity purification column with the PTMScan® Motif antibody conjugated to protein A agarose beads. Unbound peptides were removed through wash and (K-ε-GG)-enriched peptides were eluted with acid, following the manufacturer’s recommendations. Desalting and isolation of enriched peptides was performed using the provided reversed-phase purification microtips. Subsequent (K-ε-GG)-enriched peptides were concentrated and submitted for MS analysis.

### LC-MS/MS analysis

For global proteomics analysis, nanoLC-MS/MS analysis of tryptic peptides were carried out with a Thermo Scientific Fusion Lumos tribrid mass spectrometer interfaced to a UltiMate3000 RSLCnano HPLC system. For each analysis, 1 µg of the tryptic digest was loaded and desalted in an Acclaim PepMap 100 trapping column (75 µm × 2 cm)) at 4 µL/min for 5 min. Peptides were then eluted into an Aurora Ultimate TS 25×75 C18 UHPLC column (Ion Opticks, Australia) and chromatographically separated using a binary solvent system consisting of A: 0.1% formic acid and B: 0.1% formic acid and 80% acetonitrile at a flow rate of 300 nL/min. A gradient was run from 1% B to 32% B over 110 min, followed by a 5-min wash step with 90% B and 14-min equilibration at 1% B before the next sample was injected. Precursor masses were detected in the Orbitrap at R=120,000 (m/z 200) in profile mode. Fragment masses were detected in the Orbitrap at R=15,000 (m/z 200) in centroid mode. Data independent MSMS was carried for m/z 400-1000 with 13 m/z windows. AGC target was set at 200%, and maximum injection time was 20 ms.

Enriched diGly peptides were analyzed with the same instrument and same chromatography method using data dependent acquisition with “top of speed” settings. Cycly time was 2 sec with 60 sec dynamic exclusion. Fragment ions were detected in the orbitrap at R=30,000 (m/z 200). All samples were analyzed in block randomized order.

### Mass spectrometry data analysis

Protein identification and quantification were carried out using Spectronaut 19 (Biognosys, Switzerland) using Direct DIA with software default settings. Raw data was searched against a Human protein database from Uniprot (2023.08.09). Methylthio (+46) of C was set as fixed modification, oxidation of M and acetylation at protein N-terminal were set as variable modification. Quantification was based on MS2 area, background signal was used for imputation with missing values.

Data from enriched diGly peptides were processed using Proteome Discoverer software package (v.2.5, Thermo Scientific). Raw data was searched against a human protein database from Uniprot (2023.08.09) along with a contaminant protein database with Sequest HT search engine. C alkylation was set as fixed modification, M oxidation, K ubiquitylation (-GG), and protein N-terminal alkylation and ubiquitylation were set as variable modifications. Peptide quantification was based on precursor intensity.

### Transcriptomics

RNA was extracted from both cell line and primary patient models using RNeasy Plus Mini Kit (Qiagen, 74134) per manufacturer’ s guidelines. Bulk RNA NGS was performed by Psomagen, Inc using TruSeq Stranded mRNA library prep kit. mRNA sequencing was performed on Illumina NovaSeqX with TruSeq stranded mRNA library kit. The 151 × 2bp reads were pre-processed and aligned using a custom pipeline (https://doi.org/10.5281/zenodo.5550459). Quality metrics were first assessed with FastQC (Andrew, S. FastQC: a quality control tool for high throughput sequence data (Brabham Bioinformatics, 2010) and FastQScreen (*62*) and unwanted sequences were removed with Trimmomatic (parameters TRAILING:5 SLIDINGWINDOW:4:15 MINLEN:35) *(63*). Additional validation was performed, and high-quality reads were aligned to GRCh38.p13 using Gencode v38 gene annotations (*64*) with STAR (*65*) (parameters -outFilterMismatchNoverLmax 0.2, --outFilterMultimapNmax 1, --outFilterType BySJout, --twopassMode Basic). Picard (“Picard Tookit.” 2019. Broad Institute, Github Repository. https://github.com/broadinstitute/picard; Broad Institute) was employed to tabulate sample level alignment statistics without strand specificity and unstranded reads were counted per gene with featureCounts (*66*). Single cell RNA-seq (scRNA-seq) was performed as previously described *(13*).

### Differential Enriched Gene and Pathway Analysis

Bulk RNA-seq transcriptomic data was analyzed for differential expression with R package (*67*). Initial steps included filtering hemoglobin-encoding and lowly expressed genes that could skew normalization (minimum of 10 counts in at least 2 samples). DESeq2’s variance stabilization transformation was fit to data and differentially expressed genes were determined on Benjamini-Hochberg adjusted pvalue < 0.05 and |log2FoldChange| > 1 thresholds. Genes were preranked by sign(log2FoldChange) × -log10(p-value) as input for gene set enrichment analysis (GSEA) (*68*). Enriched MSigDB gene sets (v2014.1) (*69*) were determined with fgsea *(70*). EdgeR (*71*) functions calcNormFactors and cpm were used to normalize counts by Trimmed Mean of M-values (TMM) for intersample comparison. Differentially expressed proteins were determined by unpaired t-test without additional adjustment, with significance thresholds set as pvalue < 0.05 and |log2FoldChange| > 0.6. Proteins only detected in one condition were omitted from statistical analysis. Pathway enrichment and functional effect analysis were carried out the same as performed for mRNA.

For scRNA-seq, differential gene expression analysis was performed between cells from VEXAS patients and from age-matched healthy controls for both CD14 monocytes and HSCs. For the CD14 monocyte analysis, donors with < 500 CD14 monocytes were excluded. Each donor was then randomly downsampled without replacement to retain 500 CD14 monocytes per donor (n = 4 healthy control donors, n = 7 VEXAS donors)(13, 17) (retaining n = 2000 healthy control cells and n = 3500 VEXAS cells for analysis. For the HSC analysis, all donors were retained due to lower cell counts (n = 4 healthy control donors, n = 10 VEXAS donors). Donors with > 200 HSCs were downsampled to 200 cells for analysis, without replacement, retaining n = 794 healthy control cells and n = 1733 VEXAS cells for analysis. Mitochondrial and ribosomal genes were removed from the gene count matrix prior to analysis, and then log normalization was run with default parameters for Seurat’s NormalizeData function (scale.factor = 10000, normalization. method = “LogNormalize”, margin = 1). Genes expressed in < 20% of cells tested were excluded from analysis. Differential expression analysis was performed using a Wilcoxon rank sum test. Gene set enrichment analysis was performed using the GOBP ERAD pathway, which is publicly available in msigdb and was obtained via the msigdbr package (v7.5.1). The fgsea package (v1.24.0) was then used to perform ranked gene set enrichment analysis. Genes were ranked by effect size (avg_log2FC).

### Gene module score analysis

HSCs and CD14 monocytes from VEXAS and healthy control donors were scored for enrichment of the GOBP ERAD, UBIQUITIN DEPENDENT ERAD, and REGULATION OF ERAD gene modules using the AddModuleScore function from Seurat within scRNA seq data. First, mitochondrial and ribosomal genes were removed, and then log normalization was run as described above. Then, briefly, the function infers the gene set activity by calculating the average expression levels of each gene set on a single-cell level, subtracted by the aggregated expression of control feature sets. All analyzed features are binned based on averaged expression, and the control features are randomly selected from each bin. Default parameters were used. Distribution of module scores were compared between VEXAS and control cells and a two-sided Wilcoxon rank sum test was used to determine significance.

### Construction of a THP1 VEXAS model cell line

To generate an inducible UBA1-/- cell line, human THP1 cells were first infected with lentivirus packaged with FUCas9Cherry (Addgene, 70182) to constitutively express Cas9 and mCherry. Next, ten UBA1 targeting guide sequences were generated using CHOPCHOP software(67). The doxycycline-inducible FgH1tUTG (Addgene, 70183) plasmid was digested with BsmBI-V2 (NEB, R0739S), DNA gel electrophoresis purified, and T4 DNA Ligase (NEB, M0202S) ligated with the annealed product of a forward (5’-TCCCTAACCAGGACAACCCCGGTG-3’) and reverse (5’-AAACCACCGGGGTTGTCCTGGTTA-3’) oligo containing BsmBI overhangs (underlined) and the UBA1-targeting sequence for the gRNA. The noted UBA1 target sequence was the most efficient in inducing KO of UBA1. This new plasmid, FgH1tUTG-UBA1, was then lentivirally packaged and infected into the Cas9 expressing THP1. These cells were then sorted using FACS on a Sony SH800 Cell Sorter with a 130 μm sorting chip, selecting for dual positive mCherry (FUCas9Cherry) and GFP (FgH1tUTG-UBA1) THP1 populations. Double positive cells were then induced with 2 μg/mL doxycycline with UBA1-/- validated by loss of endogenous UBA1a/b on immunoblot and eventual cell death.

Next, pHAGE_puro (Addgene, 118692) was modified to contain an IRES regulating puromycin expression and encode either a modified UBA1WT (pHAGE-UBA1_WT-IRES-Puro) or UBA1M41V (pHAGE-UBA1_M41V-IRES-Puro) cDNA product with a FLAG-HA tag. To avoid Cas9-mediated KO of the UBA1 transgene, the cDNA regions corresponding to the gRNA target sequence were modified to contain synonymous mutations (in bold) in the codons corresponding to amino acids 213-222 (5’-GTC ACG AAA GAT AAT CCT GGA GTT GTG ACT-3’). These constructs were then packaged into lentivirus, transduced into the dox-inducible UBA1-/- THP1 cell line, and selected for 7 days in 1 μg/mL puromycin. After selection, trans-UBA1 expression was validated with immunoblot against HA-tag. In addition to generating UBA1M41V (c.121 A>G) we also generated UBA1M41L (c.121 A>T) (pHAGE-UBA1_M41L-IRES-Puro) as an additional control for loss of UBA1b and VEXAS related studies.

Finally, 3×10^6^ control UBA1WT or UBA1M41V expressing inducible UBA1-/- THP1 cells were plated in a 10-cm diameter tissue culture dish with 15 mL of media containing 2 μg/mL doxycycline and grown for 6 days. For monocyte experimentation, on day 6, cells were then split 1:3 into new media containing 2 μg/mL doxycycline. Media was changed every day by aspirating half the volume and replacing with new media prior to being harvested on day 9. For macrophage experimentation, induced cells were counted, spun at 300 × g for 10 min, and supernatant removed. Cell pellets were then resuspended in new media containing 2 ug/ mL doxycycline and 20 nM PMA at a concentration of 1×10^6^ cells/mL. These were then plated on tissue culture treated 6-well plates. Adherence and macrophage differentiation was observed by day 8 and cultures harvested on day 9.

### Quantitative PCR of cytokines and interferon score calculation

Quantitative PCR was performed using TaqMan Universal PCR Master Mix (Applied Biosystems, 4304437) on a QuantStudio 5 Real-Time PCR System. Each reaction utilized cDNA isolated using SuperScript IV Reverse Transcriptase (Invitrogen, 18090010) generated from 200 ng of total RNA input. Cytokines TNFα (Applied Biosystems, Hs01113624_ g1) and IL-6 (Applied Biosystems, Hs00174131_m1) were measured relative to housekeeping genes 18S (Applied Biosystems, Hs99999901_s1) and HPRT1 (Applied Biosystems, Hs02800695_m1). For interferon score, expression of IFI44L (Applied Biosystems, Hs00199115_ m1), IFI27 (Applied Biosystems, Hs01086370_m1), IFIT1 (Applied Biosystems, Hs00356631_g1), ISG15 (Applied Biosystems, Hs00192713_m1), RSAD2 (Applied Biosystems, Hs01057264_m1), and SIGLEC1 (Applied Biosystems, Hs00988063_m1) were measured relative to housekeeping genes 18S (Applied Biosystems, Hs99999901_s1) and HPRT1 (Applied Biosystems, Hs02800695_m1). The arithmetic mean of the six individual ISG fold changes was taken to generate the final interferon score for each sample. All assays were performed in technical duplicates with biological triplicates. Samples were normalized to the geometric mean of 18S and/or HPRT1 expression using the comparative Ct method (2^(-ΔCt)).

### Cytokine measurements

Cytokine measurements were performed using commercial enzyme-linked immunosorbent assay kits according to manufacturer’s instructions for human IL-6 (R&D Systems, D6050), IL-1β (R&D Systems, DLB50), and TNF-α (R&D Systems, DTA00D). Briefly, cell culture supernatants were harvested, spun down for 30 min at 18,000 × g at 4 °C and assayed in technical duplicate. Optical density was determined using a microplate reader set to 450 nm with wavelength correction at 540 nm. Cytokine concentrations were calculated using a standard curve generated with recombinant human proteins provided in the kits.

### Vacuole imaging and quantification with light microscopy

Samples were washed with PBS and resuspended to a concentration of 1×105 cells/mL. 200 μl of sample was then cytospun at 800 rpm for 4 min onto microscope slides (Corning® 75×25 mm) and fixed with methanol at room temperature for 8 min. After fixing, cells were washed with PBS and stained with Wright-Giemsa Stain (Azer Scientific) for 15 min. The cells were washed with PBS followed by MilliQ water for 5 min. Cells were mounted by toluene with glass coverslips and allowed to cure for 4 h before imaging. All vacuole quantifications were performed using Nikon NIS Segment.ai. Training data for cell segmentation and vacuole counting was generated in NIS elements by first converting cell images to gray and auto contrasting. Vacuoles that were present were marked either by auto detection or were manually traced using the program. The program was trained on the annotated images for 1000 iterations. Once the training data was generated, images of cells and vacuole quantifications from all experiments were analyzed by NIS Segment.ai using the trained set.

### Immunofluorescence of Vacuoles

Immunofluorescence was performed as previously described (*73*). Briefly, cells were cytospun onto microscope slides (Corning 75×25 mm) and fixed with 4% paraformaldehyde in PBS for 30 min at room temperature. Cells were subsequently washed with phosphate buffered saline for 3 × 5 min. Following wash, cells were blocked in 2% bovine serum albumin (BSA)/0.1% Triton X-100 in PBS (0.1% PBST) for 30 min and then incubated with sheep anti-TGN46 (1:100; BioRad, AHP500GT) or rabbit Anti-Calnexin (Cell Signaling, 2433S) primary antibody diluted in 2% BSA/0.1% PBST for 1 h at room temperature. Primary antibody was washed off 3 × 5 min with PBST. Cells were incubated with Alexa Fluor 488 AffiniPure Donkey Anti-Sheep (1:500; Jackson Immuno, 711-545-152), Alexa-647 Donkey Anti-Rabbit (1:500; Jackson Immuno, 711-605-152), or CF647 Donkey Anti-Rabbit IgG (1:500; Biotium, 20811) secondary antibody diluted in 2% BSA/0.1% PBST for 30 min at room temperature. Lastly, cells were incubated with Hoechst 33342 (1:500; R&D systems, 5117) diluted in 2% BSA/0.1% PBST for 5 min at room temperature. Cells were washed 3 × 5 min with 0.1% PBST and mounted by glass coverslips with ProLong Gold Antifade Mountant (ThermoFisher, P36930) to cure overnight.

### Immunoblotting

Immunoblotting was performed as previously described (*11*).Antibodies targeting Ubiquitin-conjugating enzymes (E2s) included UBE2A (Protein Tech, 11080-1-AP), UBE2B (Invitrogen, MA5-42600), UBE2C (Protein Tech, 66087-1-Ig), UBE2D1/2/3/4 (Cell Signaling, 4330S), UBE2E1 (Protein Tech, 55457-1-AP), UBE2E2 (Protein Tech, 11844-1-AP), UBE2E3 (Protein Tech, 15488-1-AP), UBE2F (Protein Tech, 17056-1-AP), UBE2G2 (Cell Signaling, 63182S), UBE2H (Protein Tech, 15685-1-AP), UBE2I (Protein Tech, 14837-1-AP), UBE2J1 (Santa Cruz Biotechnology, sc-377002), UBE2K (Protein Tech, 11834-3-AP), UBE2L3 (Protein Tech, 14415-1-AP), UBE2L6 (Protein Tech, 17278-1-AP), UBE2M (Protein Tech, 67482-1-Ig), UBE2N (Protein Tech, 10243-1-AP), UBE2O (Protein Tech, 15812-1-AP), UBE2Q2 (Protein Tech, 12581-1-AP), UBE2R1 (Protein Tech, 10964-2-AP), UBE2R2 (Protein Tech, 14077-1-AP), UBE2S (Protein Tech, 14115-1-AP), UBE2V1 (Protein Tech, 10207-2-AP), UBE2V2 (Protein Tech, 10689-1-AP), and UBE2Z (Protein Tech, 16928-1-AP).

Other primary antibodies used in this study included those targeting HA-tag (Biolegend, 901501), β-actin (Cell Signaling, 3700S), UBA1a/b (Cell Signaling, 4891S), GFP (Protein Tech, 66002-1-Ig), PERK (Cell Signaling, 3192S), phospho-PERK (Invitrogen, PA5-40294), IRE1α (Cell Signaling, 3294S), phospho-IRE1α (Invitrogen, PA1-16927), ATF6 (Cell Signaling, 65880S), PKR (Cell Signaling, 12297S), phospho-PKR (Abcam, AB32036), GCN2 (Cell Signaling, 3302S), phospho-GCN2 (Cell Signaling, 94668S), XBP1s (Cell Signaling, 12782S), BiP (Cell Signaling, 3177S), ATF4 (Cell Signaling, 11815S), CHOP (Cell Signaling, 2895S), eIF2α (Cell Signaling, 5324S), phospho-eIF2α (Cell Signaling, 3398S), GADD34 (Protein Tech, 10449-1-AP), HRD1 (Protein Tech, 13473-1-AP), STING (Cell Signaling, 13647S), phopsho-STING (Cell Signaling, 19781S), cGAS (Cell Signaling, 15102S), RIG-I (Cell Signaling, 3743T), MDA-5 (Cell Signaling, 5321T), IRF-3 (Cell Signaling, 11904T), phospho-IRF3 (Cell Signaling, 4947T), IRF7 (Cell Signaling, 4920S), phospho-IRF7 (Cell Signaling, 5184S), TBK (Cell Signaling, 3504T), phospho-TBK (Cell Signaling, 5483T), STAT1 (Cell Signaling, 14994S), phospho-STAT1 (Cell Signaling, 9167S), STAT3 (Cell Signaling, 9139S), phospho-STAT3 (Cell Signaling, 9145S), pro-IL-1β (Cell Signaling, 12242S), Caspase 1 (Cell Signaling, 3866T), Cleaved Caspase 1 (Cell Signaling, 4199T), Caspase 3 (Cell Signaling, 14220S), Cleaved Caspase 3 (Cell Signaling, 9661S), Caspase 8 (Cell Signaling, 4790S), Cleaved Caspase 8 (Cell Signaling, 9496S), RIPK1 (Cell Signaling, 3493S), phospho-RIPK1 (Cell Signaling, 65746S), MLKL (Cell Signaling, 14993S), phospho-MLKL (Cell Signaling, 91689S), Gasdermin D (Cell Signaling, 39754S), Cleaved Gasdermin D (Cell Signaling, 36425T), ORF1p (Cell Signaling, 88701S), and SERCA2 (Cell Signaling, 9580S). Lastly, HRP-conjugated secondary antibodies used in this study were used at a concentration of 1:1000 (anti-rabbit [Cell Signaling, 7074S] or anti-mouse [Cell Signaling, 7076S]) and visualized using Immobilon Western Chemiluminescent HRP Substrate (low sensitivity) (Millipore, WBKLS0500) or Immobilon ECL Ultra Western HRP Substrate (high sensitivity) (Millipore, WBULS0500).

### In vivo E2 characterization using CHO ts20

Characterization of UBA1 variants on the E2 proteome was conducted as previously described (*2, 11*). Briefly, UBA1 variant cell lines were generated by lentiviral transduction of WT UBA1, single isoform UBA1 (UBA1a, UBA1b, or UBA1c) or VEXAS Syndrome UBA1 variants (M41V, M41L, S56F, or A478S) with 5 μg/mL puromycin selection. UBA1 variant CHO ts20 cell lines were harvested by trypsinization and pellet resuspended in complete CHO ts20 medium normalized to 1×106 cells/mL of media. Resuspended cells were then transferred to 1.5 mL microcentrifuge tubes and moved to pre-warmed thermomixers set to 39.5 °C. Samples were then incubated shaking at 500 rpm for 6 h. Following incubation, samples were analyzed by immunoblot in non-reducing conditions. Ratios of Ub∼E2/Total E2 were analyzed using ImageJ(*74*).

### ERAD activity assessment using a CD3δ-GFP reporter

Parental, UBA1^WT^, or UBA1^M41V^ THP1 cells were lentivirally infected with a plasmid expressing CD3δ-GFP (gift of Michelle Pagano Lab) and selected with 100 μg/mL hygromycin for 7 days before experimentation. For all experiments, cells were differentiated into macrophages with 48 h treatment in media containing 20 nM PMA. To test the efficacy of the reporter, parental THP1 containing the reporter were treated with either DMSO or 5 μM CP26. Cells were then treated with 50 μg/mL cycloheximide (CHX) (Sigma-Aldrich, C4859) and samples harvested a 0, 2, 4, and 6 h post CHX pulse. Samples were then immunoblotted against GFP and, in biological triplicate, CD3δ-GFP levels analyzed using ImageJ(74). UBA1WT, or UBA1M41V model THP1 cells containing the CD3δ-GFP reporter were examined in a similar fashion without the addition of DMSO or CP26.

### Expression and purification of UBA1 enzymes

UBA1aFLAG-HA (aa 1-1058), UBA1bFLAG-HA (aa 41– 1058), and UBA1bFLAG-HA (aa 67–1058) proteins were purified from HEK293T cells after transient transfection with pHAGE-UBA1-M41AM67AFLAG-HA, pHAGE-UBA1-41-1058-M67AFLAG-HA, or pHAGE-UBA1-67-1058FLAG-HA vectors (4 × 15-cm dishes). Cells were harvested by scraping in 1×PBS and centrifuged at 300 × g for 5 min. Cells were lysed in two pellet volumes of ice-cold lysis buffer (20 mM HEPES pH 7.3 containing 110 mM potassium acetate, 2 mM magnesium acetate, 50 mM NaCl, 1 mM EGTA, 0.1% NP-40, 1 × protease inhibitors (Roche), 1× Phos-Stop (Roche), and 2 mM phenanthroline. Cells were sonicated, and the lysates were cleared by centrifugation at 20,000 × g for 25 min. To remove residual lipids, the supernatant was filtered through a 0.22-μm filter (Millex-GV). Subsequently, the lysates were quantified using Pierce 660 nm reagent (ThermoFisher, 22660), and equal amounts of lysates were incubated with ANTI-FLAG-M2 agarose (Sigma) for 2 h at 4 °C. To remove endogenous interaction partners, beads were washed three times with lysis buffer containing 1 M NaCl, three times with lysis buffer containing 2% NP-40, and three times with lysis buffer, followed by elution from the beads with lysis buffer containing 3×FLAG peptide (0.5 mg/ml). UBA1a/b/c proteins were, concentrated, aliquoted, and snap-frozen in liquid N2and stored at −80 °C. Protein purity and quantity was determined by Coomassie gels.

### In vitro E2 ubiquitin charging assays

The ability of UBA1a/b/c to transfer ubiquitin to different E2s was either determined using Coomassie-based or ubiquitin-FITC fluorescence in vitro assays. For Coomassie based in vitro assays, 50 nM UBA1a/b/c was incubated with 1 uM E2 enzyme (1 of 24 different E2s, Boston Biochem, K-982), 100 uM ubiquitin (Enzo Life Sciences), and 5 mM ATP in UBA1 in reaction buffer (25 mM Tris-HCl [pH 8.0], 150 mM NaCl, 10 mM MgCl2) for 30 min at 30 °C. Reactions were quenched via addition of 2× urea sample buffer (150 mM Tris [pH 6.5], 6 M urea, 6% SDS, 25% glycerol, <0.1% bromophenol blue) and separated by non-reducing SDS-PAGE followed by Coomassie staining. For ubiquitin-FITC fluorescence in vitro assays, the reactions were performed using previously established protocols (Collins et al, 2024). In short, (i) 0.25 µM UBA1 in reaction buffer (25 mM Tris-HCl [pH 8.0], 150 mM NaCl, 10 mM MgCl2) was incubated with 10 µM ubiquitin-FITC and 10 mM ATP for 10 min at 30 °C. (ii) 1 µM of indicated E2 was added to initiate ubiquitin transfer for 30min. Reactions were quenched via addition of 2X urea sample buffer and separated by non-reducing SDS-PAGE. UBA1 and E2 charging were detected via a fluorescence scanner (Chemidoc MP, Biorad).

### Base editing of human HSPCs, 32D, and HL-60

Male donor bone-marrow derived CD34+ HSPCs were acquired from StemCell Technologies. For base editing, pCMV_ABEmax_P2A_GFP (112101; Addgene, Watertown, MA) was used. A single guide RNA (sgRNA) was designed to reproduce the p.Met41Thr c.122T>C mutation (Target Sequence: 5’-TGGCCATTCCCTAGGAATAG-3’). To induce a STING knockdown in HSPCs using single base-editing, a sgRNA was designed to induce a splicing mutation in the donor site of exon 4 (Target Sequence: 5’-GTCATACCTTGAGGCCCAGG-3’). We used chemically modified synthetic sgRNAs harboring 2′-O-methyl analogs and 3′-phosphorothioate nonhydrolyzable linkages at the first three 5′ and 3′ nucleotides (Synthego). 1×106 cells HSPCs per condition were electroporated two days after thawing with 3.0 μg of the ABE-encoding mRNA and 3.2 μg of the synthetic sgRNAs targeting UBA1, STING, or both using the P3 primary cell 4D-Nucleofector X Kit S (Lonza, Basel, Switzerland) and the CA-137 program (Nucleofector 4D). For editing HL-60, 4×105 cells were electroporated using 3.0 μg of the ABE-encoding mRNA and 3.2 μg of the synthetic sgRNA targeting UBA1 using the SF Cell Line 4D-Nucleofector X Kit S (Lonza, Basel, Switzerland) and the EN-138 program. 32D cells were edited as previously described (*22*).

Editing efficiency was assessed by genomic DNA extraction (Qiagen, 56304) and digital droplet PCR using a specific probe for UBA1 c. 122 T>C or the STING exon 4 splice site variant A>G. Reactions were performed using 2× ddPCR Supermix (Bio-Rad). 20 µL of PCR mixture and 70 μL of Droplet generation oil were mixed and droplet generation was performed using a Bio-Rad QX100 Droplet Generator. The droplet emulsion was thermally cycled and PCR amplification in the droplets was confirmed using Bio-Rad QX200 Droplet Reader. A patient with VEXAS syndrome and M41T mutation was used as a positive control to determine the threshold of positivity. All the data were evaluated above the threshold. QuantaSoft (Biorad) was used to analyze the % variant allele fraction data. Sanger sequencing after PCR amplification of the region of interest and Western blot validation were also performed.

### siRNA- and shRNA-mediated depletion of UBE2J1 and UBE2G2

For siRNA-mediated depletion, a 6-well culture plate was seeded with 1.25×10^6^ parental THP1 cells with media containing 20 nM PMA (∼12.5 ng/mL PMA) and allowed to differentiate into macrophages for 48 h. 24 μL of Hiperfect (Qiagen), 1.2 μL of 100 μM each siRNA stock targeting UBE2J1 (Dharmicon, D-007266-03-0005 and D-007266-01-0005) and UBE2G2 (Dharmicon, D-009095-02-0005 and D-009095-03-0005) were mixed in a 1.5 mL centrifuge tube with 400 μL of Opti-mem and briefly vortexed and let incubate at RT for 10 min. PMA-containing media on cells was aspirated, washed with sterile PBS, and 400 μL of preheated THP1 media with no PMA added. After 10 min the siRNA mixture was added dropwise and after 6 h 1.6 mL of regular THP1 media added. siRNA KD was assayed by qPCR, Western blot, and ELISA, 72 h after siRNA addition. As a control, a non-targeting siRNA pool (Dharmicon, D-001206-14-05) was used.

For shRNA-mediated depletion, sequences targeting either UBE2J1 (5’-GATGATATACCTACAACATTC-3’) or UBE2G2 (5’-GATGAGATTTACCTGTGAGAT-3) were selected. These sequences were then synthesized to follow a TargetSequence – 6-nt Loop – ComplementTargetSequence (where the 6-nt loop is 5’-CTCGAG-3’) format and cloned either into EZ-Tet-pLKO-Puro (Addgene, 85966) to create EZ-Tet-pLKO-UBE2J1-Puro or EZ-Tet-pLKO-Hygro (Addgene, 85972) to create EZ-Tet-pLKO-UBE2G2-Hygro. Human THP1 cells were then infected with lentivirus packaged with either construct and dual selected in 1 μg/mL puromycin and 200 μg/mL hygromycin. Induction of shRNA production was performed by adding 2 μg/mL doxycycline and assayed 72 h post induction.

### Transmission electron microscopy

Cells for TEM were prepared using the following protocol. First, samples were fixed in a mixture of 2.5% glutaraldehyde and 1% paraformaldehyde in 0.1 M Sodium cacodylate buffer, pH 7.4, for 60 min at room temperature and overnight at 4 ºC.

The following day, samples were washed three times for 10 min each in 0.1 M Sodium cacodylate buffer, pH 7.4, before being embedded in 4% low melting agarose (Sigma-Aldrich, A4018). Agarose embedded cell block was trimmed and post-fixed in 1% OsO4 in 0.1 M cacodylate buffer for 60 min on ice. Next, samples were rinsed and washed two times for 10 min each in water and incubated with 1% uranyl acetate overnight at 4 °C. The following day samples were rinsed and washed in water for 10 min and gradually dehydrated through a graded ethanol series followed by propylene oxide. Samples were then infiltrated in a gradient mix of propylene oxide and resin (Embed 812 resin) before being infiltrated with three changes of pure resin and embedded in 100% resin, and baked at 60 ºC for 48 h. Ultrathin sections (65 nm) were cut on an ultramicrotome (Leica EM UT7), and digital micrographs were acquired on JEOL JEM 1200 EXII operating at 80Kv and equipped with AMT XR-60 digital camera.

### FIB-SEM

After ultrathin sectioning, the blocks were cut with a saw blade and mounted on an SEM stub (Electron Microscopy Sciences, 75190) using double sided carbon tape with the cells facing up. Colloidal silver was used to coat the sides of the block to increase conductivity, followed by a 40 nm layer deposition of gold using a sputter coater (Electron Microscopy Sciences). FIB-SEM imaging was carried out with a Crossbeam 540 (Carl Zeiss Microscopy, White Plains, NY) using Atlas 5 (Fibics Inc., Ottawa, Canada). FIB-SEM images were acquired at a resolution of 9 nm in XY and a slice thickness of 9 nm. The SEM beam was used at 1.5 kV and 2 nA, and the FIB was used at 30 kV. Images were acquired with the enhanced back-scatter detector with a grid voltage set to 500 V. After volume acquisition, the image contrast was inverted, and the stack was registered using the Atlas 5 registration module.

### Segmentation of Volume EM Dataset

Subcellular structures in the volume EM dataset were segmented using a U-Net (75) model implemented with the segmentation_models (https://github.com/qubvel/segmentation_models) library based on Keras and TensorFlow. The U-Net architecture used a ResNet34 encoder pretrained on ImageNet to leverage feature representations. Ground truth annotations were manually generated using the Labkit plugin in ImageJ and 20 percent of the dataset was used for training and validation. The grayscale EM images were normalized and divided into non-overlapping 256×256 pixel patches using the patchify (https://github.com/dovahcrow/patchify.py) library for training. To enhance model generalization, data augmentation (geometric transformation: rotation, zoom, shear, and flipping) was applied using Keras ImageDataGenerator. The U-Net model was compiled using the Adam optimizer and binary cross-entropy as loss function and trained for 100 epochs with early stopping and model checkpoint callbacks to prevent overfitting. Model performance was monitored using the Intersection over Union (IoU) score as the primary evaluation metric. Two separate models were trained for the endoplasmic reticulum (ER) and mitochondria, respectively. During inference, output probability maps were thresholded at 0.5 to generate binary segmentations. Segmentation artifacts were manually inspected and corrected in ImageJ. Final binary masks were imported into Imaris (Oxford Instruments, ver. 9.9) for 3D volume reconstruction and quantitative analysis.

### Immunofluorescence of STING localization

Live, doxycycline-induced UBA1^WT^ or UBA1^M41V^ THP1 model monocytes were differentiated into adherent macrophages on poly-L-lysine cover slips. On day 9, cover slips were gently washed with PBS prior to fixation with 4% PFA for 15 min at room temperature. Following three PBS washes, cover slips were permeabilized with 0.1% Triton X-100 (MilliporeSigma) in PBS for 10 min at room temperature and washed three times with PBS. To quench GFP/RFP reporter signal in model macrophages, slides were incubated with cold 1% w/v sodium borohydride (MilliporeSigma) for 10 min. Cover slips were then visualized for GFP/RFP signal with the quenching step repeated until no fluorescence was appreciable. Cover slips were then incubated in blocking buffer (5% Normal Goat Serum + 1% BSA in PBS) for 60 min at room temperature. Cover slips were then incubated with both Mouse anti-GM130 (1:200; BD Bioscience, 610823) and Rabbit anti-STING (1:200; Cell Signaling, 90947S) overnight at 4 °C. Following three washes with PBS + 0.05% Tween-20, Cover slips were then incubated with Goat anti-mouse Cy3 (1:500; Jackson Immuno Research, 115-165-164) and Goat anti-rabbit Alexa Fluor488 (1:500; Jackson Immuno Research, 111-545-144) secondary for 60 min at room temperature. Coverslips were then washed and briefly stained with DAPI prior to mounting on slides using ProLong Gold Antifade Mountant (ThermoFisher, P36930).

Slides were imaged using a modified Nikon Eclipse Ti widefield microscope using an Olympus 60x objective. For each condition, twelve hyperstacks were acquired, at 200 μm × 200 μm × 6.25 μm with 27 z-steps and a step size of 0.25 μm. Z-stacks were centered on the z-position where nuclei appeared uniformly in focus. Three channels were used, with excitation wavelengths at 405 nm (DAPI), 488 nm (STING), 568 nm (GM130). Laser intensity and exposure time for each channel were standardized across all images. Analysis was performed by creating maximum intensity projections of DAPI, STING, and GM130 channels, and segmenting nuclei, cytoplasm, golgi, respectively, using the Cellpose 4.0 SAM model. Diameters for segmentation were set as 150 px for nuclei, 300 px for cytoplasm, and 50 px for golgi, with all other parameters left as the default setting. Masks were exported as.tif files and imported into a python script for further analysis. Segmentation accuracy was checked by eye after processing to confirm the parameters used were optimal. Additionally, each hyperstack was split into individual channel stacks and imported into the same python script. Mask files were used to assemble cell regions together. Only valid cells (cytoplasmic masks fully encompassing exactly one nucleus mask and one golgi mask) were used to record signal intensity. Signal intensities were calculated by summing all pixels found in regions defined by masks. The STING enrichment metric was calculated by taking the intensity recorded in the golgi mask and dividing it by the intensity recorded in the cytoplasmic mask and multiplying by 100 to generate a percentage of STING signal found within the golgi for each valid cell. The resulting tables for each condition were exported as.csv files from the python script for plotting and statistical analysis.

### Neon electroporation and gRNA-mediated KO of STING and cGAS

Electroporation was performed using the Neon Transfection System (Thermo Fisher Scientific, MPK5000) following the manufacturer’s guidelines with specific modifications. Briefly, a 100 µL Neon tip was used for each reaction and the electroporation parameters were optimized to 1700 volts, 20 milliseconds pulse width, and 1 pulse. These parameters were programmed into the device prior to electroporation. On day 0 of model induction, for each reaction 4×106 THP1 UBA1WT or UBA1M41V cells were washed with sterile PBS and resuspended in 100 μL Buffer R (Thermo Fisher Scientific, MPK10096). Chemically modified synthetic guide RNA (Synthego) targeting STING (5’- GGTGCCTGATAACCTGAGTA-3’), cGAS (5’- GACTCGGTGGGATCCATCG-3’), or a negative control scrambled sgRNA (Synthego, Pool #1) were added to each sample to achieve a final concentration of 400 nM. The cell-gRNA mixture was gently pipetted to ensure uniform distribution while avoiding bubble formation. Immediately following electroporation, cells were transferred to a 10-cm diameter tissue culture dish with 15 mL of media containing 2 μg/mL doxycycline and handled as previously described.

### Tetramethylrhodamine (TMRE) mitochondrial membrane potential assay

Live, doxycycline-induced UBA1^WT^ or UBA1^M41V^ THP1 model monocytes were treated with either DMSO or 200 μM carbonyl cyanide 3-chlorophenylhydrazone (CCCP) for 10 min at 37 °C in a humidified atmosphere with 5% CO2. Next, TMRE was added to a final concentration of 50 nM to each sample and cells were incubated for 30 min. Cells were then harvested, washed with PBS, and analyzed by flow cytometry by 488 nm laser excitation and detection at 575 nm peak emissions.

### Indo-1AM ratiometric intracellular calcium assay

Live, doxycycline-induced UBA1^WT^ or UBA1^M41V^ THP1 model monocytes were pelleted at 300 × g and resuspended in 1 mL PBS + 10 mM HEPES. To this, 1 mL of loading mix (3 µM Indo-1AM + 0.02% v/v Pluronic + 2.5 mM Probenecid) was added and the mixture incubated for 30 min at 37 °C in the dark. After incubation, cells were pelleted at 300 × g and resuspended in 1 mL assay buffer (PBS + 10 mM HEPES + 2.5 mM Probenecid) and incubated for 30 min at 37 °C in the dark to de-esterify dye. Cells were then analyzed immediately by flow cytometry to assay free versus calcium-bound cytosolic Indo-1AM.

### Cellular fractionation and cytosolic mtDNA quantification

Cellular fractionation and qPCR quantification of cytosolic mtDNA was conducted as previously described (*76*). Briefly, digitonin lysis buffer (50 mM HEPES pH 7.4, 150 mM NaCl, 20 μg/mL digitonin, protease inhibitors), NP-40 lysis buffer (50 mM Tris pH 7.5, 150 mM NaCl, 1 mM EDTA, 1% NP-40 (v/v), and 10% Glycerol (v/v), protease inhibitors), and SDS lysis buffer (20 mM Tris pH 8, 1% SDS (v/v), protease inhibitors) were made fresh. Samples of each THP1 UBA1^WT^ and UBA1^M41V^ macrophages were resuspended with SDS lysis buffer and incubated at 95 °C for 15 min to yield whole cell extract (WCE). Different samples of THP1 UBA1^WT^ and UBA1M41V were gently resuspended in digitonin lysis buffer, lysed for 9 min at 4 °C, and centrifuged 950 × g for 5 min at 4 °C. The resultant supernatant was collected to harvest the cytosolic fraction. The remaining pellet was resuspended in 500 μL of NP-40 lysis buffer, incubated on ice for 10 min, and then spin at ∼21,000 × g for 10 min at 4 °C. The resultant supernatant was collected to harvest the mitochondrial fraction. DNA purification from the isolated aqueous WCE, cytosolic, and mitochondrial fractions was conducted using phenol/chloroform/isoamyl alcohol precipitation followed by ethanol wash and resuspension of the DNA pellet in nuclease free water. Quantification of mtDNA in cellular fractions was performed using TaqMan Universal PCR Master Mix (Applied Biosystems, 4304437) on a QuantStudio 5 Real-Time PCR System per the manufacturer’s directions. A probe targeting KCNJ10 (Applied Biosystems, Hs01922935_s1) was used as a nuclear DNA control in conjunction with two mtDNA probes targeting MTDN1 (Applied Biosystems, Hs02596873_s1) and Mt7s D-Loop (Applied Biosystems, Hs02596861_s1).

### Xenotransplantation of NSG-S mice with base edited HSPCs

For each mouse, 1.0×10^6^ of base edited HSPCs were injected retro-orbitally 24 h after editing into sublethally irradiated (single dose 250 cGy 3 h prior to transplant) NSG-S mice. Mice were all male aged 4-8 weeks of age and randomly distributed to each experimental group. Non-injected HSPCs were set aside for ddPCR to calculate initial UBA1 and STING variant allele frequency at week 0. Human CD45+ cell engraftment was monitored by serial collection of blood at week 4 and at the end of the experiment 7 weeks after transplantation by flow cytometry. At the end of the experiment, bone marrow cells were extracted and purified by AutoMACs using human CD45 microbeads (Miltenyi). These cells were analyzed by scRNA-seq and ddPCR to assess endpoint variant allele frequencies.

## Supporting information

Magaziner_2026_Supplement

## ACKNOWLEDGEMENTS

We thank Alan Schwartz (Washington University in St. Louis) for CHO cells, Defne Ercelen (New York University) for transcriptomic analysis, Mei-Kay Wong (New York University) for clinical data. We would like to thank patients and their families for supporting our research along with the VEXAS Foundation.

## FUNDING

National Institutes of Health grant R00AR078205 (DBB) National Institutes of Health grant R01AR083407 (DBB)Department of Defense: Bone Marrow Failure Research Program Idea Development Award HT9425-23-1-0507 (DBB) The Edward P. Evans Foundation and Gabrielle’s Angel Foundation (DBB) National Institutes of Health grant F30HL170731 (SJM) National Institutes of Health grant GM136542 (SJM) NYU Grossman School of Medicine Medical Scientist Training Program (SJM) National Institutes of Health grant 5TL1TR001447 (BM) National Institutes of Dental and Craniofacial Research ZIA DE000749 (AW, JCC, PZ, ME, JB), ZIC DE000744 (DT), ZIA DE00075 (YW) NationalHeart Lung Blood Institute ZIC HL005906 (ZAS, VB)

## AUTHOR CONTRIBUTION

SJM designed, performed, and analyzed most experiments and helped write the manuscript; JCC designed, performed, and analyzed proteomic experiments and in vitro E1-E2 transhiolation assays, BM analyzed RNAseq and proteomic datasets; PZ performed and analyzed light and fluorescence microscopy experiments, AWa performed cell culture experiments, JH performed experiments with base editing, JCB designed, performed and analyzed HSC experiments, MS analyzed HSC experiments, performed in vitro E1-E2 transthiolation assays; JB established cell culture conditions and machine learning algorithms to quantify vacuoles, RM analyzed single cell RNA seq data, PHW analyzed immunofluoresence, DR performed patient sample experiments and purifications, YW performed MS analysis; DTT supervised, performed, and analyzed light and fluorescence microscopy experiments; ZAS and VB performed and analyzed TEM and FIB-SEM experiments; TL supervised immunofluorescence experiments, KVR supervised transcriptomic and proteomic analyses; IA supervised HSC experiments, DAL supervised scRNA seq experiments, AWe and DBB conceived the study, supervised experiments, and wrote the manuscript. All authors reviewed and edited the manuscript.

## DECLARATION OF INTERESTS

The authors declare that they have no competing interests.

## Notes

### Competing Interest Statement

DAL has served as a consultant for Abbvie, AstraZeneca and Illumina and is on the Scientific Advisory Board of Mission Bio, Pangea, Alethiomics, and C2i Genomics; and has also received prior research funding from BMS, 10x Genomics, Ultima Genomics, and Illumina unrelated to the current manuscript. DBB has served as a consultant for Sobi, Novartis, GSK and Alexion Pharmaceuticals.

