## Supplementary material for "Cell autonomous inflammation in VEXAS is mediated by cGAS-STING": Magaziner_2026_Supplement

**The PDF file includes:**

Supplementary Figs. 1 to 20
Captions for Supplementary Tables 1 to 10
Captions for Supplementary Movies 1 to 2

**Other Supplementary Materials for this manuscript include the following:**

Supplementary Tables 1 to 10
Supplementary Movies 1 to 2

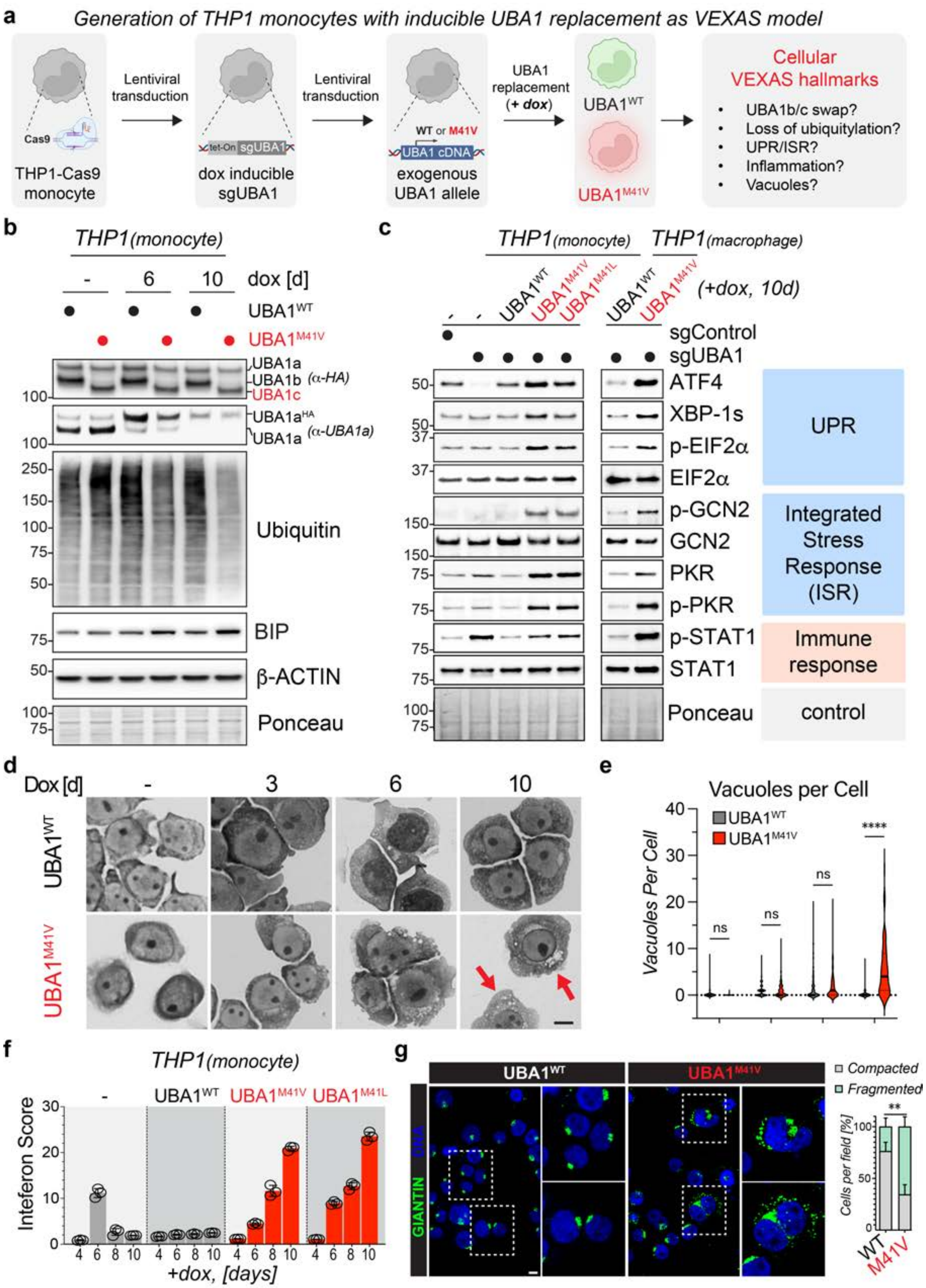


**Supplementary Figure 1. An inducible THP1 model recapitulates cellular VEXAS hallmarks.**
**(a)** Schematic representation of THP1 inducible cell line stably expressing Cas9, a doxycycline

inducible sgRNA targeting *UBA1*, expressing either *UBA1$^{WT}$* or *UBA1$^{M41V}$* cDNA resistant to sgRNA targeting. **(b)** Validation of loss of endogenous UBA1 and rescue with either *UBA1$^{WT}$* and *UBA1$^{M41V}$* cDNA leading to a loss of ubiquitylation and accumulation of ER stress, as monitored by immunoblotting of indicated THP1 lysates with indicated antibodies. **(c)** Loss of cytoplasmic UBA1 function in THP1 model cells either as monocytes or macrophages leads to UPR and ISR activation and inflammatory signaling, recapitulating VEXAS. *UBA1$^{WT}$* and *UBA1$^{M41V}$* THP1 cells were treated with doxycline for 10d and subjected to immunoblotting using the indicated antibodies. Results are representative of n ≥ 3 biological replicates. **(d)** THP1 *UBA1$^{M41V}$* monocytes exhibit cytoplasmic vacuole formation, as assessed via Wright-Giemsa staining after dox treatment for indicated time periods. **(e)** Vacuoles were quantified via a machine learning algorithm (see methods). n = 3 biological replicates, **** = $p < 0.0001$, student's t-test. **(f)** THP1 VEXAS model monocytes exhibit type I interferon signatures which increase over time. UBA1$^{WT}$ and *UBA1$^{M41V/L}$* THP1 cells were treated with dox for indicated time periods and analyzed by qPCR for Interferon Score by averaging the fold change of 6-hallmark ISGs (*IFI44L, IFI27, IFIT1, ISG15, RSAD2,* and *SIGLEC1*). n ≥ 3 biological replicates, error bars = s.e.m., **** = $p < 0.0001$, student's t-test. **(g)** Loss of cytoplasmic UBA1 function in THP1 model cells leads to Golgi fragmentation, validating the results of the transcriptomic and proteomic analyses. *UBA1$^{WT}$* and *UBA1$^{M41V}$* THP1 cells were treated with dox for 10 d and subjected to anti-GIANTIN immunofluorescence analysis followed by quantification of Golgi morphology. n = 3 biological replicates, ** = $p < 0.01$, error bar = s.e.m, student's t-test. Scale bar = 10 mm.

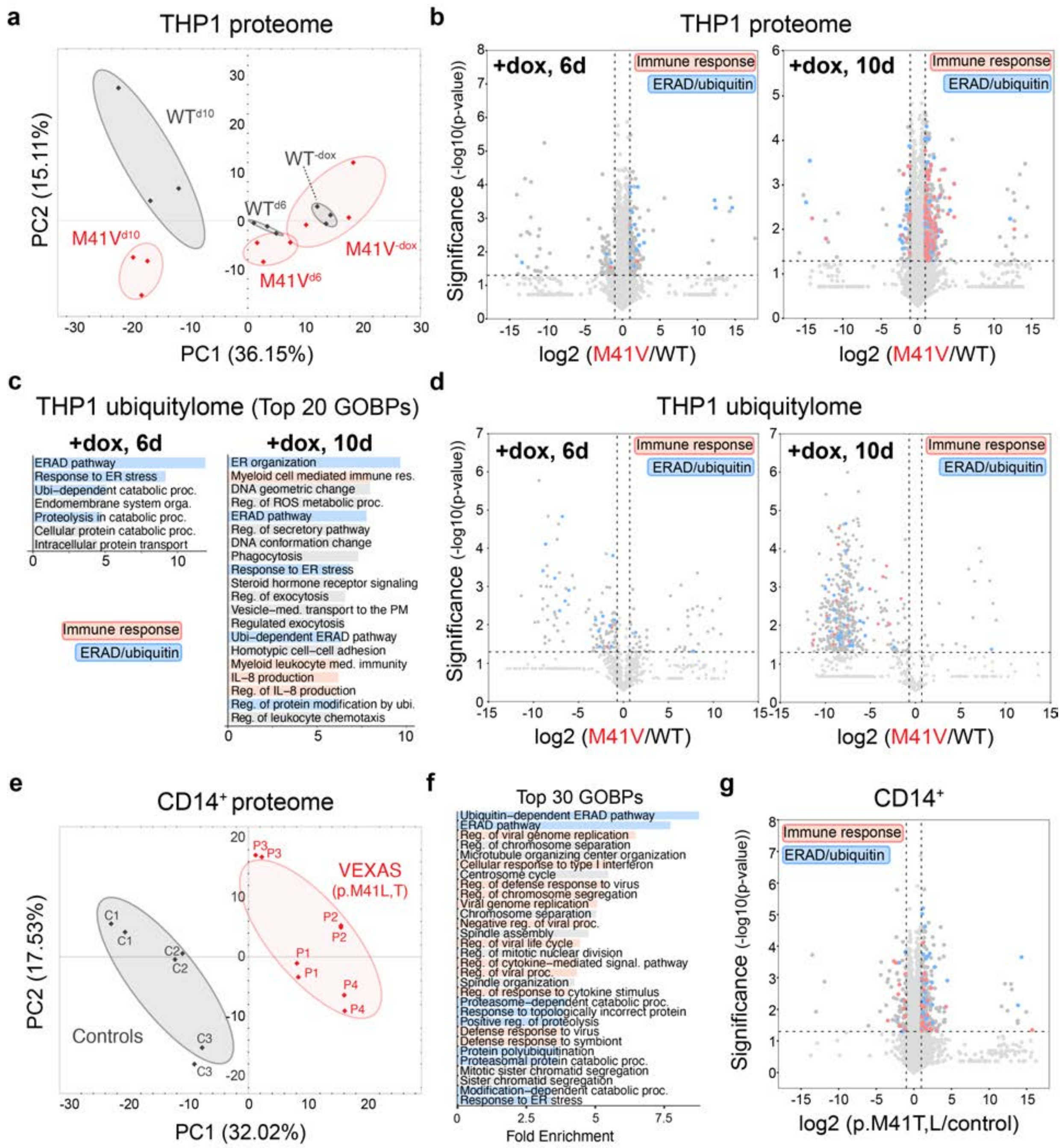


**Supplementary Figure 2. Proteomic analyses of THP1 monocytes and patient derived CD14+ myeloid cells identify ERAD dysregulation as an early VEXAS signal preceding inflammation.** **(a)** Induction of UBA1 replacement in the THP1 VEXAS model causes progressive differences between the proteomes of *UBA1^WT^* and *UBA1^M41V^* cells. THP1 model cells were treated with doxycycline for indicated time periods and total proteomes were determined by DIA mass spectrometry followed by principal component (PC) analysis. **(b)** THP1 VEXAS model cells exhibit dysregulated ERAD prior to immune response activation. The volcano plots depict differentially expressed proteins (DEPs) of *UBA1^WT^* and *UBA1^M41V^* THP1 monocytes 6 days post dox induction (right) and 10 days post dox induction (left). DEPs related to immune responses and ERAD are highlighted. **(c)** Similar to the total proteome, the ubiquitylome of THP1 VEXAS model cells exhibit dysregulated ERAD prior to immune response activation. THP1 model cells were treated with doxycycline for indicated time periods and were subjected to diGly proteomics followed by GOBP analysis of significantly differentially ubiquitylated proteins ($p<0.05$, $-0.7 < \log_2(\text{VEXAS/control}) > 0.7$). Up to top 20 significantly enriched GOBPs (FDR < 0.05) ranked according to fold enrichment score are shown. **(d)** The volcano plots depict differentially ubiquitylated proteins of *UBA1*$^{WT}$ and *UBA1*$^{M41V}$ THP1 monocytes 6 days post dox induction (left) and 10 days post dox induction (right). **(e)** The proteomes of VEXAS patient derived CD14$^+$ monocytes are different from those of healthy donor controls. CD14$^+$ monocytes were isolated from VEXAS patients or healthy donors and total proteomes were determined by DIA proteomics followed by principal component (PC) analysis. **(f)** CD14$^+$ monocytes from VEXAS patients exhibit dysregulated ERAD and immune responses, as revealed by GOBP analysis of significantly DEPs ($p<0.05$, $-0.7 < \log_2(\text{VEXAS/control}) > 0.7$). Top 30 significantly enriched GOBPs (FDR < 0.05) ranked according to fold-enrichment score are shown. **(g)** Volcano plot depicting DEPs of healthy donor and VEXAS CD14$^+$ monocytes. DEPs related to immune responses and ERAD are highlighted.

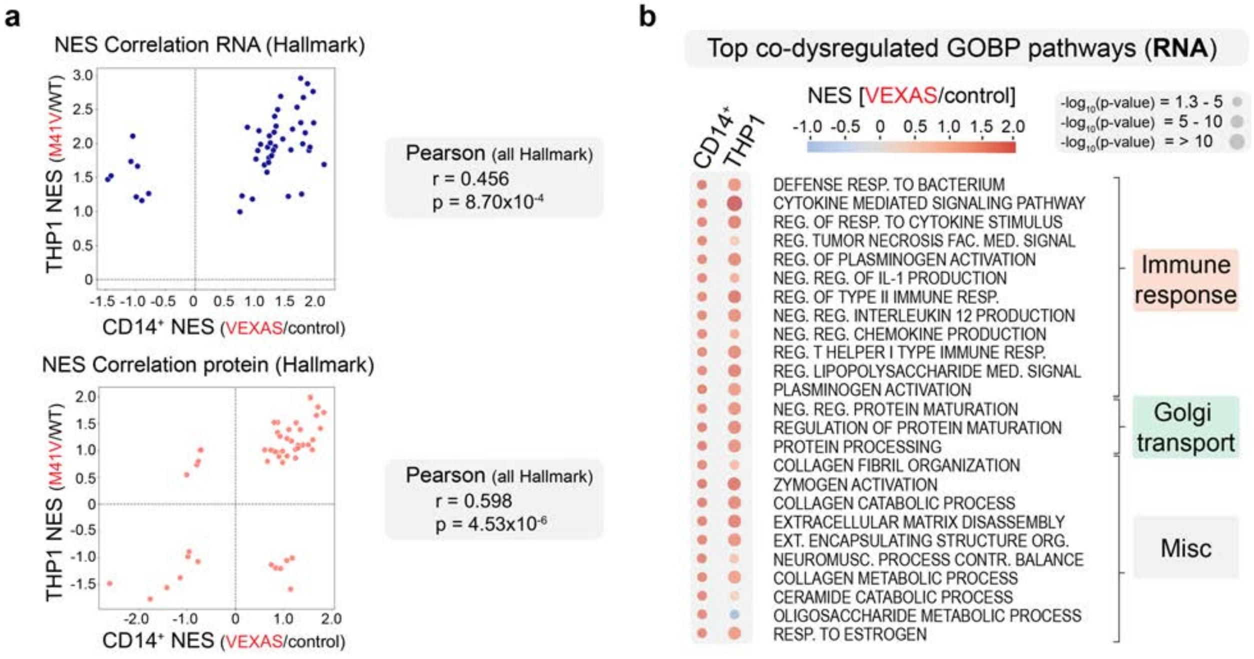


**Supplementary Figure 3**. **THP1 monocyte model and patient-derived CD14+ VEXAS cells exhibit highly correlated dysregulated GOBPs. (a)** Proteomic studies correlate more closely between VEXAS model THP1 and patient-derived CD14$^+$ VEXAS cells than transcriptomic analyses. Plots depict the Pearson correlations of normalized enrichment scores (NES) of hallmark pathways for RNA (top) and protein (bottom). **(b)** Dot plots show gene set enrichment scores for the top 25 dysregulated GOBPs as identified by transcriptomic comparison of patient-derived control and VEXAS monocytes (CD14$^+$). For each CD14$^+$ top dysregulated GOBP, the mean normalized enrichment score (NES) and the Benjamini-Hochberg-adjusted p-values are shown for the two indicated comparisons (CD14$^+$ RNA, THP1 RNA). n=3 biological replicates for each condition.

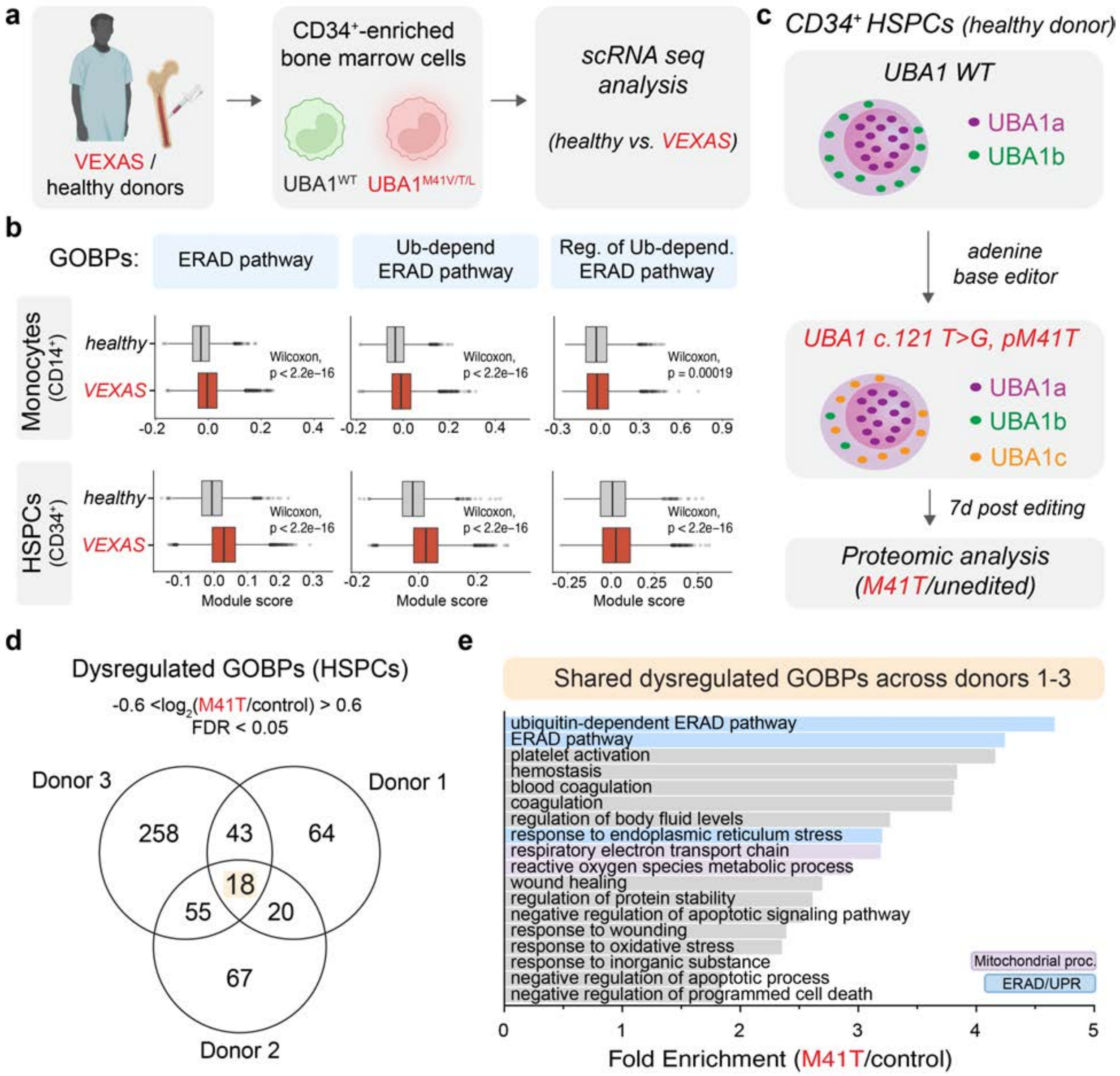


**Supplementary Figure 4**. **scRNA seq of patient cells and bulk proteomics of *UBA1$^{M41T}$* base edited CD34+ HSPCs identify ERAD as a prominently dysregulated pathway. (a)** Schematic representation of samples used for scRNA sequencing from bone marrow from VEXAS patients or healthy controls. **(b)** ERAD is dysregulated in monocytes and hematopoietic stem cells (HSPCs) from patients with VEXAS as compared to healthy controls, as demonstrated by scRNA seq analysis. Graphs depict GOBP analyses between controls and patients with VEXAS in monocytes (CD14$^+$, top row) and HSPCs (CD34$^+$, bottom row). **(c)** Schematic of base editing approach to generate *UBA1$^{M41T}$* HSPCs *in vitro*. **(d)** Venn diagram demonstrating overlap in dysregulated GOBPs of total proteomes of three healthy donor HSPCs, each analyzed as a comparison between unedited and *UBA1$^{M41T}$* cells. All three HSPCs studies demonstrated > 85% editing of *UBA1*. **(e)** ERAD is the top dysregulated pathway across *UBA1$^{M41T}$* mutant HSCs proteomes. Graph depicts the 18 shared significantly dysregulated GOBPs (FDR < 0.05) upon *UBA1$^{M41T}$* editing in HSPCs ranked according to the fold enrichment score.

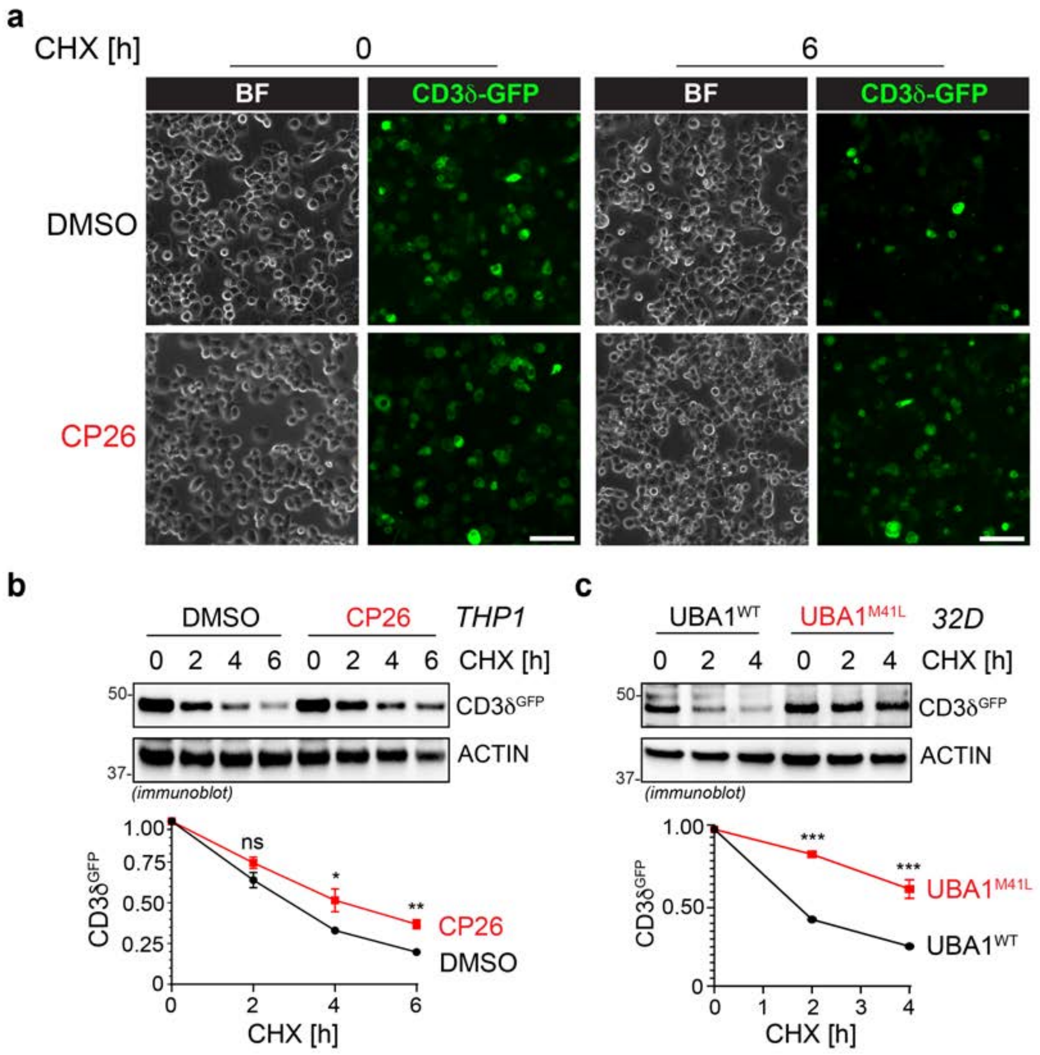


**Supplementary Figure 5. CD3d-GFP is an ERAD model substrate and can be used to determine ERAD dysfunction in 32D model cells. (a)** CD3d-GFP is degraded via ERAD in THP1 cells. THP1 cells expressing the ERAD model substrate CD3d-GFP were treated with cycloheximide (CHX) in the absence and presence of the ERAD inhibitor CP26 for indicated time periods followed by brightfield (BF) and GFP fluorescent microscopy. Scale bar = 50 mm. **(b)** CD3d-GFP expressing THP1 cells treated with cycloheximide (CHX) for indicated time periods were subjected to immunoblotting and CD3d-GFP quantifications were normalized to ACTIN. n ≥ 3 biological replicates, error bars = s.e.m., * = $p < 0.05$, ** = $p < 0.01$, *** = $p < 0.001$, student's t-test. **(c)** ERAD impairment is present in murine models of VEXAS. $Uba1^{WT}$ and *$Uba1^{M41L}$* 32D cells expressing the ERAD reporter, CD3d-GFP, were treated with cycloheximide (CHX) for indicated time periods, and subjected to immunoblotting with indicated antibodies. CD3d-GFP quantifications were normalized to ACTIN. n ≥ 3 biological replicates, error bars = s.e.m., * = $p < 0.05$, ** = $p < 0.01$, *** = $p < 0.001$, student’s t-test.

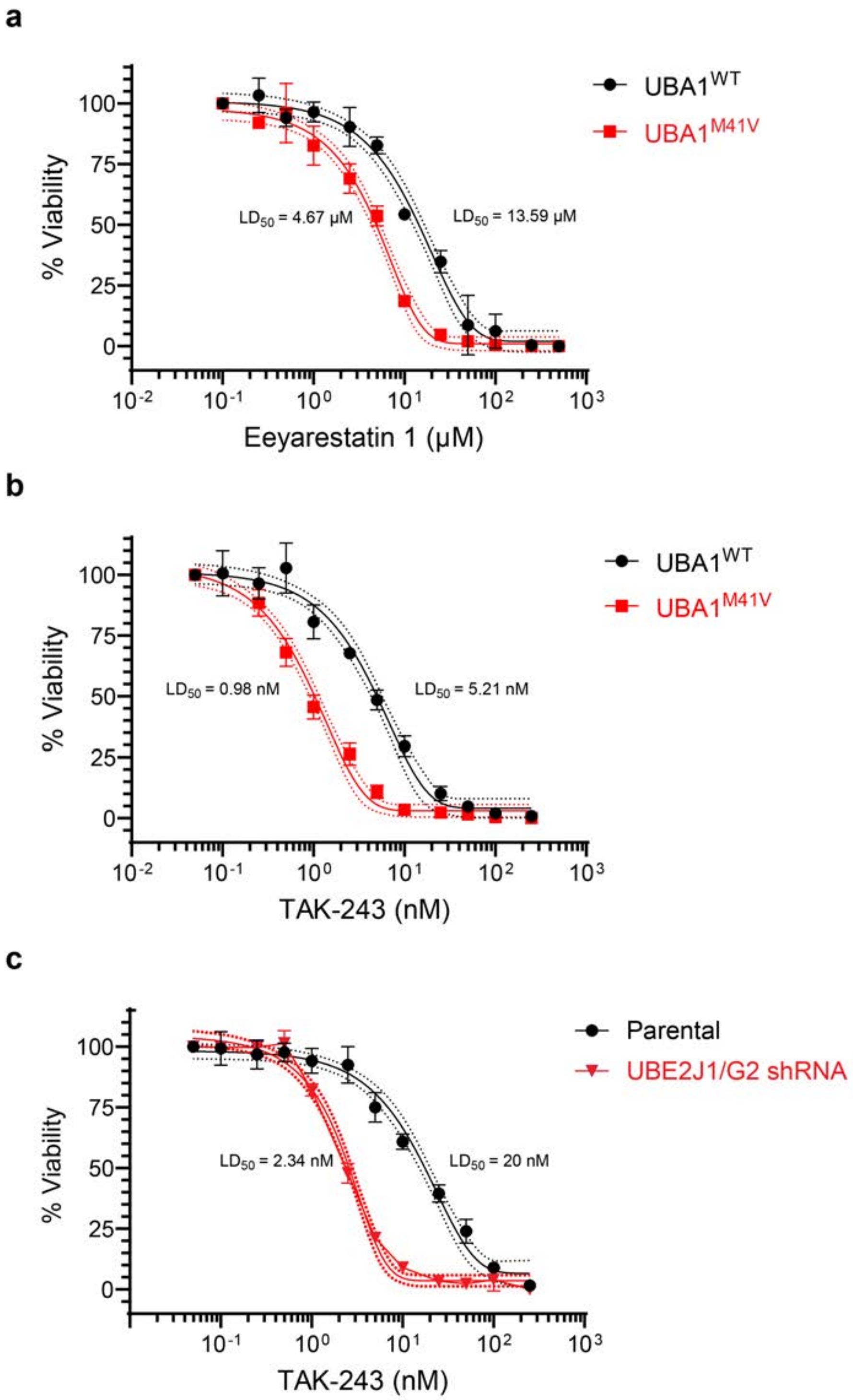


**Supplementary Figure 6. ERAD inhibition induces synthetic lethality in THP1 cells. (a)** VEXAS model THP1 cells demonstrate increased sensitivity to ERAD inhibition. Viability is measured at increasing concentrations of ERAD inhibition (Eeyarestatin I) between $UBA1^{WT}$ and $UBA1^{M41V}$ THP1 cells. $LD_{50}$ shown with standard error of the mean plotted from $n \geq 3$ biological replicates. **(b)** VEXAS model THP1 cells demonstrate increased sensitivity to UBA1 inhibition. Viability is measured at increasing concentrations of UBA1 inhibition (TAK-243) between $UBA1^{WT}$

and *UBA1*$^{M41V}$ THP1 cells. $LD_{50}$ shown with standard error of the mean plotted from n ≥ 3 biological replicates. **(c)** Genetic inhibition of ERAD results in increased sensitivity to UBA1 inhibition as compared to control THP1 cells. Viability is measured at increasing concentrations of UBA1 inhibition (TAK-243) between control and UBE2J1/G2 shRNA-depleted THP1. $LD_{50}$ shown with standard error of the mean plotted from n ≥ 3 biological replicates.

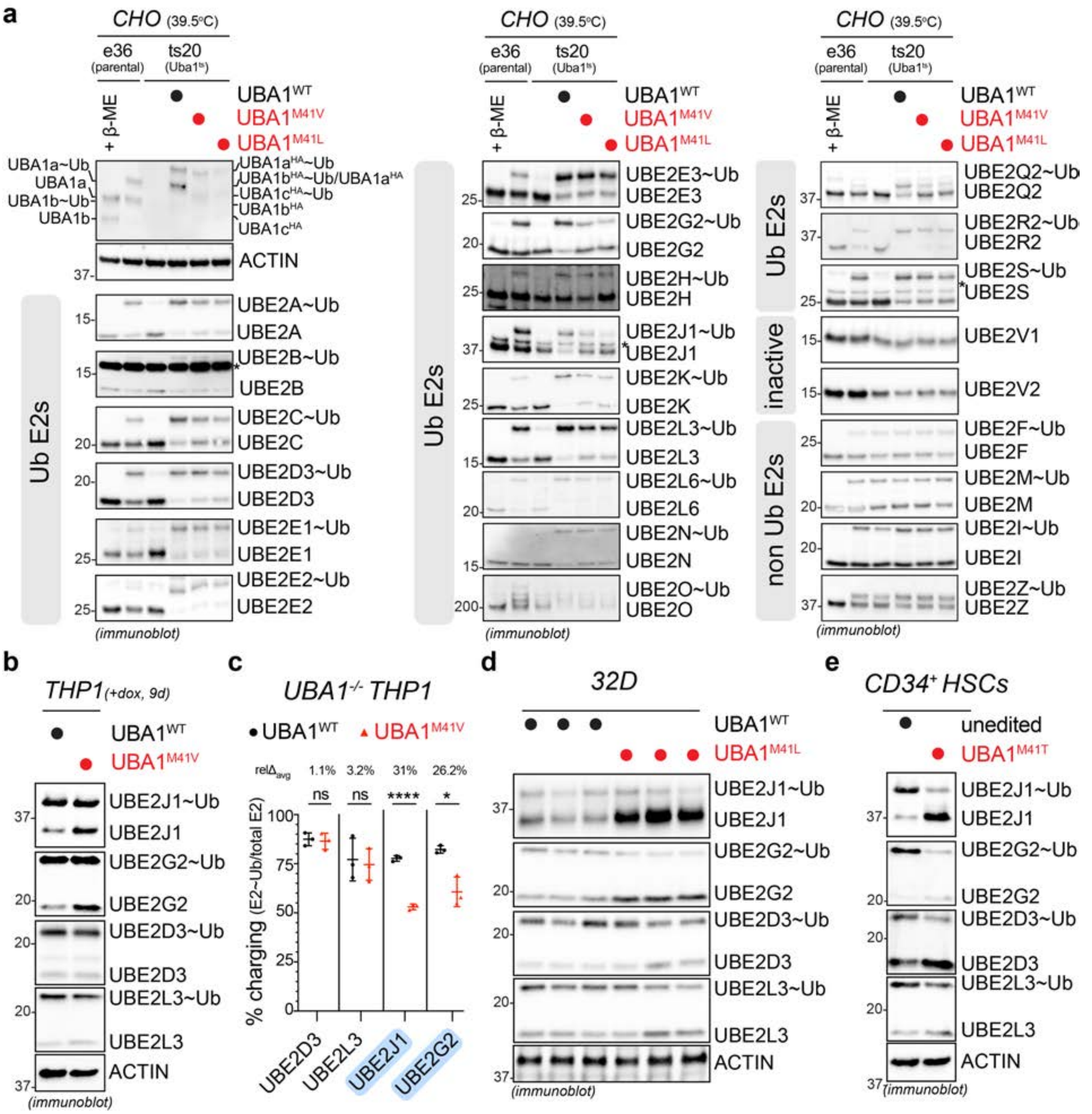

**Supplementary Figure 7. The preferential loss of ubiquitin charging of ERAD E2s in the setting of VEXAS is conserved across cell types. (a)** Loss of cytoplasmic UBA1 activity causes a preferential loss of ubiquitin charging of the ERAD-associated E2 enzymes UBE2J1 and UBE2G2. Chinese hamster ovary (CHO) cells with a temperature sensitive Uba1 allele (ts20) and complemented with human *UBA1^WT^*, *UBA1^M41V^*, or *UBA1^M41L^* were incubated at the restrictive temperature and subjected to non-reducing immunoblotting using antibodies against the 28 detectable E2s in CHO cells. Representative immunoblots of the quantifications shown in the heat map in Figure 2C, for which the ubiquitin charging status of each E2 enzyme was calculated based on immunoblot signal of ubiquitin-charged over total E2, normalized to *UBA1^WT^*. **(b)** Confirmation of the specific E2 charging defect of ERAD-associated UBE2J1 and UBE2G2 in THP1 VEXAS model cells. *UBA1^WT^* and *UBA1^M41V^* THP1 cells were treated with doxycycline for 9 days, subjected to non-reducing immunoblotting using the indicated antibodies. Representative immunoblots of n = 3 biological replicates are shown. Of note, in *UBA1*$^{M41V}$ THP1 cells, the total amount of UBE2J1 and UBE2G2 are increased, likely as a cellular compensation mechanism. **(c)** Quantification of the ubiquitin charging status for each E2 (E2~Ub/ total E2) depicted in panel B. Rel$\Delta_{avg}$ denotes internal % charging change between between *UBA1*$^{WT}$ and *UBA1*$^{M41V}$ for each E2. n = 3 biological replicates, error bar = s.e.m, * = p < 0.05, **** = p < 0.0001, student's t-test. **(d)** Confirmation of the specific E2 charging defect of ERAD-associated UBE2J1 and UBE2G2 in 32D VEXAS model cells. Three independent single-cell selected *Uba1*$^{WT}$ and *Uba1*$^{M41L}$ 32D clones were subjected to non-reducing immunoblot analysis using the indicated antibodies. **(e)** Confirmation of the specific E2 charging defect of ERAD-associated UBE2J1 and UBE2G2 in base edited *UBA1*$^{M41T}$ mutant HSCs. *UBA1*$^{M41T}$ mutant and unedited HSCs were subjected to non-reducing immunoblotting using indicated antibodies. Representative immunoblots of n = 3 biological replicates are shown.

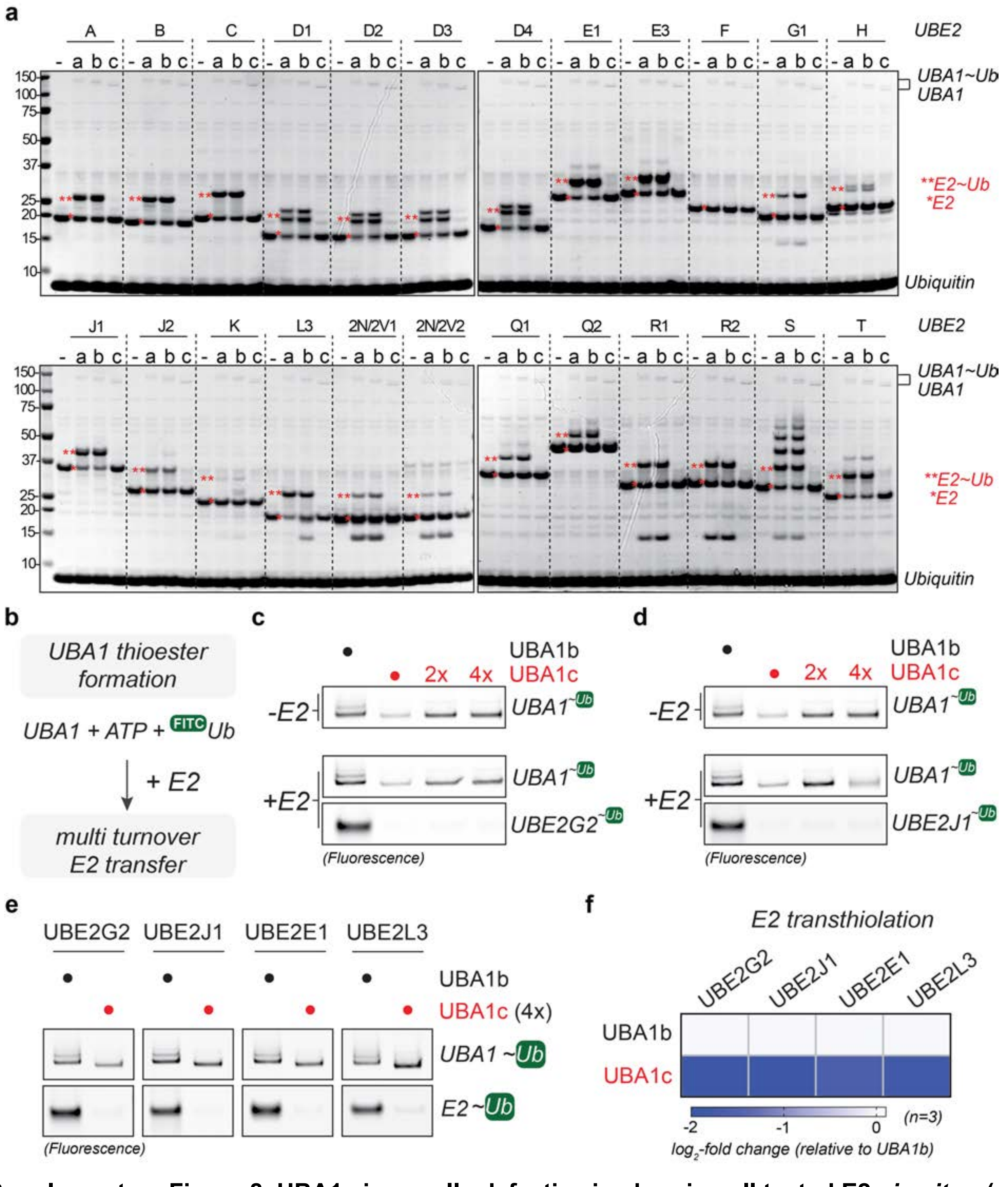


**Supplementary Figure 8. UBA1c is equally defective in charging all tested E2s *in vitro*. (a)** *In vitro* E2 charging with recombinant UBA1 isoforms demonstrates that UBA1c is equally defective at charging all 24 tested E2 enzymes. Recombinant E2 charging (denoted with *) was measured in the presence of no E1 enzyme, UBA1a, UBA1b, or UBA1c across 24 E2 enzymes tested. **(b)** UBA1c is impaired in E1 and E2 charging *in vitro*. Ubiquitin conjugation is measured by the transfer of FITC-labeled ubiquitin in the presence of no E2 or **(c)** UBE2G2 or **(d)** UBE2J1. Increasing concentrations of UBA1c were utilized in red. **(e)** UBA1c is unable to charge E2 enzymes i*n vitro*. E2 charging was measured using FITC labeled ubiquitin with a subset of E2 enzymes in the presence of UBA1b or UBA1c (at 4x higher concentration to ensure maximal ubiquitin loading of UBA1c that is comparable to that of UBA1b). **(f)** Quantification of the charging defect from panel E, quantified from n = 3 biological replicates.

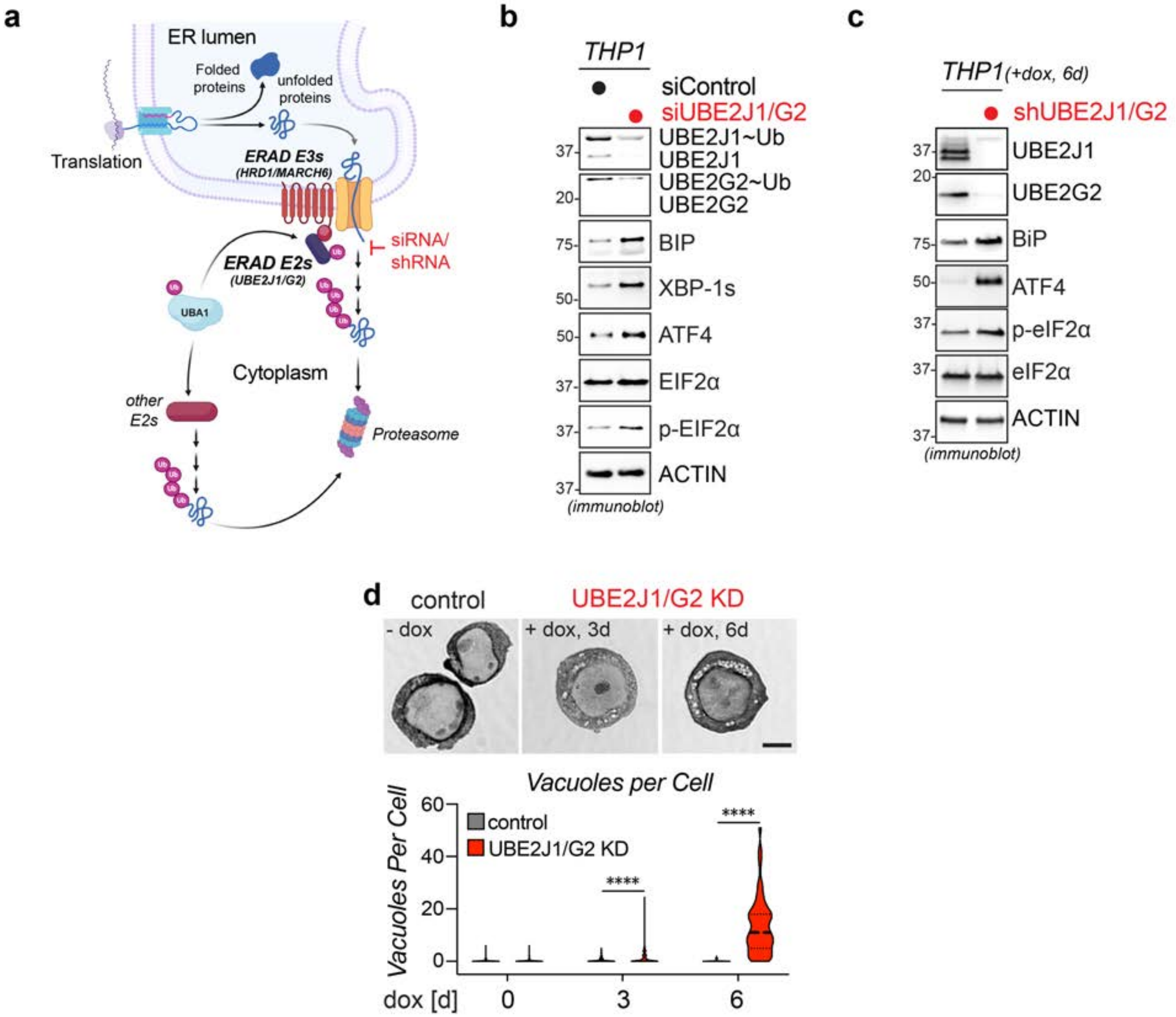


**Supplementary Figure 9. UBE2J1/UBE2G2 depletion in THP1 cells causes ER stress, UPR activation, and vacuole formation** (a) Schematic representation of the ERAD pathway and genetic depletion approaches (generated with Biorender). **(b)** Loss of UBE2J1 and UBE2G2 leads to activation of UPR and ISR. THP1 cells were treated with siRNA for 3 days and then subjected to immunoblotting using the indicated antibodies. **(c)** Loss of UBE2J1 and UBE2G2 leads to activation of UPR and ISR. THP1 cells were infected with inducible lentiviral shRNA for UBE2J1 and UBE2G2 and induced with doxycycline for 6 days followed by immunoblotting using the indicated antibodies. **(d)** ERAD inhibition leads to cytoplasmic vacuole formation, as assessed via Wright-Giemsa staining of THP1 control or conditionally UBE2J1/UBE2G2-depleted cells. Vacuoles were quantified via a machine learning algorithm (see methods). n = 3 biological replicates, **** = $p < 0.001$, student's t-test.

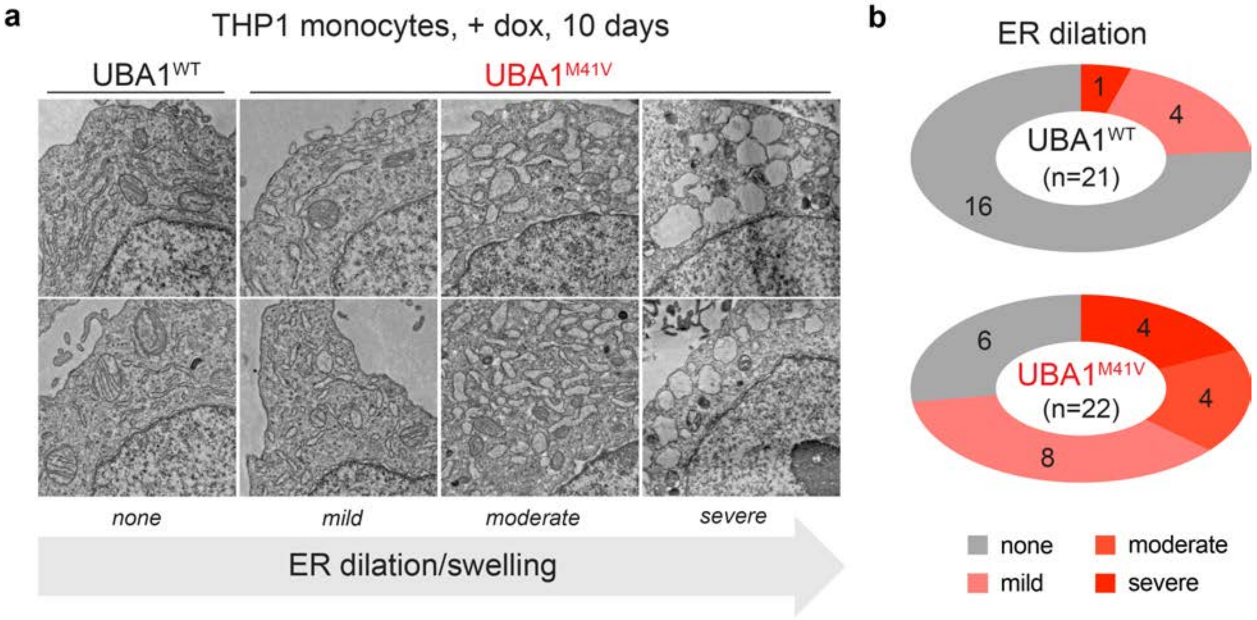


**Supplementary Figure 10. Most THP1 *UBA1$^{M41V}$* monocytes exhibit dilated ER cisterna filled with proteins.** *UBA1$^{WT}$* and *UBA1$^{M41V}$* THP1 monocytes induced with doxycycline for 10 days and imaged by transmission electron microscopy (TEM) were analyzed for ER morphology. (a) *UBA1$^{M41V}$* monocytes display a range of swollen ER defects, as revealed by transmission electron microscopy (TEM). Representative TEM images from the different classifications of ER dilation and swelling are shown. Scale bar = 600 nm. **(b)** Quantification of classes of ER dilation between *UBA1$^{WT}$* and *UBA1$^{M41V}$* THP1 cells. n > 20 cells per condition.

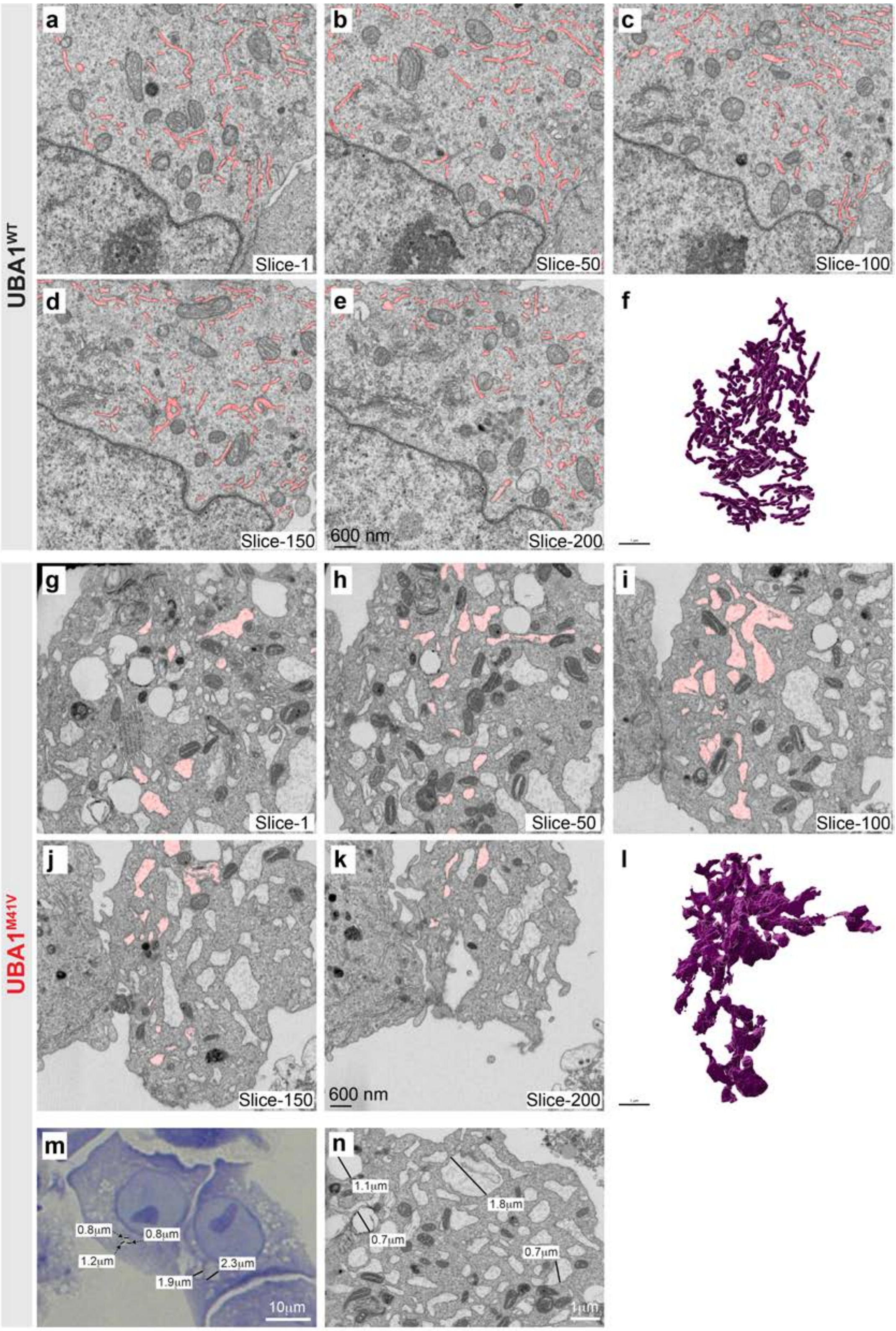


**Supplementary Figure 11. TEM and FIB-SEM analysis reveal that THP1 *UBA1$^{M41V}$* monocytes exhibit massively dilated ER consistent in size with vacuoles observed in Wright-Giemsa stains.** *UBA1$^{WT}$* and *UBA1$^{M41V}$* THP1 monocytes were treated with doxycycline for 10 days, fixed, and subjected to FIB-SEM analysis. **(a-e)** Representative FIB-SEM z-slices of a *UBA1*$^{WT}$ THP1 monocyte at different intervals showing the interconnected morphology of ER. The grayscale FIB-SEM images are overlaid with segmentation mask pseudo-colored in red highlighting the ER network. Scale bar = 600 nm. **(f)** A 3D volume rendering of the ER network of the *UBA1*$^{WT}$ THP1 monocyte. This data is also depicted in Fig. 2H and Movie 1. **(g-k)** Representative FIB-SEM z-slices of a *UBA1*$^{M41V}$ THP1 monocyte at different intervals showing the dilated morphology of ER. The grayscale FIB-SEM images are overlaid with segmentation mask pseudo-colored in red highlighting the ER network. Scale bar = 600nm. (**i**) A 3D volume rendering of the ER network of the *UBA1*$^{M41V}$ THP1 monocyte. This data is also depicted in Fig. 2H and Movie 1. **(m-n)** Vacuoles in *UBA1*$^{M41V}$ THP1 monocytes detected by Wright Giemsa staining (panel m) are in size consistent with dilated ER depicted on FIB-SEM image (panel N). *UBA1*$^{M41V}$ THP1 monocytes were treated with dox for 10 d.

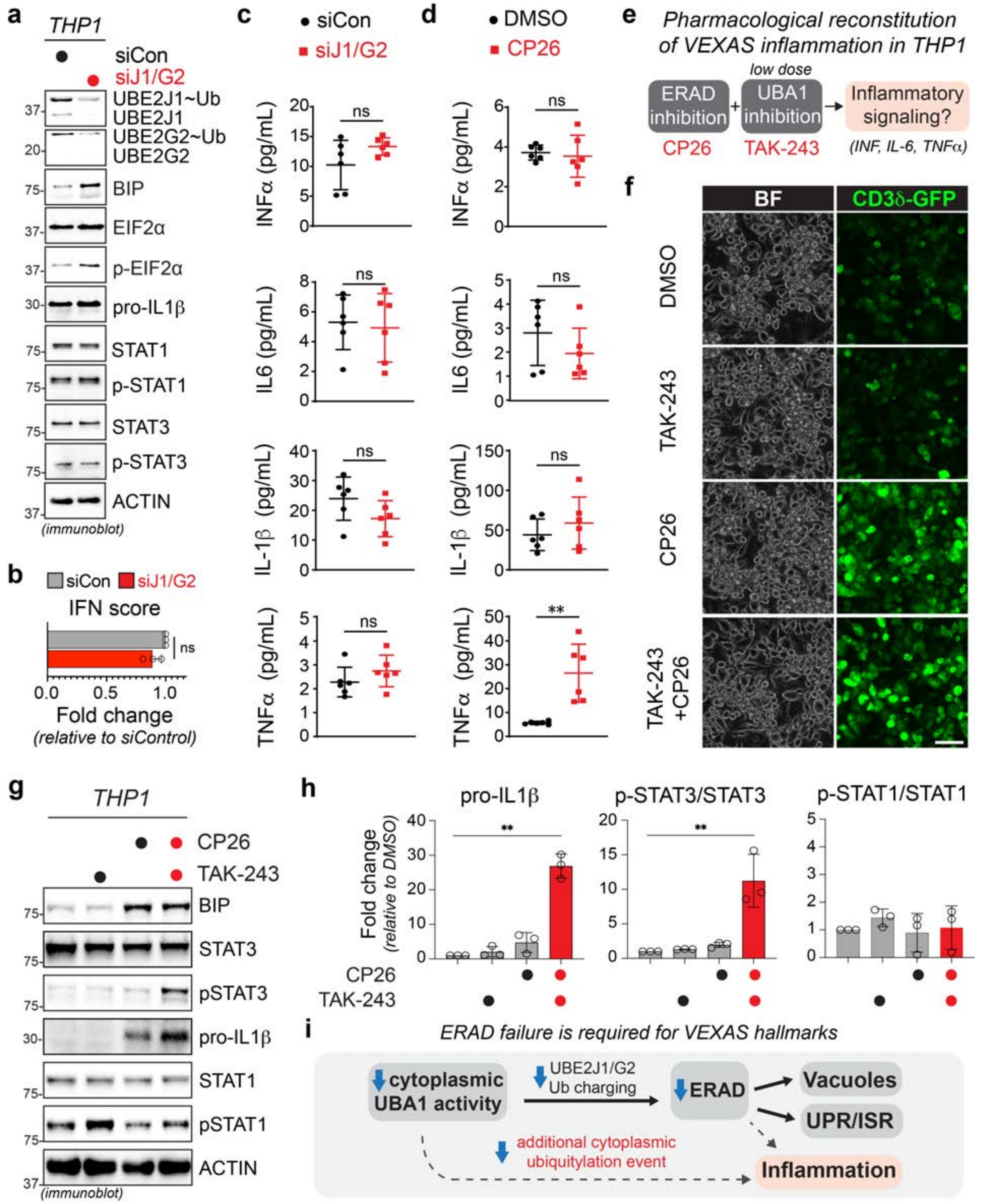


**Supplementary Figure 12. Combing ERAD and low dose UBA1 inhibition can elicit canonical VEXAS inflammatory markers in THP1 cells. (a)** Loss of UBE2J1 and UBE2G2 causes activation of UPR and ISR, but not inflammatory signaling. THP1 macrophages were treated with siRNA for 3 days and then subjected to immunoblotting using the indicated antibodies. This is the same experiment as shown in Fig. S9B, but probing for inflammation markers in addition to UPR/IRS markers. **(b)** Loss of UBE2J1 and UBE2G2 does not increase type I interferon production. THP1 macrophages were treated with indicated siRNA for 3 days and then subjected qPCR analysis for Interferon Score by averaging the fold change of 6-hallmark ISGs (*IFI44L, IFI27, IFIT1, ISG15, RSAD2,* and *SIGLEC1*). n = 3 biological replicates, error bars = s.e.m., ns = not significant, student's t-test. **(c)** Loss of UBE2J1 and UBE2G2 does not cause inflammatory cytokine secretion. THP1 cells were treated with indicated siRNA for 3 days and cell supernatants were analyzed by ELISA to determine the amount of indicated secreted cytokines. n = 3 biological replicates, error bars = s.e.m., ns = not significant, student's t-test. **(d)** ERAD inhibition does not cause VEXAS-associated inflammatory cytokine secretion. THP1 macrophages were treated with DMSO of 1.25 µM CP26 for 2 days and cell supernatants were analyzed by ELISA to determine the amount of indicated secreted cytokines. n = 3 biological replicates, error bars = s.e.m., ns = not significant, ** = $p < 0.01$, student's t-test. **(e)** Schematic explaining the rationale to pharmacologically mimic VEXAS in THP1 cells. **(f)** CD3d-GFP expressing THP1 macrophages, treated with DMSO, 1 nM TAK-243 (UBA1 inhibition), CP26 1.25 µM (ERAD inhibition) or combination treatment for 2 days. Dosing for UBA1 inhibition was based on levels that did not elicit ERAD failure, and dosing for ERAD inhibition was based on lowest dose causing CD3d-GFP accumulation. Scale bar = 50 mm. **(g)** ERAD inhibition is required to elicit UBA1 dependent inflammation in THP1 cells. THP1 macrophages treated with DMSO, CP26, TAK-243, or a combination of both treatments were subjected to immunoblotting using the indicated antibodies. **(h)** Graphs depict quantifications of pro-IL-1β, p-STAT1/STAT1 ratio, and p-STAT3/STAT3 ratio normalized to DMSO treated THP1 macrophages. n = 3 biological replicates, error bar = s.e.m, ** = $p < 0.01$, student's t-test. **(i)** Schematic outlining ERAD failure is required for VEXAS hallmarks including activation of UPR and ISR, vacuole formation, and inflammation.

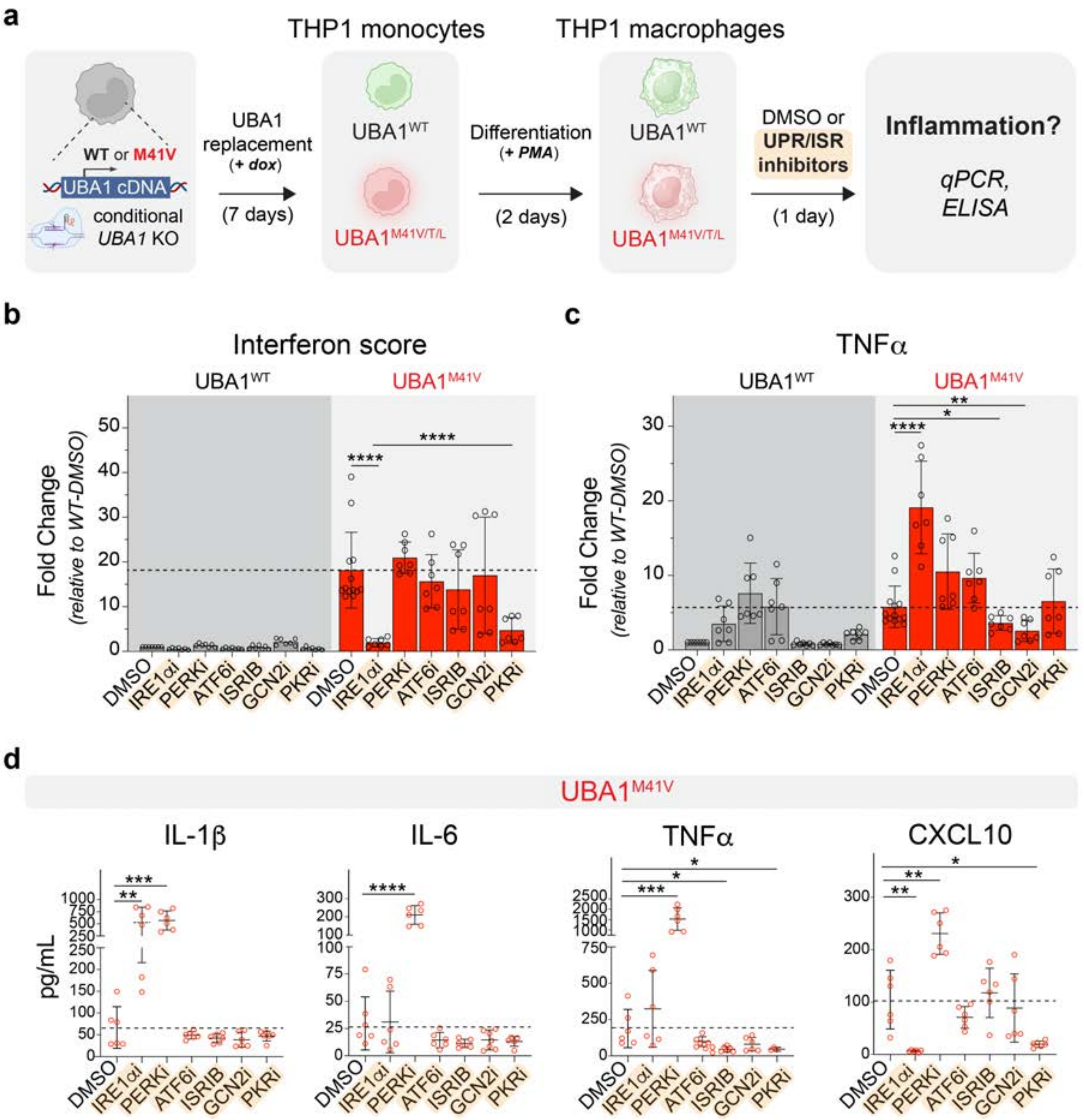


**Supplementary Figure 13. Inhibition of UPR/ISR components in THP1 model cells either does not or only blocks select VEXAS-associated inflammatory pathways while exacerbating others. (a)** Schematic of experimental workflow. *UBA1$^{WT}$* and *UBA1$^{M41V/L}$* THP1 macrophages were treated with dox for 10 d, followed by UPR/ISR inhibition, and analyzed for changes in inflammation. **(b)** Distinct UPR and ISR inhibitors reduce Type I interferon-based inflammation in VEXAS models. qPCR for Interferon Score was taken by averaging the fold change of 6-hallmark ISGs (*IFI44L, IFI27, IFIT1, ISG15, RSAD2,* and *SIGLEC1*). Dotted line represents normalization to DMSO control. n ≥ 3 biological replicates, error bars = s.e.m., **** = $p < 0.0001$, student's t-test. **(c)** Some but not all UPR and ISR inhibitors blunt TNFα. *UBA1$^{WT}$* and *UBA1$^{M41V/L}$*. THP1 cells were treated with dox for 10 d, followed by UPR/ISR inhibition and analyzed by qPCR for TNFα. Notably, some inhibitors show contradictory (e.g. IRE1ai) or independent (e.g. ISRIB and GCN2i) on TNFα expression. **(d)** Commensurate with qPCR results, no UPR or ISR inhibitor reduces multiple hallmark cytokines from VEXAS with some showing contradictory increases across different axes of inflammation. *UBA1$^{M41V}$* THP1 macrophages were treated with dox for 10d, followed by UPR/ISR inhibition and analyzed by ELISA. Cell supernatants were subjected to ELISA to measure secreted protein concentrations of IL-6, TNFα, IL-1β, and CXCL10. n = 3 biological replicates, error bar = s.e.m, * = $p < 0.05$, ** = $p < 0.01$, *** = $p < 0.001$, **** = $p < 0.0001$, student's t-test.

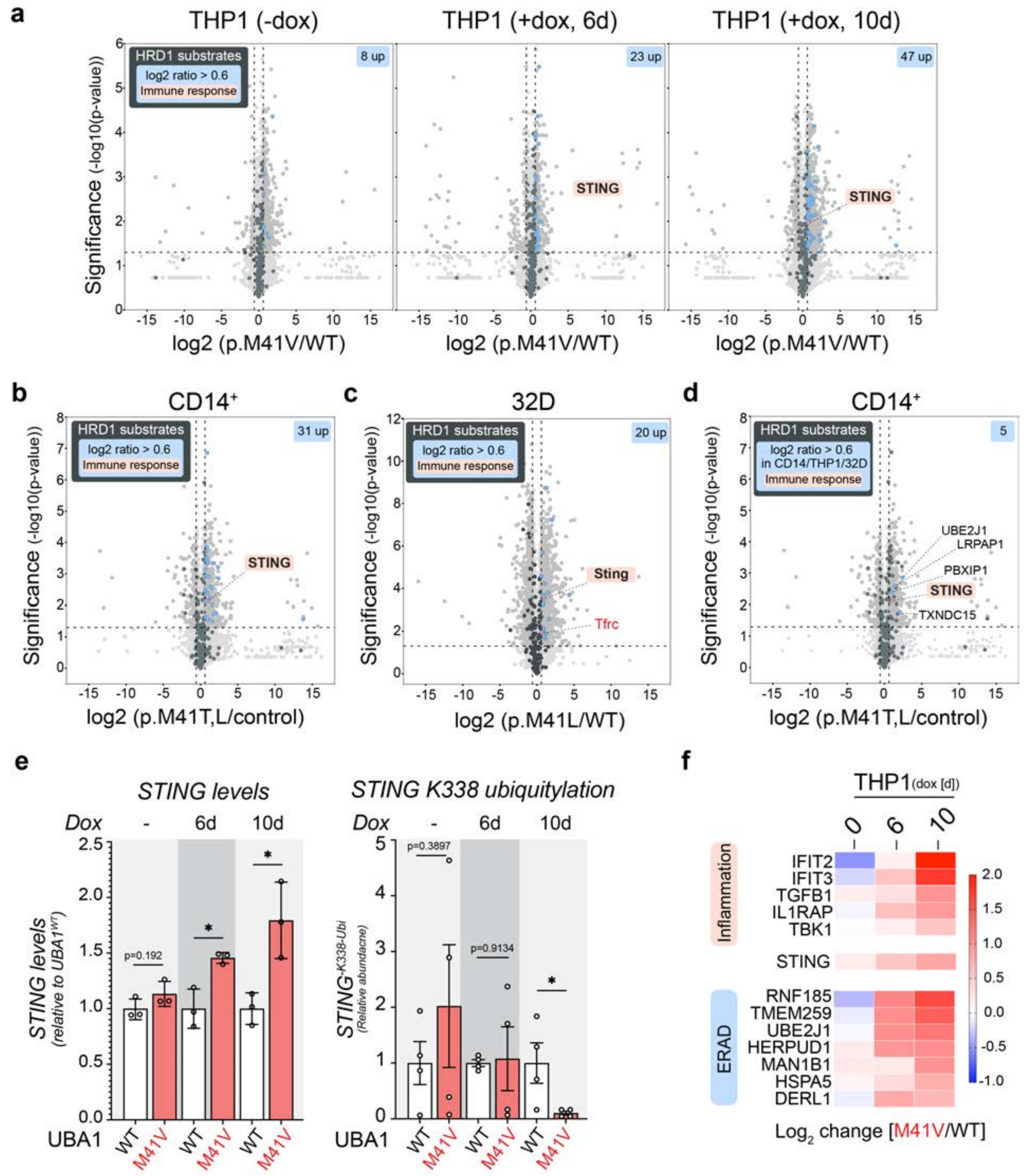


**Supplementary Figure 14. STING is the only immune response related ERAD substrate that accumulates in model and patient-derived VEXAS cells.** DIA proteomics datasets were analyzed to identify HRD1/ERAD substrates amongst differentially expressed proteins (DEPs). In all volcano plots, previously reported HRD1/ERAD substrates(see Supplementary Table 6) that were detected are highlighted in dark grey. Significantly upregulated HRD1/ERAD substrates are highlighted in blue, with immune-response related proteins highlighted in red. **(a)** Induction of UBA1 replacement in the THP1 VEXAS model causes progressive accumulation of various ERAD substrates in U*BA1*$^{M41V}$ cells, with STING being the only ERAD substrate associated with immune responses. The volcano plots depict DEPs between *$UBA1^{WT}$* and *$UBA1^{M41V}$* THP1 monocytes after 6 or 10 days of doxycycline induction. **(b)** VEXAS patient monocytes accumulate many ERAD substrates with STING being the only one associated with immune responses. $CD14^+$ monocytes were isolated from VEXAS patients (UBA1p.M41T/L) or healthy donors (control. The volcano plot depicts DEPs between healthy donor and VEXAS patient cells. **(c)** 32D genetic VEXAS model cells accumulate ERAD substrates, with Sting and Tfrc being the only ones associated with immune responses. The volcano plot depicts differentially expressed proteins (DEPs) between *$Uba1^{WT}$* and *$Uba1^{M41L}$* 32D cells. **(d)** STING is the only immune response related ERAD substrate across all VEXAS models and patient cells. The volcano plot depicts DEPs of healthy donor and VEXAS $CD14^+$ monocytes. Overlapping DEPs significantly upregulated in VEXAS $CD14^+$ cells, THP1 model cells, and 32 model cells that were also reported to be ERAD substrates are highlighted in blue. **(e)** Quantification of total STING levels by DIA-based mass spectrometry (*left graph*) and ubiquitylated STING at K338 by diGly proteomics (*left graph*), showing that STING progressively accumulate while ubiquitylation of STING at K338 decreases upon loss of cytoplasmic ubiquitylation in *$UBA1^{M41V}$* THP1 monocytes. n = 3 biological replicates, error bar = s.e.m, * = $p < 0.05$, student's t-test. **(f)** Heatmap representation of relative protein levels upon loss of cytoplasmic ubiquitylation in THP1 $UBA1^{M41}$ monocytes across indicated time periods after doxycycline induction, revealing that STING levels increases associate with increases in inflammatory cytokines and proteins. Representative proteins were selected from those significantly dysregulated in patient $CD14^+$ cell proteomics. Color scale indicates the $\log_2$-fold change of $UBA1^{M41V}$ versus $UBA1^{WT}$ THP1 monocytes.

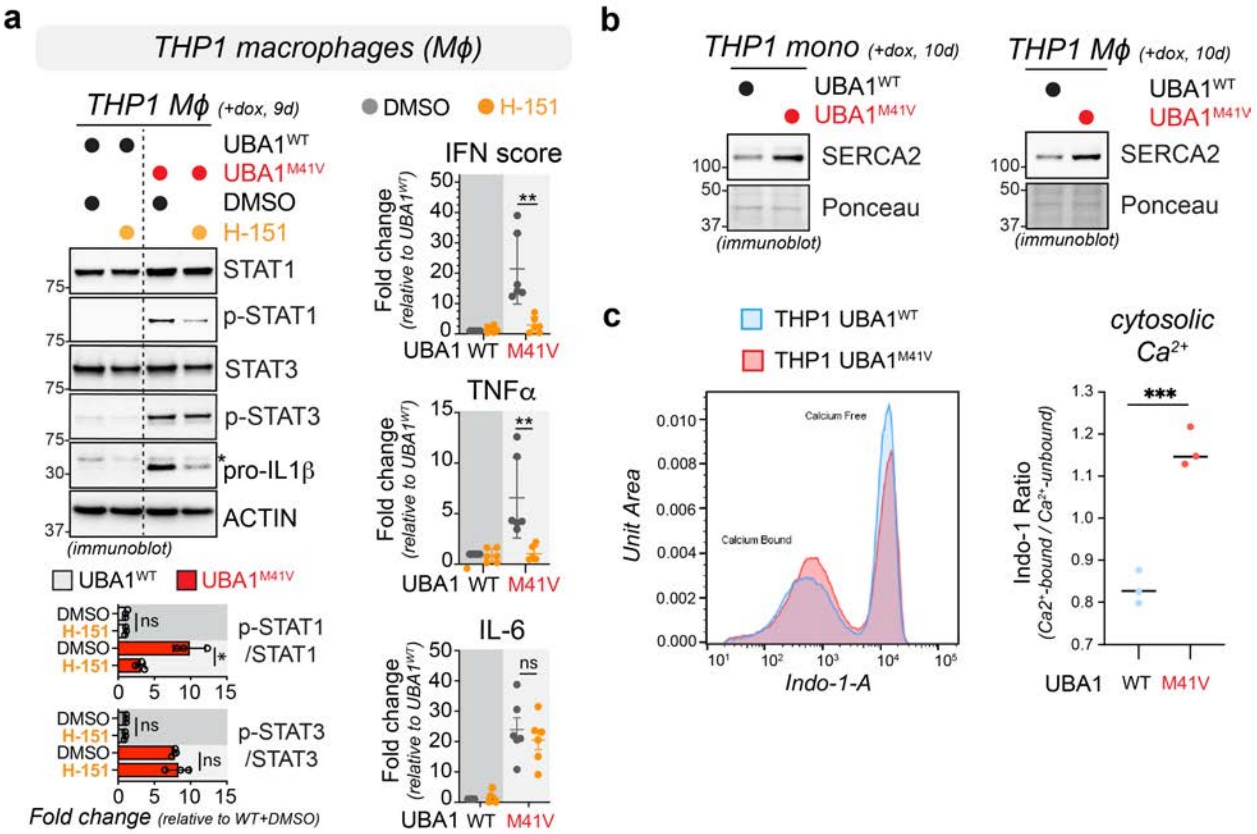

**Supplementary Figure 15. Pharmacological STING inhibition reverses most axis of inflammation in THP1 VEXAS macrophages and THP1 VEXAS model cells exhibit dysregulated calcium homeostasis. (a)** STING inhibition reverses most VEXAS-associated inflammatory signaling in THP1 model cells. Left panel shows immunoblot analysis of $UBA1^{WT}$ and $UBA1^{M41V}$ THP1 macrophages (Mf) treated with STING inhibitor H-151 (20 µM) for 24 h using indicated antibodies. Graphs below depict quantifications of p-STAT1/STAT1 and p-STAT3/STAT3 ratio normalized to DMSO-treated $UBA1^{WT}$ THP1 macrophages. n = 3 biological replicates, error bar = s.e.m, * = p < 0.05 ** = p < 0.01, student's t-test. Right panel shows qPCR analysis of these samples to determine mRNA levels of type I interferon stimulated genes (IFN score), TNFα, and IL-6. Samples were normalized to DMSO-treated $UBA1^{WT}$ THP1 macrophages. n = 3 biological replicates with 2 technical replicates each, error bar = s.e.m, * = p < 0.05 ** = p < 0.01, student's t-test. **(b)** Loss of cytoplasmic UBA1 function in THP1 model cells leads to increased levels of the ER-resident calcium pump SERCA2. $UBA1^{WT}$ and $UBA1^{M41V}$ THP1 monocytes (*left panel*) or macrophages (Mf, *right panel*) were treated with dox for 10 d and subjected to immunoblotting using the indicated antibodies. Results are representative of n = 3 biological replicates. **(c)** Loss of cytoplasmic UBA1 function in THP1 monocytes leads to increased cytosolic calcium levels. $UBA1^{WT}$ and $UBA1^{M41V}$ THP1 monocytes were treated with dox for 10 d, stained with 3 µM Indo-1, and subjected to flow cytometry (*left panel*) followed by calculation of the bound calcium to unbound calcium ratio (*right panel*). n = 3 biological replicates, *** = p < 0.001, student's t-test.

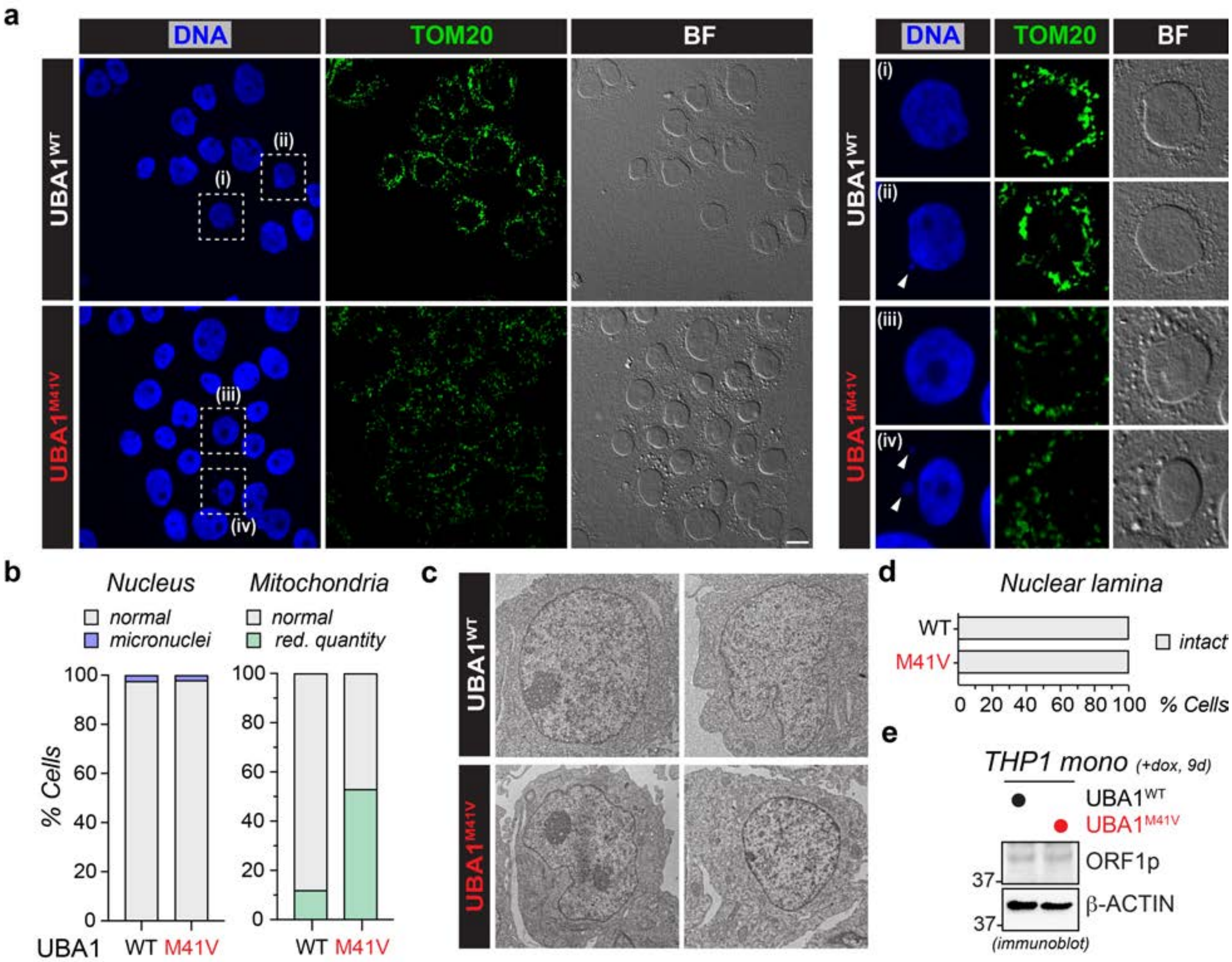


**Supplementary Figure 16. *UBA1$^{M41V}$* THP1 monocytes exhibit mitochondrial but not nuclear abnormalities. (a)** The quantity of mitochondria in vacuole-containing VEXAS THP1 cells are reduced while the number of micronuclei is not changed, as demonstrated by brightfield (BF), single-plane confocal anti-TOM20 immunofluorescence, and DAPI staining images of *UBA1$^{WT}$* and *UBA1$^{M41V}$* THP1 cells treated with doxycycline for 10 days. Representative images of 2 biological replicates are shown. White arrows highlight micronuclei. **(b)** Quantification of cells containing micronuclei (*left graph*) or abnormal amount of mitochondria (*right graph*). n £ 800 cells from 2 biological replicates. **(c)** THP1 *UBA1$^{M41V}$* monocytes exhibit an intact nuclear lamina. *UBA1$^{WT}$* and *UBA1$^{M41V}$* THP1 monocytes induced with doxycycline for 10 days and imaged by transmission electron microscopy (TEM). This is the same experiment depicted in Fig S10 but analyzed for nuclear integrity. **(d)** Quantification of nuclear lamina integrity of *UBA1$^{WT}$* and *UBA1$^{M41V}$* THP1 cells. n > 20 cells per condition. **(e)** Loss of cytoplasmic UBA1 function in THP1 model cells does not lead activate LINE-1 retrotransposon elements, as indicated by anti-ORF1p immunoblotting. UBA1$^{WT}$ and UBA1$^{M41V}$ THP1 monocytes were treated with dox for 9 d and subjected to immunoblotting using the indicated antibodies.

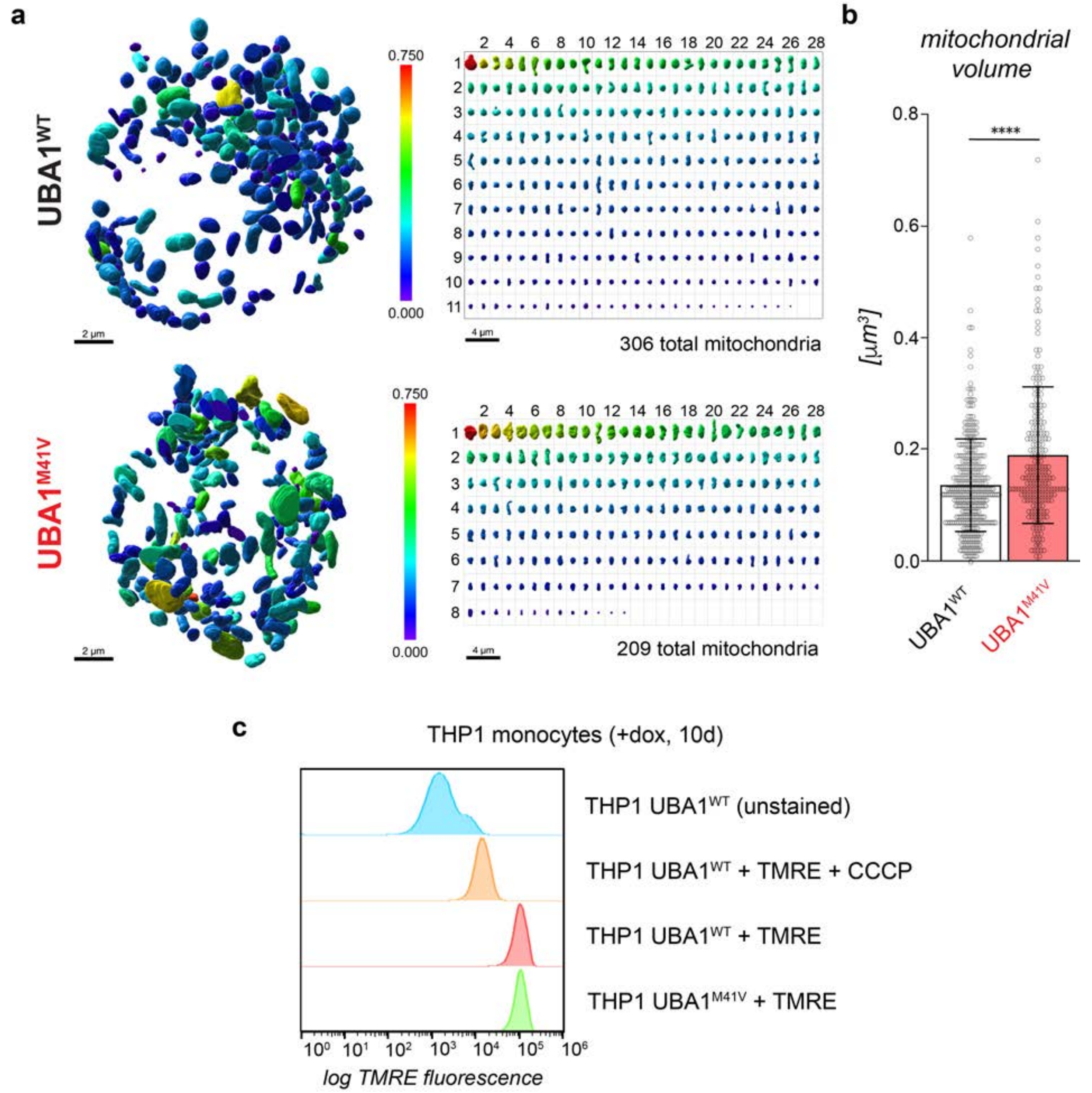


**Supplementary Figure 17. *UBA1$^{M41V}$* monocytes contain less mitochondria that are morphologically abnormal but exhibit no difference in membrane potential. (a)** Analysis of 3D-reconstructed mitochondria from FIB-SEM volumes reveal that *UBA1$^{M41V}$* THP1 monocytes have less mitochondria that are on average larger than those found in *UBA1$^{WT}$* cells, indicative of abnormal morphology. One representative *UBA1$^{WT}$* and *UBA1$^{M41V}$* THP1 monocyte cell was subjected to FIB-SEM followed by mitochondria segmentation and volume determination. See also Movie 2. **(b)** Quantification of volumes of segmented mitochondria shown in panel a, revealing a significant increase in mitochondrial volume in *UBA1$^{M41V}$* THP1 monocytes. **(c)** Representative flow cytometry experiment of THP1 VEXAS model cells, showing that the mitochondrial membrane potential of THP1 UBA1$^{WT}$ and UBA1$^{M41V}$ does not differ. THP1 cells were treated with dox for 10 d, subjected to 200 µM CCCP treatment as indicated, and incubated with 50 nM TMRE, followed by flow cytometry analysis with 100,000 events per condition. Of note, the THP1 VEXAS model cells were generated using mCherry as selection marker and thus the control (no TMRE) exhibits high background fluorescence.

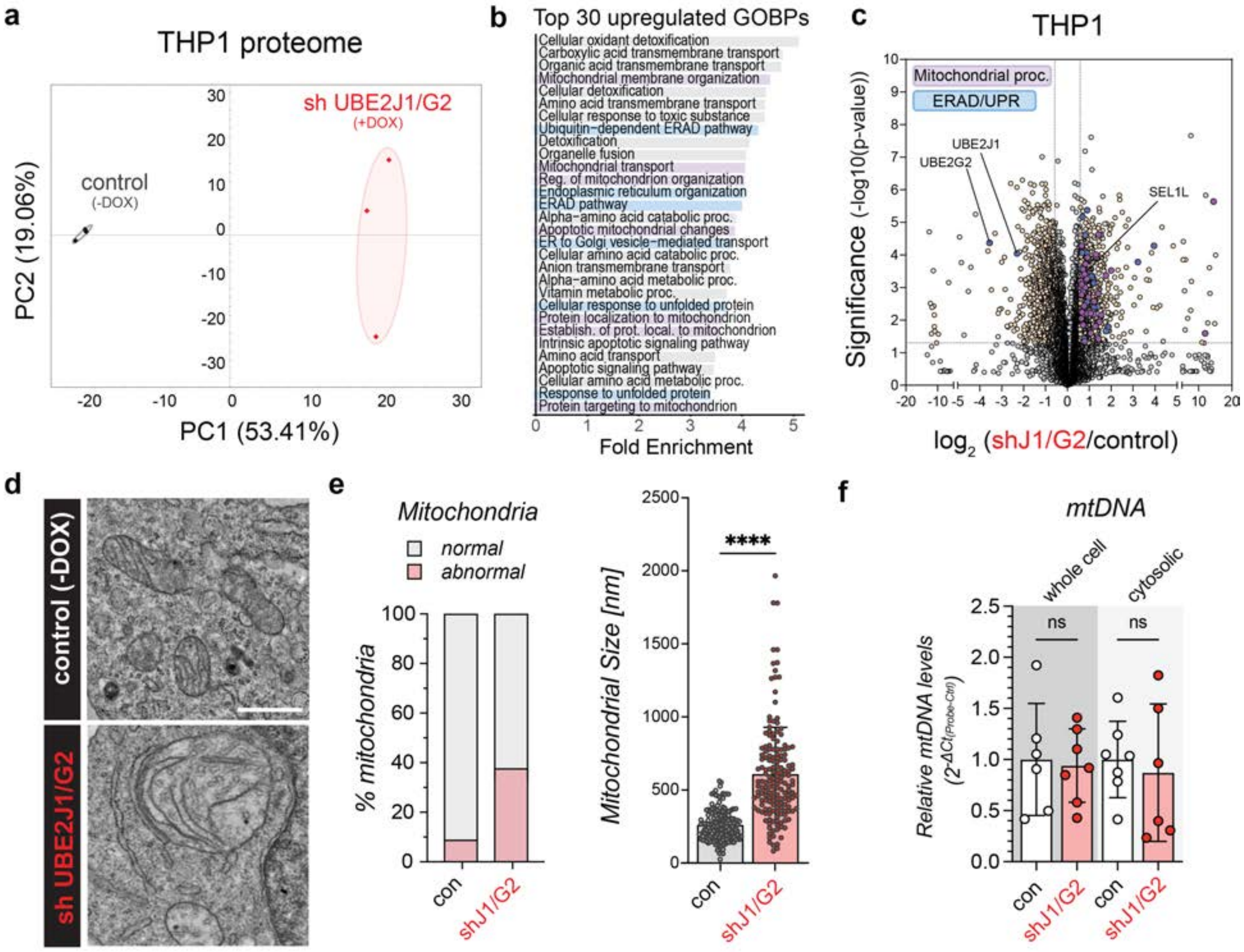


**Supplementary Figure 18. ERAD inhibition in THP1 monocytes causes mitochondrial pathway dysregulation and morphological abnormalities but no cytosolic mtDNA leakage.** **(a)** UBE2J1/G2 depletion in THP1 monocytes causes differences in the proteome. THP1 monocytes were treated with doxycycline for 6 days to induce shRNA-mediated UBE2J1/G2 knockdown and total proteomes were determined by DIA mass spectrometry followed by principal component (PC) analysis. **(b)** UBE2J1/G2-depleted THP1 cells exhibit dysregulated ERAD and mitochondrial pathways. Control or UBE2J1/G2 depleted THP1 cells were subjected to DIA mass spectrometry followed by GOBP analysis of significantly differentially upregulated proteins ($p<0.05$, $log_2$(UBE2J1/G2/control) > 0.58). The top 30 significantly enriched GOBPs (FDR < 0.05) ranked according to fold enrichment score are shown. GOBPs related to ERAD (blue) and mitochondrial processes (purple) are highlighted. **(c)** UBE2J1/G2-depleted THP1 cells exhibit dysregulated ERAD and mitochondrial pathways. The volcano plots depict differentially expressed proteins (DEPs) of control and UBE2J1/G2-depleted THP1 monocytes 6 days post dox induction. DEPs ($p<0.05$, $log_2$(UBE2J1/G2/control) > 0.58) related to ERAD and mitochondrial processes are highlighted. **(d)** UBE2J1/G2-depleted THP1 monocytes exhibit mitochondrial morphology abnormalities, indicative of impaired function. Control or UBE2J1/G2-depleted THP1 cells were treated with dox for 6 days and analyzed by transmission electron microscopy. Scale bar = 1 $\mu$m. **(e)** UBE2J1/G2-depleted THP1 monocytes exhibit a higher percentage of abnormal mitochondria that are larger in size as compared to control cells. The left graph depicts the quantification of mitochondrial morphology on TEM images of the experiment shown in panel d. n > 10 cells. The right graph depicts the quantification of the mitochondrial sizes on TEM images of the experiment shown in panel d. n > 10 cells, error bar = s.d., **** = $p < 0.0001$, student's t-test. **(f)** UBE2J1/G2-depleted THP1 cells exhibit no significant changes in cytoplasmic mitochondrial DNA levels, as revealed by qPCR analysis of whole cell or cytoplasmic fractions of control and UBE2J1/G2-depleted THP1 monocytes. n = 2 biological replicates with 2-3 technical replicates for mitochondrial DNA probe *MTDN1*, error bar = s.d., ns = not significant , student's t-test.

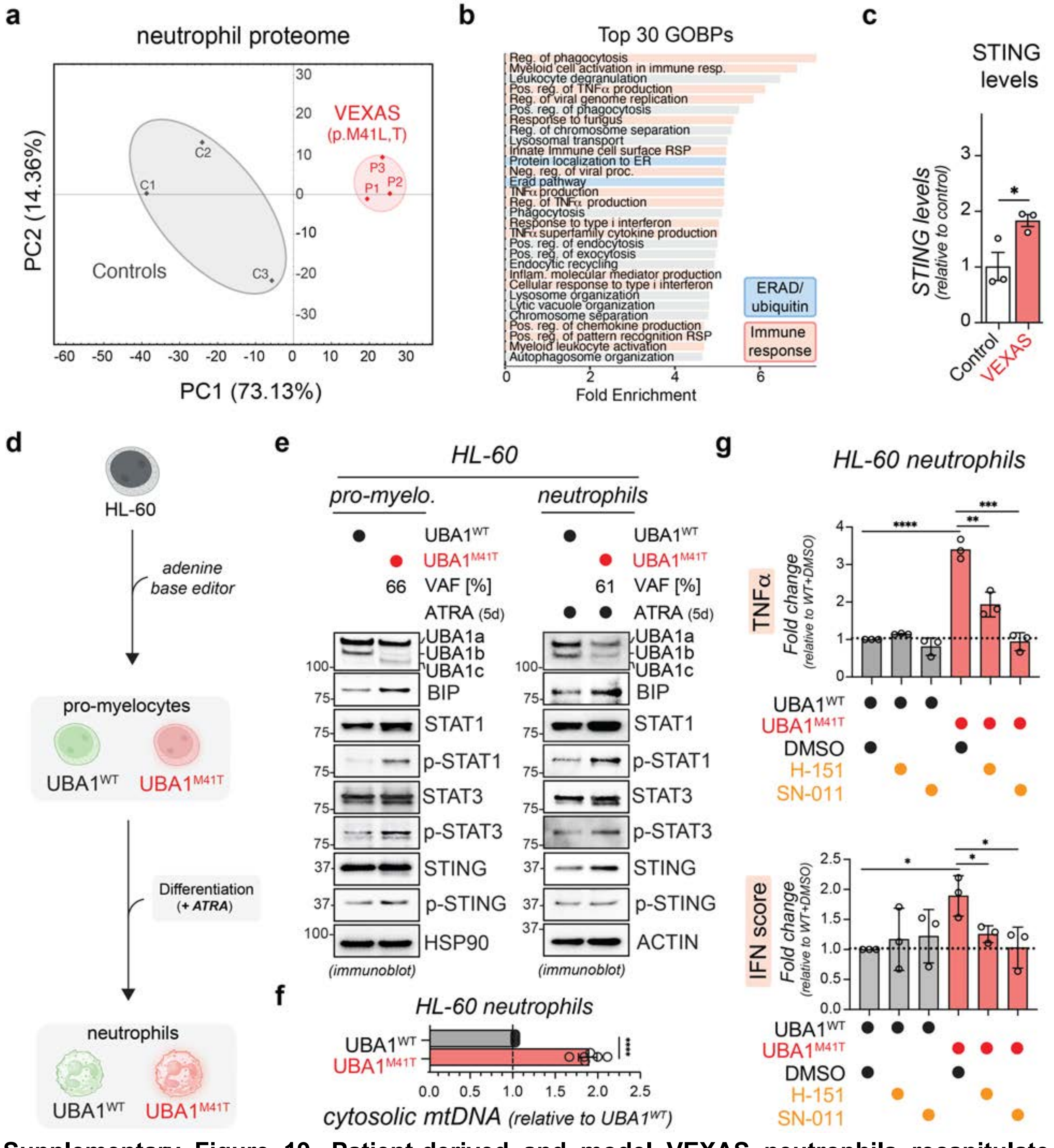


**Supplementary Figure 19. Patient-derived and model VEXAS neutrophils recapitulate ERAD dysregulation and STING-dependent inflammation observed in VEXAS monocytes and macrophages. (a)** The proteomes of VEXAS patient derived neutrophils are different from those of healthy donor controls. Neutrophils were isolated from VEXAS patients or healthy donors and total proteomes were determined by DIA proteomics followed by principal component (PC) analysis. **(b)** Neutrophils from VEXAS patients exhibit dysregulated ERAD and immune responses, as revealed by GOBP analysis of significantly DEPs (p<0.05, -0.7 <

$\log_2$(VEXAS/control) > 0.7). Top 30 significantly enriched GOBPs (FDR < 0.05) ranked according to fold-enrichment score are shown. **(c)** Quantification of total STING levels by DIA-based mass spectrometry, showing that STING levels are significantly higher in VEXAS neutrophils. n = 3 biological replicates, error bar = s.e.m, * = p < 0.05, student's t-test. **(d)** Schematic of base editing approach to generate *UBA1$^{M41T}$* HL-60 promyelocytes or neutrophils. **(e)** *UBA1$^{M41T}$* HL-60 promyelocytes or neutrophils exhibit UPR activation, elevated inflammatory signaling, and STING dysregulation, similar to VEXAS monocytes and macrophages. HL-60 cells were subjected to base editing, cultured for 4 days and differentiated with 1 µM of ATRA for 5 days as indicated, followed by immunoblot analysis with indicated antibodies. **(f)** *UBA1$^{M41T}$* HL-60 neutrophils exhibit elevated levels of cytoplasmic mitochondrial DNA, as revealed by qPCR analysis of cytoplasmic fractions of *UBA1$^{WT}$* and *UBA1$^{M41T}$* cells. n = 3 biological replicates with 2 technical replicates for two different mitochondrial DNA probes (see methods), error bar = s.e.m, **** = p < 0.0001, student's t-test. **(g)** STING inhibition reverses VEXAS-associated inflammatory signaling in HL-60 VEXAS model cells. UBA1$^{WT}$ and UBA1$^{M41T}$ HL-60 neutrophils were treated with STING inhibitors H-151 (1 µM) or SN-011 (10 µM) for 24 h and subjected to qPCR analysis to determine mRNA levels of or TNFα (*upper graph)* or different type I interferon stimulated genes (*lower graph*, IFN score) or TNFα (*lower graph).* n = 3 biological replicates, error bar = s.e.m, * = p < 0.05, ** = p < 0.01, *** = p < 0.001, **** = p < 0.0001, one-way ANOVA.

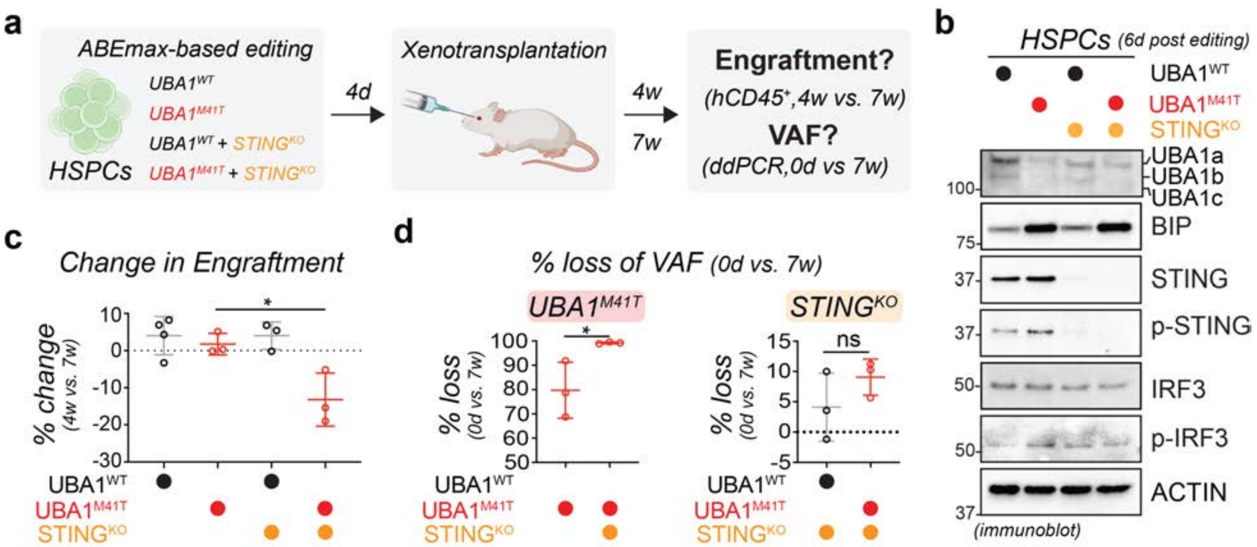


**Supplementary Figure 20. STING inhibition reduces engraftment of VEXAS HSPCs in mice.** **(a)** Experimental scheme for adenine base editing (ABE) of human hematopoietic stem and progenitor cells (HSPCs) to generate the indicated genotypes followed by xenotransplantation and assessment of engraftment and mutation burden or variant allele fraction (VAF). **(b)** Immunoblot analysis of HSPCs after 6 d post editing with indicated antibodies, validating successful base editing and showing that UBA1$^{M41T}$-edited HSPCs exhibit abnormal activation of STING signaling, which is reversed by simultaneously editing the *STING* locus for STING knock out (STING$^{KO}$). **(c)** Loss of STING decreases engraftment of *UBA1$^{M41T}$* but not *UBA1$^{WT}$* HSPCs in xenotransplantation. HSPCs base-edited as depicted in panel e were engrafted into mice, hCD45$^{+}$ cells isolated after 4 and 7 weeks of engraftment, and the loss percentage of engraftment was calculated for each genotype. n = 3 biological replicates, error bar = s.e.m, * = $p < 0.05$, one-way ANOVA. **(d)** Loss of STING decreases the mutational burden of UBA1$^{M41T}$ HSPCs in xenotransplantation. Variant allele fraction for *UBA1$^{M41T}$* mutations are lost upon co-editing of STING; however STING knockout is not impacted by *UBA1$^{M41T}$* mutations, as shown by digital droplet PCR (ddPCR) analysis of HSPCs before (0 d) and 7 weeks after engraftment. n = 3 biological replicates, error bar = s.e.m, * = $p < 0.05$, student's t-test.

**Supplementary Table 1:** Combined table of GSEA output for proteomic and transcriptomics datasets for all GOBOP gene sets, conveying enrichment of each gene set in sample group of interest.

**Supplementary Table 2:** Table of significantly differentially expressed proteins, with their pvalue and log2FoldChange values calculated from differential analysis. Values are considered in CD14 + VEXAS patient cells (UBA1 p.Met41V/T/L) relative to healthy donor CD14+ cells (UBA1 WT).

**Supplementary Table 3:** Table of significantly differentially expressed proteins HGNC ids, with their pvalue and log2FoldChange values calculated from differential analysis. Enrichment is considered in UBA1 p.Met41V THP-1 monocytes relative to UBA1 WT THP-1 monocytes.

**Supplementary Table 4**: Table of significantly differentially expressed gene HGNC ids, with their pvalue and log2FoldChange values calculated from differential analysis. Values are considered in CD14 + VEXAS patient cells (UBA1 p.Met41V/T/L) relative to healthy donor CD14+ cells (UBA1 WT).

**Supplementary Table 5**: Table of significantly differentially expressed gene HGNC ids, with their pvalue and log2FoldChange values calculated from differential analysis. Enrichment is considered in UBA1 p.Met41V THP-1 monocytes relative to UBA1 WT THP-1 monocytes.

**Supplementary Table 6:** Table of known ERAD substrates.

**Supplementary Table 7:** Combined table of GSEA output for each dataset for all GOBOP gene sets in STING inhibition experiments, conveying enrichment of each gene set in sample group of interest.

**Supplementary Table 8:** Table of significantly differentially expressed gene HGNC ids, with their pvalue and log2FoldChange values calculated from differential analysis. Values are considered in DMSO treated VEXAS patient cells (UBA1 p.Met41V/T/L) relative to DMSO treated healthy donor cells (UBA1 WT).

**Supplementary Table 9:** Table of significantly differentially expressed gene HGNC ids, with their pvalue and log2FoldChange values calculated from differential analysis. Values are considered in STING inhibited (H151 treated) VEXAS patient cells (UBA1 p.Met41V/T/L) relative to STING inhibited (H151 treated) healthy donor cells (UBA1 WT).

**Supplementary Table 10:** Demographic, clinical characteristics, treatment, and *UBA1* genetic testing of VEXAS patients who participated in this study.

**Supplementary Movie 1: THP1 *UBA1$^{M41V}$* monocytes exhibit dilated ER network.** A subset of 100 slices of grayscale FIB-SEM volume from *UBA1$^{WT}$and UBA1$^{M41V}$* cells overlayed with ER segmentation mask pseudocolored in red. The side-by-side z-slice traversal reveals morphological differences between the *UBA1$^{WT}$* and the *UBA1$^{M41V}$* ER network. Scale bar 500 nm.

**Supplementary Movie 2: *UBA1$^{M41V}$* THP1 monocytes exhibit abnormal mitochondria morphology**. A subset of 300 slices from each of the FIB-SEM volumes shows side-by-side comparison of a representative region of cytoplasm from *UBA1$^{WT}$ and UBA1$^{M41V}$* cells. Mitochondria are pseudo-colored in red. Note large, dilated cristae most often seen in *UBA1$^{M41V}$* cell as well as dilated ER (not pseudo-colored) across the cytoplasm. Scale bar 1 um.
